# Learning Universal Representations of Intermolecular Interactions with ATOMICA

**DOI:** 10.1101/2025.04.02.646906

**Authors:** Ada Fang, Michael Desgagné, Zaixi Zhang, Andrew Zhou, Joseph Loscalzo, Bradley L. Pentelute, Marinka Zitnik

## Abstract

Molecular interactions underlie nearly all biological processes, yet most representation models describe isolated entities or specialize in a single molecular setting. Here, we introduce ATOMICA, an interaction-centered geometric deep learning model designed to learn transferable representations of intermolecular interfaces across proteins, small molecules, metal ions, and nucleic acids. Self-supervised pretraining on 2,037,972 interaction complexes yields representations spanning atoms, molecular building blocks, and complete interfaces. The latent space captures molecular identity and interaction context, supporting sequence recovery and zero-shot prioritization of residues involved in non-covalent interactions. ATOMICA provides structural information complementary to sequence representations on RNA and protein-pocket ligand classification. Across protein-pocket analyses, ATOMICA distinguishes ATP- and ADP-associated pocket states and retrieves ligand-matched pockets across proteins without detectable structural alignment. The latent space also enables cross-modal comparison, with orthosteric inhibitor embeddings retrieving regions proximal to native peptide and protein interfaces. Applied to the dark proteome, ATOMICA-Ligand predicts candidate ions or cofactors for 2,646 pockets, and five heme candidates show Soret-band shifts consistent with heme association. Together, these results show how interaction-centered molecular representations can transfer structural information across molecular interaction types and generate experimentally testable hypotheses.

## Main

Molecular interactions underpin chemical and biological processes, from enzymatic catalysis and signaling to gene regulation and drug binding. Recent multimolecular models, including AlphaFold3 [1] and RoseTTAFold All-Atom [2], show that a single model can capture structures across diverse classes of interacting biomolecules. These advances point to shared principles underlying biomolecular interactions. Many problems extend beyond reconstructing the structure of complexes and instead require transferable representations of intermolecular interfaces that can be used for comparison, annotation, and prediction across molecular modalities. Since intermolecular recognition is governed by local geometry and chemistry, this motivates representation learning directly on interaction interfaces.

In parallel, unimolecular foundation models, including protein language models [3–7], nucleicacid language models [8–11], and small-molecule encoders [12–14], learn strong within-modality priors from large-scale single-entity datasets. These models typically do not incorporate paired interacting partners as a primary pretraining signal and therefore do not necessarily align interaction features across molecular modalities. Complementing these efforts, a large body of work learns interaction representations for specific modality pairs, such as protein–protein [15–21], protein–ligand [22–25], protein–RNA [26–31], protein–ion [32–34], and RNA–ligand [35–37]. Because these approaches are typically built around particular molecular types, featurizations, and datasets, extending them to additional modalities often requires new encoders, new supervision, or re-specifying the interaction representation. These limitations motivate an interaction-centered representation model grounded in 3D interface geometry and local chemistry, since non-covalent interactions are governed by the spatial arrangement of atoms [38]. Interface geometry varies across molecular modalities [39–41], and a shared model can provide a common representation across interface types, allowing features learned in one setting to transfer to others.

We introduce ATOMICA, an all-atom geometric deep learning model that learns representations of intermolecular complexes (Fig. 1a) across four molecular modalities: proteins, small molecules, metal ions, and nucleic acids, encompassing eight interaction types. ATOMICA is trained on a dataset of 2,037,972 interaction complexes (Fig. 1b) from the Cambridge Structural Database (CSD) [42] and the Protein Data Bank (PDB) [43]. The model uses a geometric, SE(3)-equivariant architecture and is trained with a self-supervised objective that combines denoising of atomic coordinates and masked block-type prediction. ATOMICA learns representations directly from interaction interfaces at the scales of atoms, molecular building blocks, and complete interfaces. Across the evaluated interaction types, multimodal pretraining improves masked block-type prediction, particularly for protein–DNA and protein–RNA interfaces. These representations also support sequence recovery and enable zero-shot ATOMICASCORE to prioritize residues involved in annotated non-covalent interactions. Representations of ions also retain the local coordination geometry.

**Figure 1.**
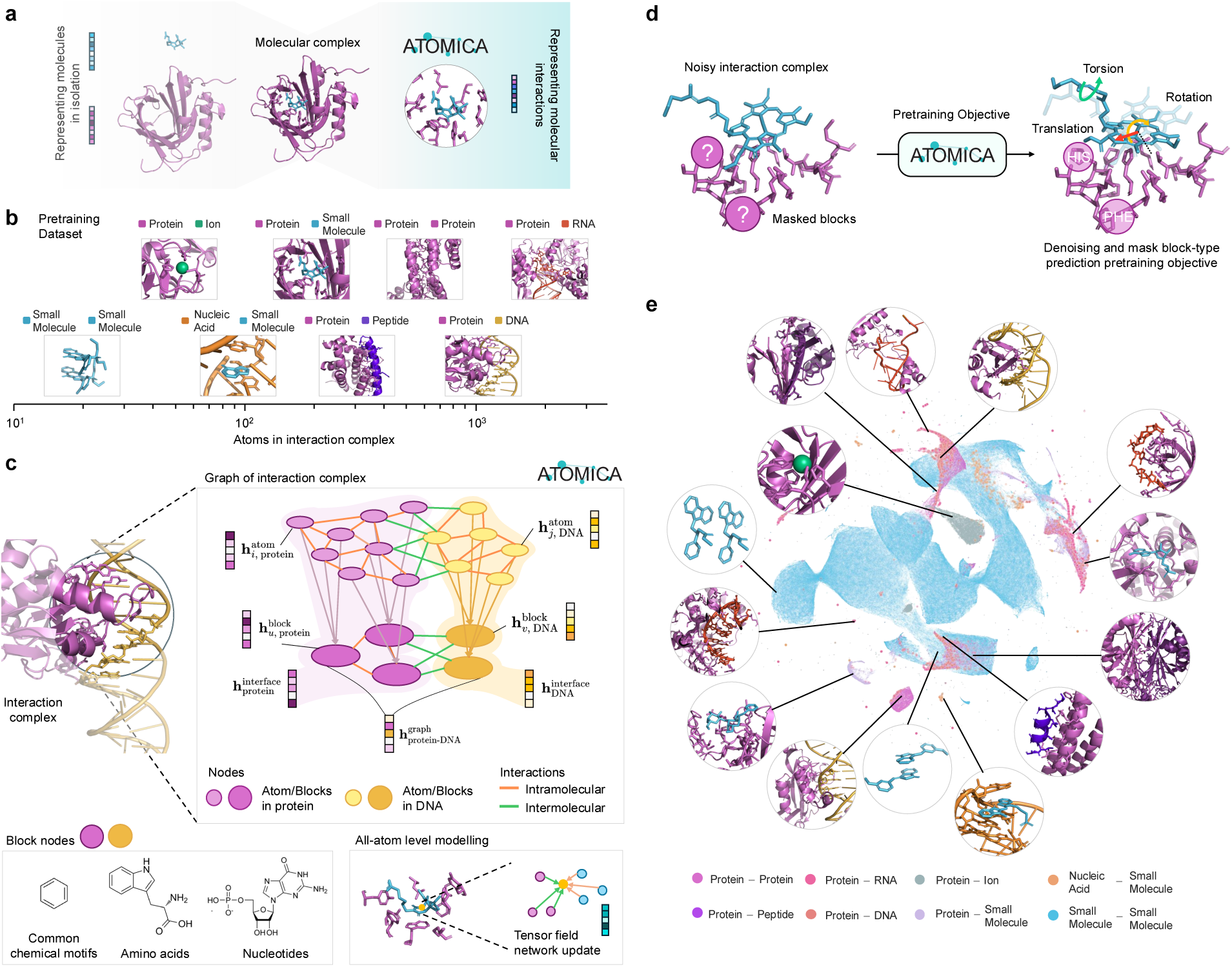
Overview of ATOMICA pretraining data, architecture, and latent space. **a** Complementary to models learning representations of individual molecules, ATOMICA focuses on representations of intermolecular interfaces within complexes, providing a shared interaction-centered representation for interaction tasks. **b** Input modalities and the number of atoms in the interaction complexes used to pretrain ATOMICA. **c** Overview of the ATOMICA architecture. Interaction complexes are modeled at the atom and block levels. Message passing between nodes at each level is done via intermolecular and intramolecular edges. All embeddings that can be derived from ATOMICA are also shown. **d** Self-supervised pretraining objectives for generating ATOMICA embeddings. **e** UMAP projection of the latent space of all interaction complexes seen during pretraining.

These interface representations transfer to RNA and protein-pocket prediction, where they contribute structural information complementary to sequence models. ATOMICA achieves the highest AUPRC among evaluated models for RNA–protein binding-site prediction and complements sequence-based models on pocket ligand classification. In analyses of local pocket geometry, it distinguishes ligand-free ATP- and ADP-associated pocket states and retrieves ligand-matched pockets across proteins without detectable structural alignment. The ATOMICA latent space further enables direct comparison of interfaces formed by chemically different partners. In orthosteric inhibitor systems, small-molecule inhibitor embeddings retrieve regions proximal to native peptide and protein interfaces across the evaluated superfamilies. This shows how the latent space relates chemically distinct partners of a common target interface.

Finally, we fine-tune ATOMICA on labeled pocket–ligand complexes to predict ligand identity from local pocket geometry, yielding ATOMICA-Ligand, which assigns putative ions and cofactors to 2,646 pockets in Foldseek-defined dark clusters with limited functional annotation [44–46]. Five heme candidates show Soret-band shifts consistent with heme association. These results illustrate how ATOMICA can generate experimentally testable molecular hypotheses.

## Results

### Dataset of protein, small molecule, ion, and nucleic acid interactions

We assembled a dataset of interacting molecular entities from the Cambridge Structural Database (CSD) [42] and Q-BioLiP [47, 48], which contains intermolecular interactions across all modalities available in the Protein Data Bank (PDB) [49]. From the CSD, we filtered for entries of small-molecule crystals and extracted unique pairs of neighboring conformers from their packed unit cells (Methods 1), resulting in 1,767,710 interacting pairs of small molecules, of which 1,747,710 are used for training. These pairs represent contacts in packed crystal lattices, distinct from curated biological complexes, and are not assigned biological function. They nevertheless provide diverse, high-resolution examples of non-covalent contact geometries, including hydrogen-bond, halogen-bond, *π*-stacking, and van der Waals arrangements, across a broader range of common chemical motifs than is available in the PDB. We use these contacts as selfsupervised pretraining examples, without treating them as labels of biological binding. We also processed 337,993 interaction complexes from Q-BioLiP, of which 290,262 are used for training. This includes structures of protein complexes with proteins, DNA, RNA, peptides, small molecules, and metal ions, as well as nucleic acid ligand structures from the PDB. Within the training dataset, CSD-derived data account for 85.8% of complexes, 58.6% of blocks, and 44.3% of atoms, whereas PDB-derived data account for 14.2%, 41.4%, and 55.7%, respectively (Methods 3). The interaction interface between two entities is defined by the atoms within 8 Å of the other molecule to capture atoms in their surrounding molecular context. In Fig. 1b, we show examples of the interaction interface for various complexes and their relative size.

We represent interaction complexes using two-level graphs to capture atomic-level details and higher-order chemical structures (Fig. 1c). This hierarchical representation has theoretically greater expressive power than atom-level graphs alone [50]. At the first level, nodes in the graph represent atoms, each defined by its element type and 3D spatial coordinates. At the second level, atoms are grouped into blocks, the model’s shared representation of an amino-acid residue, a nucleotide, a common chemical motif, or a single atom, to form a block-level graph [51–53] (Methods 1.3). Within each graph level, we define two types of edges: intramolecular and intermolecular edges, which connect the *k* nearest nodes in Euclidean space within and on the interface of interacting molecules, respectively.

### Geometric representation learning across intermolecular modalities

ATOMICA is a self-supervised geometric graph neural network that generates multiscale embeddings at the atom, block, and graph levels from the structure of two interacting molecules (refer to Fig. 1c for the embedding notation used throughout). ATOMICA can be fine-tuned and adapted with task-specific heads for predictive tasks such as binding-site annotation. Embeddings are learned with SE(3)-equivariant tensor field networks that pass messages over intra- and intermolecular edges (Methods 2), a framework used for interatomic potentials, docking, and RNA structure scoring [54–57].

To train ATOMICA, we use self-supervised pretraining objectives that combine denoising and masked block-type prediction. Denoising is an effective pretraining strategy for learning molecular representations [24, 58–61]. In ATOMICA, denoising requires the model to reconstruct an interaction complex graph after it is perturbed by a rigid SE(3) transformation and random torsion angle rotations of one molecular entity at the interface (Fig. 1d). SE(3) equivariance enforces the appropriate transformation of model features under global rotations and translations, while denoising trains the model to represent relative geometry between molecular partners and local conformation at the interface. Inspired by masked modeling objectives used to learn protein [5] and nucleic-acid [11] representations, ATOMICA is also trained to predict the type of masked blocks at the interaction interface. Together, these objectives encourage the model to learn geometric and chemical features that are relevant to intermolecular interfaces.

### Organization of ATOMICA embeddings across scales

A two-dimensional UMAP projection [62] of ATOMICA graph-level embeddings shows broad organization across interaction types (Fig. 1e). We expect the protein–protein and protein– peptide graph embeddings to be relatively close in the latent space as they share similar intermolecular interactions. Consistent with this expectation, 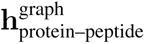 embeddings have greater cosine similarity to 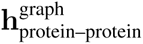 graph embeddings than to graph embeddings from other interaction types (KS *D* = 0.100). This provides a quantitative check that the graph-level ATOMICA embeddings retain the expected similarity between these related interaction classes. In the first two principal components of ATOMICA mean atom-level embeddings, elements show partial separation by elemental category, including partial separation of metals and nonmetals (Fig. 2a). In the first two principal components of ATOMICA mean block-level embeddings, amino-acid blocks and nucleotide blocks (DNA and RNA) occupy partially distinct regions (Fig. 2b). Together, these analyses evaluate how ATOMICA organizes its latent space at the graph, block, and atom levels.

**Figure 2.**
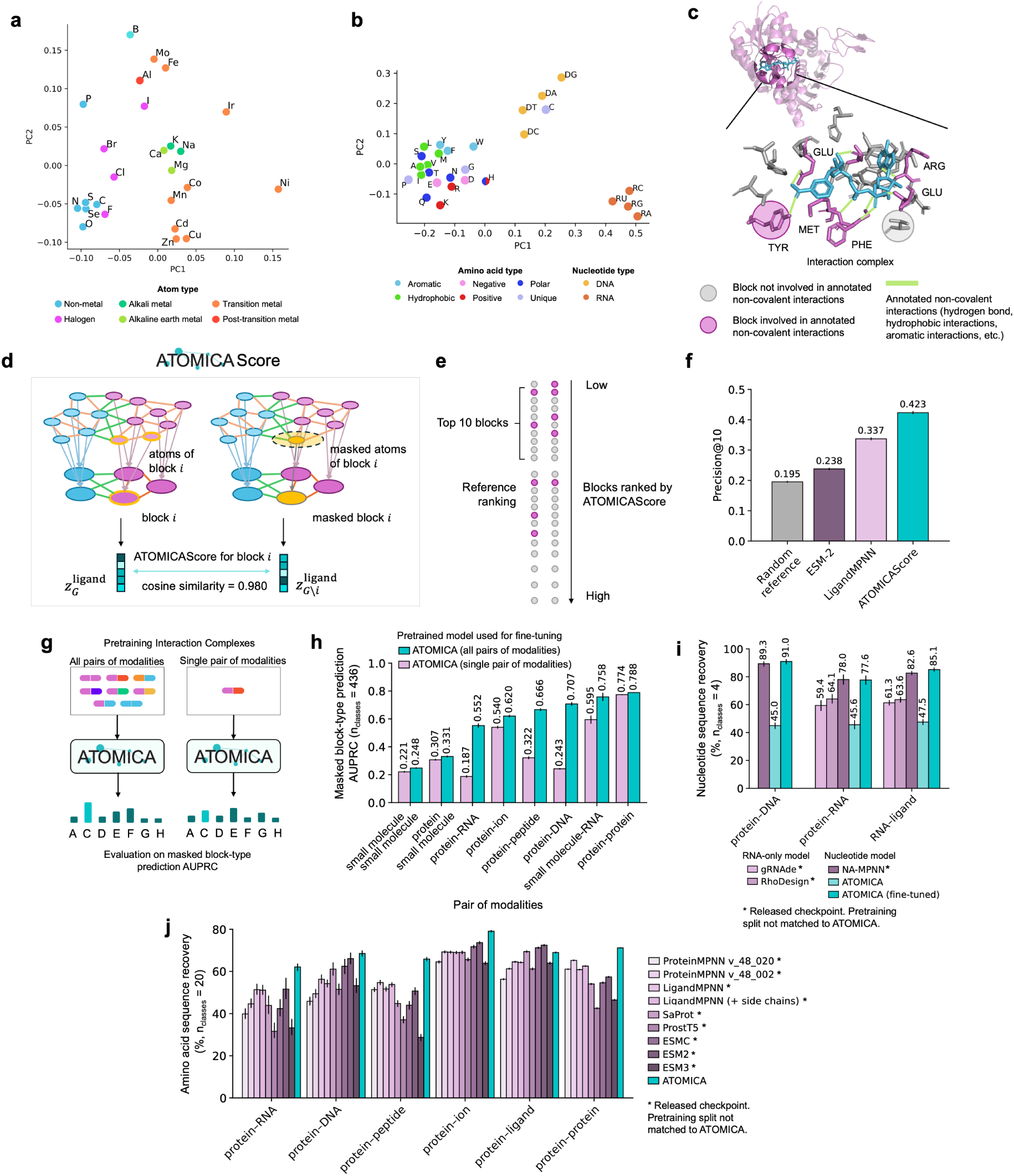
Characterization of learned representations in ATOMICA. **a** Principal component analysis of mean learned atom embeddings by element. **b** Principal component analysis of mean learned block embeddings by amino-acid and nucleotide identity. **c** Example of residues involved in annotated non-covalent interactions at a protein–small-molecule interface. **d** Definition of ATOMICASCORE by masking residue *i* and comparing the interaction representation before and after masking. Lower scores indicate greater importance. **e** Ranking of interface residues by ATOMICASCORE to identify residues involved in annotated non-covalent interactions. **f** Precision@10 for residues involved in annotated non-covalent interactions on held-out complexes, comparing ATOMICASCORE with LigandMPNN, ESM-2, and a random reference. **g** Schematic comparison of multimodal and modality-specific pretraining. **h** Masked block-type prediction AUPRC for multimodal and modality-specific models. **i** Nucleotide sequence recovery at 10% interface masking, reported for ATOMICA (fine-tuned). **j** Amino-acid sequence recovery at 10% interface masking, reported for the pretrained ATOMICA model without task-specific fine-tuning. Error bars denote 95% bootstrap confidence intervals over pockets in **f** or complexes in **h–j**.

Cysteine is the one exception to this separation because it is the only amino acid whose mean block embedding is closer to the deoxyribonucleotides than to the other amino acids in the projection. However, cysteine does not sit inside the nucleotide cluster in the full ATOMICA embedding space. Because block embeddings are contextual, this pattern does not necessarily indicate a chemical resemblance to nucleotides, but is consistent with the interaction environments in which cysteine occurs, including its threefold enrichment in protein–ion complexes relative to the average amino acid.

Different molecular interfaces involve distinct local interaction geometries. We therefore tested whether ATOMICA’s shared interface architecture retains the directional coordination information characteristic of metal-binding sites. We fitted linear probes to frozen 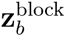 for each metal-ion block *b* in the held-out protein–ion split, where 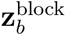 is the block-level representation that retains rotation-invariant information from the equivariant features (Methods 9 and Supplementary Table S1). We use coordination-number labels from MetalPDB [63] and coordination-geometry classes from FindGeo [64] for our protein–ion held-out test set. For geometry classification, an element-only probe reached a balanced accuracy of 0.171, while frozen ATOMICA reached 0.340. Among the 635 sites with a deposited coordination number of six, chance and the element-only probe both reached 0.333 across the three observed geometry classes, while frozen ATOMICA reached 0.499. Across all 3,654 labeled sites, ATOMICA predicted the full MetalPDB coordination number over all deposited donors with a balanced accuracy of 0.368 and the coordination number restricted to protein donors visible in the model input with a balanced accuracy of 0.578.

### ATOMICASCORE highlights key residues at molecular interfaces

Here we distinguish between the *interface region* (all residues within a certain distance neighborhood of the partner) and residues involved in *annotated non-covalent interactions*, such as hydrogen bonds, hydrophobic interactions, and aromatic interactions, identified by PLIP using established geometric criteria [65] (Fig. 2c).

To probe the model in a zero-shot setting, we quantify the importance of block *i* by masking it and measuring the resulting change in the complex embedding. Specifically, we define ATOM-ICASCORE 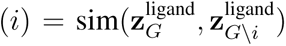, where *G* \ *i* denotes masking block *i* in graph *G* (Fig. 2d), 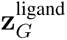 is obtained by pooling 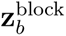 over the blocks of the bound molecule, and sim is the cosine similarity function. A lower ATOMICASCORE indicates a larger representational change and therefore higher block importance (Methods 7).

The annotated non-covalent interactions (hydrogen bonds, hydrophobic interactions, *π*-stacking, metal complexes, and halogen bonds) are derived from standard geometric interaction annotations from PLIP [65]. We compare ATOMICASCORE to the unimolecular baseline ESM-2 (3B) [4], which provides zero-shot residue importance estimates via mutation-effect scoring [66], and to LigandMPNN [67], which is conditioned on the ligand atoms and is therefore given the same context as ATOMICA. We score each residue by how much removing the ligand context changes the log likelihood of its native identity (Methods 7). We evaluate precision@10 as the fraction of residues ranked by ATOMICASCORE in the top 10 that are involved in an annotated non-covalent interaction (Fig. 2e). On an unseen test set of 5,497 protein–small-molecule complexes, ATOMI-CASCORE has the highest precision@10 among the evaluated methods at 0.423 (Fig. 2f), against 0.337 for LigandMPNN, 0.238 for ESM-2 (3B), and 0.195 for a random reference. ATOMICA retrieves residues across hydrogen bonds, hydrophobic interactions, and *π*-stacking (Fig. S1). As a further control, pretraining ATOMICA on only PDB-derived structures, with the CSD excluded, did not reduce zero-shot ATOMICASCORE performance on held-out protein–small-molecule interfaces. The ATOMICA PDB-only checkpoint reached a Precision@10 of 0.438. Thus, zero-shot ATOMICASCORE performance is robust to the inclusion or exclusion of CSD-derived structures during pretraining.

### Comparison of multimodal and interaction-type-specific pretraining

To compare multimodal and interaction-type-specific pretraining, we evaluate ATOMICA alongside otherwise identical models pretrained exclusively on a single interaction type (Fig. 2g). We then fine-tune both the multimodally pretrained model and each corresponding single-interaction-type model using the same interaction-type-specific masked block-type prediction objective, and evaluate masked block-type prediction AUPRC on test complexes selected to have low sequence similarity and low chemical similarity to any training or validation examples (Methods 8). Values are macro-AUPRC with a 95% confidence interval obtained by resampling test structures (Methods 8). After this identical fine-tuning, the multimodally pretrained model has an AUPRC of 0.707 [0.694, 0.720] on protein–DNA interfaces, compared with 0.243 [0.237, 0.248] for the interaction-type-specific model (Fig. 2h). The corresponding values are 0.552 [0.537, 0.568] versus 0.187 [0.179, 0.196] for protein–RNA interfaces and 0.666 [0.656, 0.676] versus 0.322 [0.312, 0.331] for protein–peptide interfaces. The observed difference between the multimodal and interaction-type-specific models varies across interaction types and is larger for protein–DNA and protein–RNA interfaces. With corresponding blocks masked in both conformers, ATOMICA also has higher observed macro-AUPRC than the small-molecule-only model on small-molecule–small-molecule complexes at 0.248 [0.242, 0.254] versus 0.221 [0.215, 0.227] (Fig. 2h and Methods 8). The same comparison stratified by residue burial and by intermolecular and intramolecular context is shown in Fig. S2. Across the evaluated interfaces, the multimodally pretrained model has higher observed macro-AUPRC than the corresponding interaction-type-specific model.

Because the CSD provides substantially greater exposure to common chemical motifs than the PDB, we separately compared common chemical motif recovery with and without these CSD data. We evaluated the pretrained masked-block heads of the full checkpoint and of an ATOMICA model pretrained on PDB complexes without the CSD, using held-out PDB protein–small-molecule complexes (Methods 8). When each of the 22,122 common chemical motif instances spanning 217 classes was masked once, the full model reached a top-1 accuracy of 0.808 and macro-AUPRC of 0.258, compared with 0.547 and 0.090, respectively, for the checkpoint pretrained without the CSD. The full model produced at least one top-1 prediction for 181/217 observed common chemical motif classes, compared with 30/217 classes for the model pretrained without the CSD. On this benchmark, the checkpoint pretrained with the CSD has higher top-1 accuracy and macro-AUPRC and produces top-1 predictions for more of the observed common chemical motif classes.

### Amino-acid and nucleotide sequence recovery at molecular interfaces

Whereas the preceding analysis evaluates masked block-type prediction across the shared vocabulary of molecular block classes, we next measure conventional top-1 sequence recovery within the amino-acid or canonical-nucleotide alphabet. This evaluation enables direct comparison with established protein and nucleic-acid sequence-design models, with every model scored on the same simultaneously masked positions at the interaction interface (Methods 8).

For amino-acid recovery we evaluate the pretrained ATOMICA model, without task-specific fine-tuning, since masked block-type prediction is one of the pretraining objectives. At the 10% interface-masking rate used during ATOMICA pretraining, ATOMICA has higher recovery than the highest-recovery ProteinMPNN [68] or LigandMPNN [67] configuration for each of the six protein-containing interaction types by 4.5–12.2 percentage points (Fig. 2j). Compared with five protein language models, ATOMICA has higher recovery on protein–RNA, protein–DNA, protein– peptide, protein–ion, and protein–protein interfaces, and lower recovery on protein–ligand interfaces at 69.0% versus 72.5% for ESM-2 (3B). At 30% masking, which removes an increasing fraction of the local interface context available to ATOMICA, while the other models have access to the full protein sequence, ATOMICA has higher recovery than the highest-recovery MPNN configuration on protein–RNA, protein–DNA, and protein–peptide interfaces. At 50% masking, its recovery is lower than that configuration across all six interaction types.

For nucleotide sequence recovery, we first evaluated pretrained ATOMICA. Although this model achieves high amino-acid recovery as shown above, its shared block-vocabulary head gave limited nucleotide-only recovery, reaching 45.6% on protein–RNA, 45.0% on protein–DNA and 47.5% on RNA–ligand interfaces. Since nucleotides comprise 1.2% of all maskable blocks of the ATOMICA pretraining dataset from the PDB, we fine-tuned ATOMICA on nucleotide sequence recovery. At 10% masking, ATOMICA (fine-tuned) reaches 77.6% compared with 78.0% for NA-MPNN [69] on protein–RNA interfaces, 91.0% compared with 89.3% on protein–DNA interfaces, and 85.1% compared with 82.6% on RNA–ligand interfaces (Fig. 2i). At higher masking rates, NA-MPNN has higher recovery on protein–RNA at 30% and 50% and on protein–DNA at 50%. On protein–DNA at 30%, ATOMICA (fine-tuned) reaches 85.2% compared with 84.8% for NA-MPNN, while on RNA–ligand it reaches 84.7% compared with 82.8% at 30% masking and 82.7% compared with 83.0% at 50% masking. At 10% masking, ATOMICA (fine-tuned) also has higher recovery than the RNA-only gRNAde [70] and RhoDesign [71] models, although these models do not receive the interacting protein partner. Recovery for both nucleotide and amino-acid residues at the higher masking rates of 30% and 50% is reported in Fig. S3.

### ATOMICA representations support RNA structure annotation

We evaluate ATOMICA on four RNAglib tasks [72, 73] using RNA 3D structures as input: RNA–protein binding-site prediction, RNA functional annotation, RNA–small-molecule binding-site prediction, and RNA pocket ligand classification (Fig. 3a–d). We compare with structure-based baselines (RNAglib/RGCN [73, 74] and gRNAde [70]) and RNA language models (RNAErnie [75], RNA-FM [10, 76], and RiNALMo [9]). Unless explicitly labeled “fine-tuned,” ATOMICA denotes frozen embeddings evaluated with the same MLP protocol as the frozen baseline embeddings (Methods 4).

**Figure 3.**
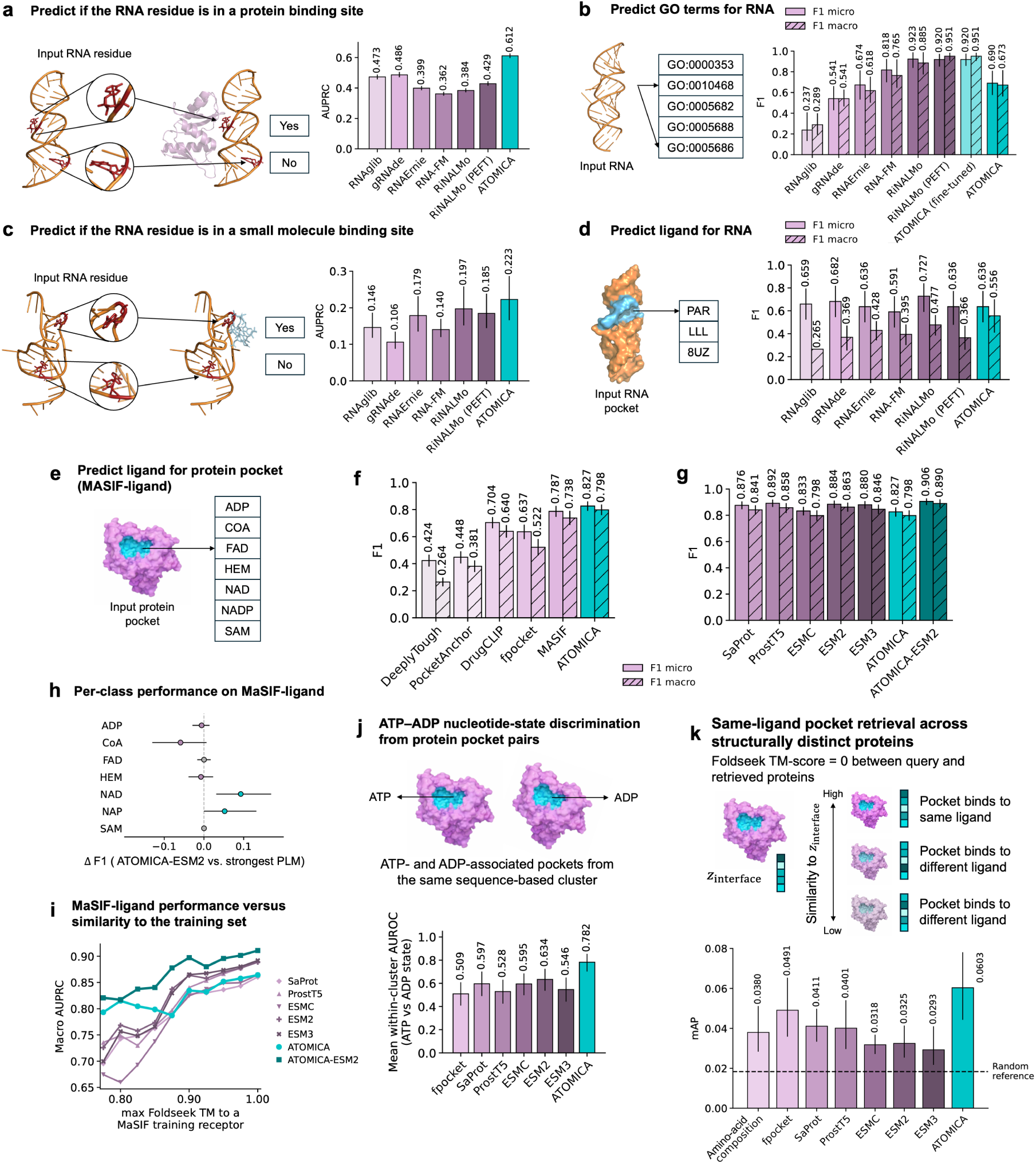
Performance of ATOMICA on protein and RNA benchmarks. **a** RNA–protein binding-site prediction, evaluated by AUPRC. **b** RNA Gene Ontology annotation, evaluated by micro- and macro-F1. **c** RNA–small-molecule binding-site prediction, evaluated by AUPRC. **d** RNA pocket ligand classification, evaluated by micro- and macro-F1. ATOMICA denotes frozen embeddings unless labeled fine-tuned. **e** Protein-pocket ligand classification across seven ligand classes. **f** Comparison with protein-pocket methods on ligand classification. **g** Comparison with protein language models, including late fusion of ATOMICA and ESM-2. **h** Observed per-class F1 difference between ATOMICA–ESM-2 late fusion and the protein language model with the highest F1 for each class. **i** MaSIF-ligand performance versus similarity to the training set across different thresholds of maximum Foldseek TM-score between training and test set receptors. **j** Mean within-cluster AUROC for distinguishing ATP- and ADP-associated pocket states. AUROC is calculated separately within each MMseqs2 sequence cluster and macro-averaged across clusters. **k** Same-ligand pocket retrieval across structurally distinct proteins, evaluated by mean average precision. Relevant pockets bind the same ligand and have no detectable Foldseek alignment to the query. Results at other Foldseek TM-score cutoffs are available in Supplementary Fig. S4h. Error bars in **a–d**, **f**, **g**, **j**, and **k** denote 95% percentile bootstrap confidence intervals for each model.

To prevent direct pretraining exposure to the molecular interaction type evaluated downstream, we use a checkpoint pretrained with all protein–RNA complexes excluded for RNA– protein binding-site prediction. For RNA–small-molecule binding-site prediction and RNA pocket ligand classification, we use a checkpoint pretrained with all nucleic-acid–ligand complexes excluded. We report only results from these interaction-type-exclusion checkpoints for these three tasks. RNA functional annotation uses the standard checkpoint because Gene Ontology (GO) annotations are never used as inputs or objectives during ATOMICA pretraining.

For residue-level RNA–protein binding-site prediction, ATOMICA achieves an AUPRC of 0.612 (95% CI [0.596, 0.627]), compared with 0.486 [0.470, 0.505] for gRNAde, which has the highest AUPRC among the evaluated baselines (Fig. 3a; test set: 10,851 residues from 190 structures). For five-label RNA functional annotation using Gene Ontology (GO) terms [77] curated from Rfam [78], ATOMICA achieves a macro-F1 of 0.673 [0.561, 0.817], below frozen Ri-NALMo at 0.885 [0.774, 0.989]. Fine-tuned ATOMICA and LoRA-adapted RiNALMo [79] both reach a macro-F1 of 0.951 and a micro-F1 of 0.920, with intervals of [0.906, 0.984] and [0.857, 0.975] for ATOMICA and [0.906, 0.984] and [0.850, 0.974] for RiNALMo, respectively (Fig. 3b; test set: 75 of 499 structures). The intervals are broad for this small test set. For residue-level RNA–small-molecule binding-site prediction, ATOMICA achieves an AUPRC of 0.223 [0.166, 0.287], compared with 0.197 [0.152, 0.258] for RiNALMo (Fig. 3c; test set: 2,158 residues from 33 structures). AUPRC remains below 0.23 for every evaluated method, indicating that this task remains unsolved. Finally, we classify RNA pockets among paromomycin (PAR), gentamycin C1A (LLL), and aminoglycoside TC007 (8UZ). ATOMICA attains a macro-F1 of 0.556 [0.410, 0.701] and micro-F1 of 0.636 [0.500, 0.773], compared with a macro-F1 of 0.477 [0.384, 0.563] for RiNALMo, which has the highest macro-F1 among the evaluated frozen baselines (Fig. 3d; test set: 44 of 290 pockets). The broad confidence intervals reflect the small, imbalanced test set. Together, these results illustrate the applicability of ATOMICA representations across diverse RNA annotation settings, spanning residue-level binding-site prediction and structure-level functional and ligand classification.

### ATOMICA and protein language models for protein pocket annotation

We evaluate protein pocket annotation using MaSIF-ligand [16], which classifies 2,509 pockets among seven ligand classes (Fig. 3e). For ATOMICA, we report frozen representations from a checkpoint pretrained after excluding protein–ligand complexes whose receptors are sequence-similar or structurally similar to the MaSIF-ligand test receptors. This checkpoint reaches a macro-F1 of 0.798 (95% CI [0.759, 0.838]) and a micro-F1 of 0.827 [0.790, 0.861]. These values exceed those of MaSIF, which reaches 0.738 and 0.787, respectively, and those of the other evaluated pocket structural encoders, but are lower than those of ESM-2 (3B), which reaches 0.863 and 0.884 (Fig. 3f,g).

Averaging the class probabilities from ATOMICA and ESM-2 (3B) [4] gives the highest observed macro-F1, micro-F1, macro-AUPRC, and macro-AUROC among the evaluated entries without training an additional fusion model (Fig. 3g). The values are 0.890 versus 0.863 for macro-F1, 0.906 versus 0.884 for micro-F1, 0.911 versus 0.889 for macro-AUPRC, and 0.982 versus 0.977 for macro-AUROC. Relative to the protein language model with the highest F1 for each class, the observed late-fusion F1 is higher for NAD and NAP, equal for FAD and SAM, slightly lower for ADP and heme, and lower for CoA (Fig. 3h).

To characterize the information this benchmark evaluates, we compare two simple label-transfer baselines that assign each test pocket the ligand class of its most similar MaSIF-ligand training receptor, identified by MMseqs2 sequence search or by Foldseek structural search, without a learned representation. Sequence label transfer reaches a macro-F1 of 0.706 and structure label transfer reaches 0.618, compared with 0.863 for ESM-2 (3B) and 0.798 for frozen ATOMICA. Of the 467 test pockets, 458 have a training receptor at 30% or higher sequence identity over at least 80% coverage, so this benchmark evaluates pocket annotation where a similar annotated receptor is available. We therefore stratify the test pockets by Foldseek TM-score to the nearest training receptor. Among the 302 pockets at TM ≥ 0.8, sequence label transfer reaches a macro-F1 of 0.812, ESM-2 reaches 0.904 and the late fusion reaches 0.915, while frozen ATOMICA reaches 0.814. Among the 165 pockets below this threshold, sequence label transfer falls to 0.401 and structure label transfer to 0.506, and ATOMICA has the highest observed macro-F1 at 0.734, compared with 0.724 for ProstT5 and 0.663 for ESM-2 (Fig. S4f,g). Across cumulative subsets defined by a maximum Foldseek TM-score to a training receptor, macro-AUPRC decreases from 0.889 to 0.726 for ESM-2 and from 0.864 to 0.793 for ATOMICA between the full test set and the 138 pockets at a maximum TM-score of 0.775, and ATOMICA has the highest observed macro-AUPRC among the single representations for every subset at a maximum TM-score of 0.85 or below (Fig. 3i). Because most MaSIF-ligand test pockets have a structurally similar training pocket, performance on this benchmark does not distinguish representations that encode local pocket geometry from those that identify a similar annotated protein.

### ATOMICA distinguishes ATP- and ADP-associated protein pocket states

We next evaluate the models on ATP- and ADP-associated protein pockets that are close in sequence but differ in nucleotide state, so that homologous proteins carry different labels (Fig. 3j). The benchmark contains 404 unique ligand-free pockets, including 223 ATP-associated and 181 ADP-associated pockets, from 120 PDB entries. Within each sequence cluster, ATP- and ADP-associated protein sequences are highly similar, with a mean pairwise sequence identity of 90.7%. These pockets are grouped into 60 sequence clusters at 50% sequence identity and 80% coverage, each containing both nucleotide states. Before representation generation, we remove the nucleotide, every other ligand, and waters, and retain only pocket positions shared between the matched ATP- and ADP-bound structures with the same residue identity (Methods 4.5). For each learned representation, we keep the pretrained encoder frozen and train a binary classifier to distinguish ATP-from ADP-associated pockets using five-fold cluster-disjoint cross-validation. To remove discrimination between sequence clusters and prevent clusters with more structures from receiving greater weight, we calculate ATP–ADP AUROC separately within each sequence cluster and report the unweighted mean across clusters.

Frozen ATOMICA reaches a cluster-macro AUROC of 0.782 across the 60 sequence clusters, compared with 0.634 for ESM-2 (3B), the highest protein language-model value. The protein language models embed the full chain and can therefore use sequence context outside the pocket, whereas ATOMICA is restricted to the local pocket representation. In the subset with concordant metal occupancy, ATOMICA reaches 0.718 across 44 evaluable clusters, compared with 0.641 for ESM-2. Within this subset, ATOMICA reaches 0.734 across 25 metal-containing clusters and 0.713 across 20 metal-free clusters. When the metal-concordant analysis is additionally restricted to pockets whose matched ATP–ADP structures are outside the ATOMICA pretraining set, ATOMICA reaches 0.778 across nine evaluable clusters, compared with 0.667 for both ESM-2 and SaProt, the highest comparator values in this subset. These analyses indicate that ATOMICA retains nucleotide-state discrimination within sequence-defined clusters after the ligand is removed, including when metal occupancy is controlled and when structures from pretraining are excluded (Fig. S4i).

### ATOMICA retrieves ligand-matched pockets across structurally distinct proteins

To test whether local pocket representations retain ligand-associated information across globally different proteins, we construct a retrieval benchmark from the held-out protein–ligand test split, whose complexes were not observed during ATOMICA pretraining (Fig. 3k). A candidate pocket counts as a relevant retrieval only if it binds the query’s ligand and has no detectable Fold-seek structural alignment [80] to the query. Thus, the ligand class of the query cannot be recovered by transferring an annotation from a structurally similar protein. Same-ligand pockets that are alignable to the query are removed from the ranking. The benchmark contains 892 pockets, 428 queries, 44 ligand classes, and 2,349 positive query–reference pairs (Methods 4.6).

Frozen ATOMICA has the highest observed macro-averaged mean average precision (mAP) among the evaluated representations on this benchmark at 0.0603 (95% CI [0.0443, 0.0781]), a 3.28-fold lift over the exact-permutation random reference, with an nDCG of 0.301 and an R-precision of 0.0586. The next-highest mAP lift is 2.67-fold for fpocket [81], while SaProt [6] and ProstT5 [7] reach 2.24-fold and 2.18-fold among the protein language models, and the full set of statistics is given in Supplementary Table S2. We further report performance at different maximum Foldseek TM thresholds between query and retrieved pairs that bind the same ligand. Relaxing this threshold progressively admits more Foldseek-alignable pockets into the relevant set. ATOMICA has the highest observed mAP among the seven evaluated representations at every fold ceiling up to and including a maximum TM of 0.3. At TM ≤ 0.3, the mAP is 0.0710 for ATOMICA and 0.0604 for ProstT5, the next highest (Fig. S4h).

### Cross-modality interface comparison for orthosteric PPI inhibitors

Orthosteric inhibitors of PPIs include small molecules that bind at a protein–protein interface and competitively block partner binding. Because orthosteric inhibitors engage the same surface features as the native partner, we compare protein-inhibitor and PPI embeddings in the ATOMICA latent space and test whether embedding similarity localizes to the inhibited PPI region.

We use the curated 2P2Idb database [82], which contains matched PPI structures and protein– inhibitor structures. After quality filtering, the analysis includes 26 protein–peptide complexes with 965 matched protein–inhibitor structures and six protein–protein complexes with 187 matched protein–inhibitor structures (Methods 5). To summarize performance, we first compute retrieval statistics for each complex and then take an unweighted mean over complexes assigned to the same 2P2Idb SuperFamily. This gives 14 protein–peptide and five protein–protein superfamilies, each of which contributes equally to the across-system summaries.

For protein–peptide complexes, we compare the ATOMICA embedding distance between peptide and inhibitor blocks, 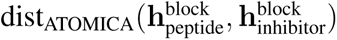, with the distance between their aligned coordinates, 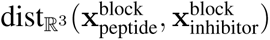 (Fig. 4a). We quantify localization using Fold Change@10, which compares the proportion of spatially close block pairs among the ten pairs with the lowest embedding distance with the reference proportion among all block pairs in the matched structures. After aggregation, Fold Change@10 is greater than one in 11 of 14 protein–peptide superfamilies (Fig. 4d), and the correlation between embedding and spatial distance is positive in 10 of 14 superfamilies (per-complex coefficients in Supplementary Table S5).

**Figure 4.**
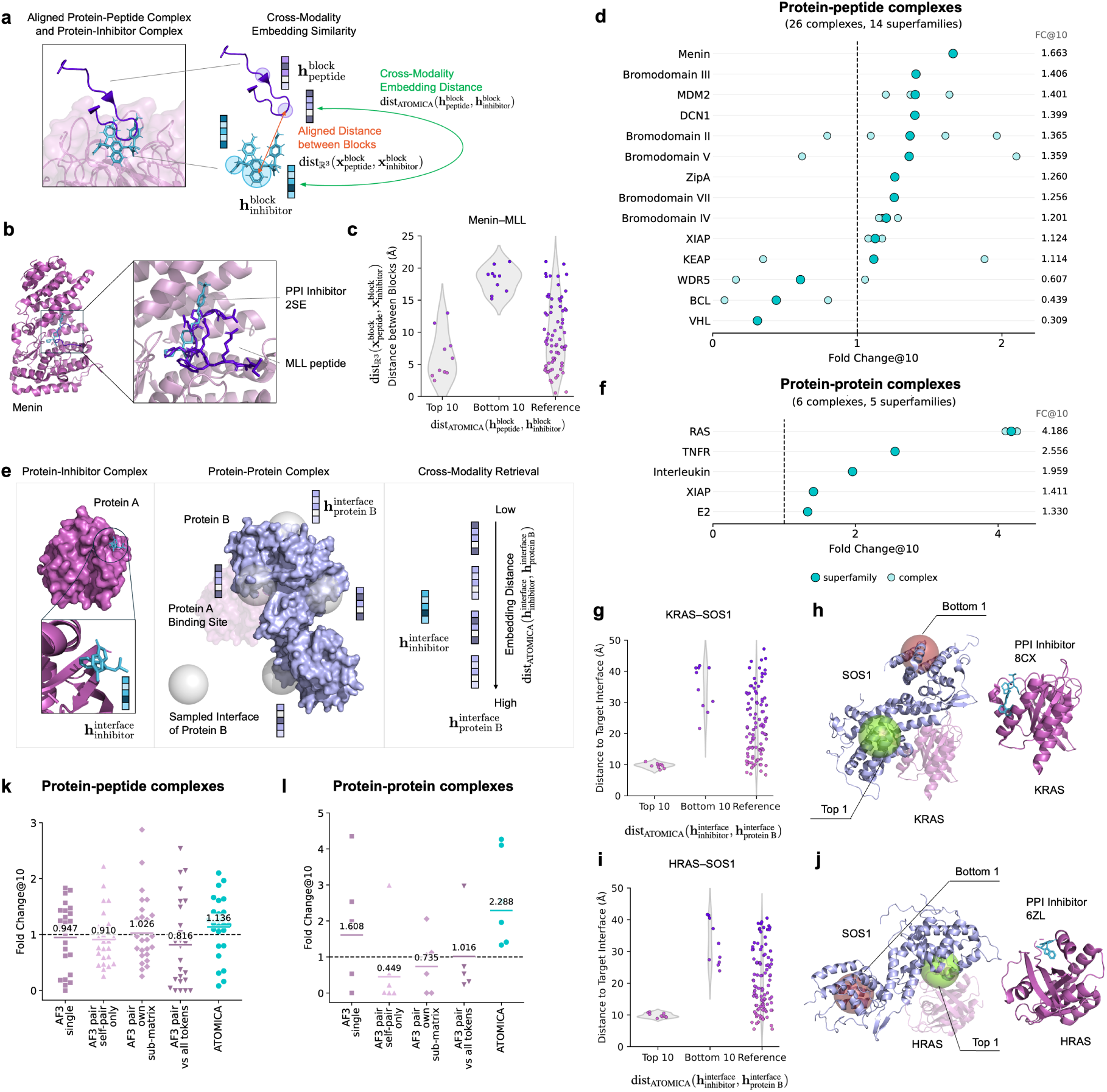
Cross-modality interface comparison of orthosteric PPI inhibitors. **a** Block-level comparison of aligned protein–peptide and protein–inhibitor structures. We compare ATOMICA embedding distance, 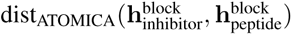, with aligned 3D distance, 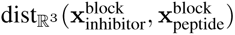. Single-atom inhibitor blocks are excluded. **b** Example alignment of the Menin–MLL interaction and Menin–MIV-7 (2SE) inhibitor complex. **c** For Menin–MLL, aligned distances for the top-10 and bottom-10 peptide– inhibitor block pairs ranked by dist_ATOMICA_; reference shows all 144 pairs. **d** Mean Fold Change@10 for retrieval of peptide– inhibitor block pairs within 4 Å across protein–peptide complexes aggregated at the superfamily level. **e** Protein–protein cross-modality retrieval. The inhibitor representation, 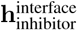, is compared with sampled protein-B interface representations, 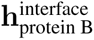, ranked by 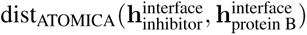. Only inhibitor ligand blocks enter the query representation, proteins A and B are never jointly embedded, and the experimental coordinates of protein A are used only to label proximity to the native interface after ranking. **f** Mean Fold Change@10 for retrieval of interfaces within 12 Å of the native A–B interface. Protein–protein complexes are aggregated at the superfamily level. **g–j** KRAS–SOS1 (**g,h**) and HRAS–SOS1 (**i,j**) examples showing top-10 versus bottom-10 retrieved SOS1 interfaces and top-1 versus bottom-1 visualizations. The two systems belong to the same RAS interaction family and are combined into one superfamily in across-system analyses. **k,l** Fold Change@10 using ATOMICA compared with the AF3 single representation and AF3 pair pooling schemes, for protein–peptide (**k**) and protein–protein (**l**) complexes. Points denote the 26 protein–peptide complexes in **k** and the six protein–protein complexes in **l**. In each panel the horizontal bar and the number above it are the mean over that level’s 2P2Idb superfamilies, 14 in **k** and five in **l**, obtained by averaging within each superfamily and then across superfamilies. Fold Change@5 and Fold Change@20 results are shown in Supplementary Figs. S5 and S6.

The Menin–MLL interaction, which plays a central role in acute leukemias [83] and is inhibited by MIV-7 (PDB ligand code 2SE) [84], illustrates the block-level comparison (Fig. 4b,c). Among the 144 inhibitor–peptide block pairs in this complex, the ten pairs with the most similar ATOMICA embeddings are closer in the aligned structures than the ten least similar pairs. These block pairs are within-complex observations used to visualize the retrieval and are not considered independent biological replicates.

For protein–protein complexes A–B, the query representation pools only inhibitor ligand blocks from the inhibitor-bound A–I complex, while the retrieval candidates are 1,000 local surface patches encoded separately from protein B (Fig. 4e). Because the inhibitor blocks are contextualized by message passing within the inhibitor-bound A–I complex, protein A can influence the inhibitor representation indirectly, although no protein-A block is pooled into the query or directly enters the similarity ranking. Proteins A and B are never jointly embedded, and the experimental coordinates of protein A are used only after ranking to label whether a patch lies within 12 Å of the native A–B interface. Fold Change@10 compares the fraction of such patches among the ten most similar candidates with their fraction among all sampled patches. After superfamily aggregation, the score is greater than one in all five protein–protein superfamilies (Fig. 4f), and the correlation between embedding and spatial distance is positive in four of five superfamilies (per-complex coefficients in Supplementary Table S6). For KRAS–SOS1 (Fig. 4g,h) and HRAS–SOS1 (Fig. 4i,j) complexes, which both belong to the RAS interaction family, the top-ranked SOS1 surface patches are closer to the native RAS–SOS1 interface than the bottom-ranked patches for a covalent KRAS inhibitor [85] and a small-molecule HRAS inhibitor [86].

We further compare ATOMICA with the internal single and pair representations from the AlphaFold3 (AF3) Pairformer trunk [1] by substituting each representation into the same retrieval analysis (Methods 5). For protein–peptide retrieval, the mean Fold Change@10 over the 14 super-families is 1.136 for ATOMICA, 0.947 for the AF3 single representation, and 1.026 for the AF3 pair representation with the highest mean, which uses own-sub-matrix pooling. The corresponding unweighted means over the 26 individual complexes are 1.148, 0.988, and 1.081 (Fig. 4k). Fold Change@10 is greater than one in 11 of 14 superfamilies for ATOMICA, compared with 7 of 14 and 5 of 14 for these two AF3 representations. For protein–protein retrieval, the mean over the five superfamilies is 2.288 for ATOMICA, 1.608 for AF3 single, and 1.016 for the AF3 pair representation with the highest mean, which uses all-token pooling. The means over the six complexes are 2.605, 1.868, and 1.016 (Fig. 4l), and Fold Change@10 is greater than one in 5 of 5, 3 of 5, and 2 of 5 superfamilies. These descriptive comparisons support the utility of the ATOMICA representation for cross-modal interface comparison.

### Prediction of ion and cofactor identities in dark proteome pockets

Ion and cofactor binding sites can recur across diverse folds and sequences [87, 88]. Proteins in the dark proteome have limited functional annotation, and many have limited sequence or structural similarity to characterized proteins. We define these proteins using Foldseek-defined dark clusters [44–46]. In this setting, we use ATOMICA to generate interaction-based hypotheses of ligand identity from local pocket geometry, leveraging patterns learned from intermolecular interfaces across molecular modalities rather than relying solely on sequence-derived homology, which provides limited coverage for proteins in the dark proteome [89]. Among AFDB Foldseek clusters, 711,705 (30.9%) are classified as dark clusters (Fig. 5a) [45, 90] and are enriched for proteins with limited functional annotation [45, 91]. We use predicted ligand identities as functional signals to prioritize biochemical hypotheses and guide follow-up experiments.

**Figure 5.**
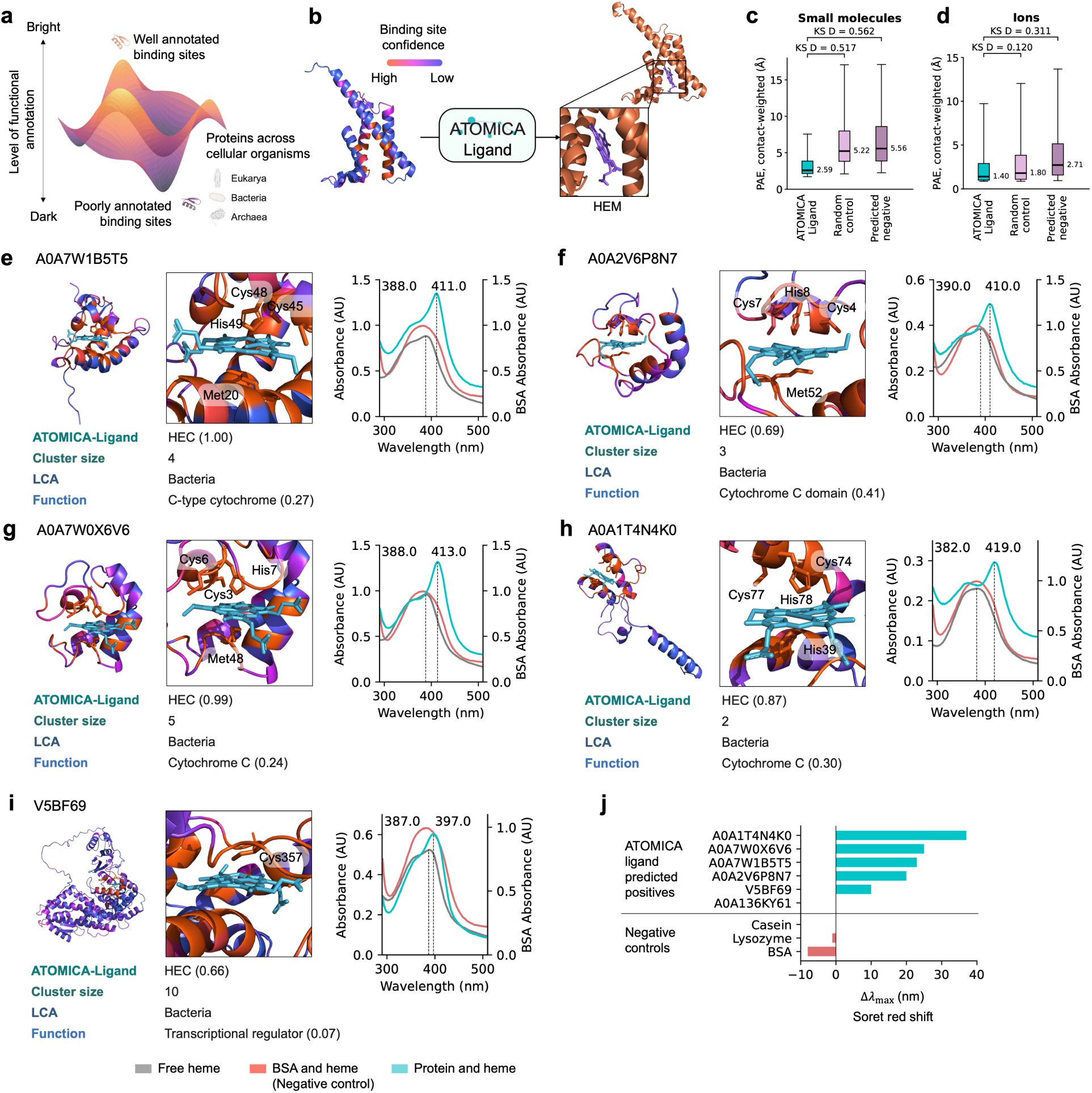
Annotation of dark proteome with ATOMICA and experimental evaluation of predicted heme binders. **a** Annotation of binding sites across proteins in cellular organisms. **b** Prediction of ligand identities for metal-ion and small-molecule binding sites in the dark proteome. **c** Contact-weighted AlphaFold3 ligand–pocket PAE for small molecules, comparing ATOMICA-Ligand predictions with a random control and predicted-negative controls. Brackets report two-sample Kolmogorov–Smirnov *D*. **d** Contact-weighted AlphaFold3 ligand–pocket PAE for metal ions, with the same controls and statistic. **e–i** Predicted structures and UV–vis spectra for five ATOMICA-Ligand-nominated heme candidates. Dashed lines indicate the Soret-band *λ*_max_ of free heme and protein with heme. All five show red shifts relative to free heme consistent with heme association under the assay conditions: **e** A0A7W1B5T5, **f** A0A2V6P8N7, **g** A0A7W0X6V6, **h** A0A1T4N4K0, and **i** V5BF69. **j** Soret-band shift, Δ*λ*_max_ relative to free heme, for all six tested candidates and protein negative controls. Additional negative controls are shown in Supplementary Fig. S8.

We fine-tune ATOMICA-Ligand to predict ligand identity from pocket geometry and apply it to high-confidence AF2 representatives [92] (pLDDT*>*90) from 33,482 dark clusters. We apply PeSTo [93] to predict candidate ion and small-molecule pockets, yielding 2,851 proteins with ion-binding sites and 969 with small-molecule sites. ATOMICA-Ligand predicts identities for 9 metal ions and 12 common cofactors for protein pockets (Fig. 5b, Methods 6), with predictions for 2,565/2,851 proteins and 81/969 proteins, respectively. The lower small-molecule assignment rate reflects the narrower classifier vocabulary. The 12 small-molecule classes cover 29.5% of QBioLiP protein–small-molecule interfaces, whereas the nine ion classes cover 95.9% of protein–ion interfaces.

To assess cross-model concordance, we fold protein–ligand complexes formed by pairing each predicted pocket with the ligand assigned by ATOMICA-Ligand using AlphaFold3 [1]. Compared with matched random controls, ATOMICA-Ligand-assigned pairs reach lower PAE_pocket-ligand_ (Methods 6) for small molecules, with a median of 2.59 Å against 5.22 Å (Kolmogorov–Smirnov *D* = 0.517), and for ions, with a median of 1.40 Å against 1.80 Å (*D* = 0.120). The ion shift is more modest, while comparison with ATOMICA-predicted-negative controls gives *D* = 0.562 for small molecules and *D* = 0.311 for ions (Fig. 5c,d). Because both models use PDB-derived structural data, we interpret their agreement as cross-model concordance. In Section S1, we use these pocket-level predictions to interpret putative biochemical roles.

### Experimental evaluation of candidate heme binders predicted by ATOMICA-Ligand

Heme is a cofactor involved in electron transfer, oxygen transport, enzymatic catalysis, and cellular signaling, and it accounts for 3.4% of protein–small-molecule interaction complexes in the ATOMICA dataset. We use ATOMICA-Ligand to nominate heme binders in the dark proteome and prioritize nine candidates using the ATOMICA-Ligand score, AF3 ipTM confidence, and experimental feasibility. Six candidates were produced at sufficient yield for experimental testing (Methods 10.1).

Across the 969 screened dark-proteome proteins, HEM and HEC scores are uncorrelated with coverage-weighted sequence identity to the corresponding training positives, with Spearman *ρ* = −0.064 and +0.017, respectively. HEM scores show a modest association with Foldseek similarity to heme-bound training receptors (*ρ* = 0.231), but little association with similarity to any training receptor overall (*ρ* = 0.041). Only two of the 65 predicted heme binders have a significant training-set heme hit at *E* ≤ 10^−3^ under MMseqs2, phmmer, or jackhmmer. Both are detected only by phmmer, with E-values of 1.4 × 10^−4^ and 9.3 × 10^−4^. These controls show that the classifiers retain heme discrimination without requiring a same-cofactor training alignment (Methods 6).

We assess heme association through shifts in the Soret-band maximum relative to free heme. Five of the six prioritized candidates produce Soret red shifts consistent with heme association under the assay conditions (Fig. 5e–i; Methods 10.2). We summarize the spectral response as Δ*λ*_max_ = *λ*_max_(protein and heme) − *λ*_max_(free heme) (Fig. 5j). Across the five candidates with Soret red shifts, Δ*λ*_max_ ranges from +10 to +37 nm. The sixth tested candidate, A0A136KY61, was nominated by the HEM classifier with a score of 0.335 but did not produce a Soret red shift (Fig. S8a).

We additionally assayed BSA, lysozyme, and casein as protein negative controls. BSA and lysozyme are scored as negative by ATOMICA-Ligand for both HEM and HEC (scores of 0.00), whereas no PeSTo pocket was detected for casein and therefore no ATOMICA-Ligand assignment was made. None of these three controls produced a positive Soret shift relative to free heme (Fig. 5j and Fig. S8b–d). Thus, the five candidates with the spectral phenotype are separated from both the unsuccessful sixth candidate and the experimentally assayed negative controls by the direction and magnitude of the Soret shift.

Four of the five candidates with Soret red shifts are assigned covalent heme by ATOMICA-Ligand, comprising A0A7W1B5T5 (ATOMICA-Ligand score = 0.997), A0A2V6P8N7 (ATOMICA-Ligand score = 0.690), A0A7W0X6V6 (ATOMICA-Ligand score = 0.992), and A0A1T4N4K0 (ATOMICA-Ligand score = 0.874) (Fig. 5e–h). UniProt [94] assigns cytochromec-related function and heme-binding terms to these candidates through automatic rule-based assertion (ARBA/UniRule) [95], which is driven primarily by sequence-derived family or domain signatures and taxonomy. These annotations are sequence-derived predictions rather than experimentally validated assignments. We nevertheless assayed these candidates, and their Soret red shifts provide evidence consistent with heme association under the assay conditions rather than validation of cytochrome function. ATOMICA complements these assignments by predicting heme identity from 3D pocket geometry.

To test whether ATOMICA-Ligand is capturing information beyond the canonical CXXCH motif found in many cytochrome c proteins [96], we performed a motif-controlled evaluation. We retained only proteins that contain CXXCH, such that the motif alone provides no discrimination, and ATOMICA-Ligand still separates heme binders from non-binders with AUROC 0.849 (vs. 0.500 for a constant CXXCH-motif-only predictor) (Fig. S9a).

Dark protein V5BF69 (ATOMICA-Ligand score = 0.663) does not contain the canonical CXXCH heme-binding motif. In the presence of protein, it shows a Soret maximum of 397 nm compared with 387 nm for free heme, corresponding to Δ*λ*_max_ = +10 nm (Fig. 5i,j). The sequence around the AF3-predicted heme-coordinating cysteine (<u>C</u>), AHGV<u>C</u>AG, is not captured by other common heme sequence motifs (GX[HR]X<u>C</u>[PLAV]G [96], FXXGXRX<u>C</u>XG [97]). Although V5BF69 contains partial overlap with motif families through the <u>C</u>XG pattern, a CXG-motif-controlled analysis shows that among proteins containing CXG, ATOMICA-Ligand still separates heme binders from non-binders with AUROC 0.990 (vs. 0.500 for a constant CXG-motif-only predictor) (Fig. S9d).

## Discussion

ATOMICA is based on the premise that intermolecular interfaces provide a transferable basis for representation learning across biomolecular interaction settings. Its multiscale representation connects interfaces formed by proteins, nucleic acids, small molecules, and ions within the ATOMICA latent space at the atom, block, and graph levels, and this shared representation organizes by interaction type, supports zero-shot identification of interface residues, and recovers native sequence identity. The same representation enables direct comparison of chemically different interfaces and molecular annotation from local pocket structure, extending to experimentally validated hypothesis generation in the dark proteome.

Multimodal pretraining is particularly beneficial for protein–DNA and protein–RNA interfaces, and evaluations with frozen interaction-type-exclusion checkpoints show that performance on the corresponding RNA tasks does not require direct pretraining exposure to matching interaction types. In protein pockets, the observed performance improvements from combining ATOMICA with ESM-2 support complementary structural and sequence information. Stratifying MaSIF-ligand by structural similarity to the training set shows that protein language models reach higher values where a similar training receptor is available, whereas the performance of ATOMICA depends less on this similarity. The ATP–ADP and ligand-matched retrieval analyses further show that the representation remains informative when local geometry distinguishes ligand-free states or shared ligand identity without detectable global structural alignment.

The ATOMICA latent space enables direct comparison across molecular modalities and hypothesis generation from local pocket structure. In the evaluated orthosteric inhibitor systems, inhibitor embeddings retrieve regions near native peptide and protein interfaces, relating chemically different partners that engage a common target interface. The inhibitor and native partner are not a physical binding pair, so the analysis compares their interfaces instead of predicting a direct interaction. Physics-based simulations could characterize the target–inhibitor and target–partner complexes separately, but they would not provide the cross-modal similarity measure evaluated here. For dark-proteome annotation, ATOMICA-Ligand assigns candidate ions and cofactors to pockets with limited functional annotation using local pocket structure. Heme discrimination is retained on held-out pockets without detectable same-cofactor training homologs, and screen scores show little relationship with sequence similarity to training positives. Five selected candidates show Soret-band shifts consistent with heme association, connecting ATOMICA predictions from local pocket structure to an experimental readout.

Our dataset of molecular interaction interfaces has several limitations. CSD-derived pairs are crystal-packing contacts rather than functional biological complexes, although their diverse non-covalent geometries provide useful pretraining examples. ATOMICA depends on access to accurate molecular structures, which remain unavailable for many complexes because of crystallization challenges, conformational heterogeneity, and disorder. Structure predictors including Al-x phaFold3 [1] and RoseTTAFold [2] may expand the set of interfaces that can be represented. Structure prediction, physics-based methods, and ATOMICA play complementary roles. Structure prediction proposes candidate complex geometries, molecular dynamics samples the conformational dynamics of selected systems, and free-energy calculations estimate thermodynamic differences between specified states. ATOMICA provides reusable interface representations for comparison, retrieval, and annotation across molecular systems and modalities, while selected hypotheses can be examined using physics-based methods when dynamical or thermodynamic characterization is needed.

The empirical scope of several analyses is also limited by the number of available systems and the form of validation. The comparison of cross-modal orthosteric inhibitors is limited to 14 protein–peptide and five protein–protein superfamilies, with multiple inhibitor queries and structures nested within these systems, so differences between ATOMICA and the internal representations of the evaluated AF3 remain unresolved. In dark-proteome analysis, the confidence of AF3 provides cross-model concordance rather than independent validation because both models use structural data derived from PDB. The heme candidates were jointly prioritized using ATOMICA-Ligand, AF3 confidence, and experimental feasibility. Their Soret-band shifts support heme association under the assay conditions but do not establish affinity, stoichiometry, binding-site specificity, or physiological function. Extending this evaluation to additional ions and cofactors will require broader experimental testing.

Understanding molecular interactions requires representations that capture local geometry and chemistry across diverse molecular partners. Here we introduce ATOMICA, an interaction-centered geometric model pretrained on a large multimodal dataset, and show that its latent space supports cross-modality comparison and fine-tuning across modalities. These results establish that multimodal interface pretraining can support more unified representations of biomolecular interactions across molecular classes.

## Data availability

Datasets are available on Harvard Dataverse at https://doi.org/10.7910/DVN/4DUBJX.

## Code availability

The code used to reproduce the results, along with documentation and usage examples, is available at https://github.com/mims-harvard/ATOMICA. The model weights are available at https://huggingface.co/ada-f/ATOMICA. Details are also available at https://zitniklab.hms.harvard.edu/projects/ATOMICA. The version corresponding to this manuscript is ATOMICA v1.0.0 and is archived on Zenodo [98]. The code is released under the MIT License.

## Acknowledgements

A.F. is supported by the Kempner Graduate Fellowship at Harvard University. M.D. is supported by an NSERC/FRQNT Postdoctoral Fellowship at the Massachusetts Institute of Technology. We gratefully acknowledge the support of NIH R01-HD108794, NSF CAREER 2339524, US DoD FA8702-15-D-0001, awards from Harvard Data Science Initiative, Amazon Faculty Research, Google Research Scholar Program, AstraZeneca Research, Roche Alliance with Distinguished Scientists, Sanofi iDEA-iTECH, Pfizer Research, Chan Zuckerberg Initiative, John and Virginia Kaneb Fellowship at Harvard Medical School, Harvard Medical School Dean’s Innovation Fund for the Use of Artificial Intelligence, and Kempner Institute for the Study of Natural and Artificial Intelligence at Harvard University. J.L. is supported, in part, by NIH grants HL155107, HL166137, and HG007691; by AHA grants AHA957729 and AHA24MERIT 1185447; and EU HorizonHealth 2021 grant 101057619 (REPO4EU). M.Z., B.L.P., and M.D. are supported, in part, by the Biswas Computational Biology Initiative in partnership with the Milken Institute. Any opinions, findings, conclusions, or recommendations expressed in this material are those of the authors and do not necessarily reflect the views of the funders.

## Author contributions

A.F. developed and implemented the ATOMICA model, conducted extensive benchmarking, and performed detailed analyses of its performance and applications. Z.Z. contributed to the interpretability framework and assisted with the development of the ATOMICASCORE analysis. A.Z. assisted with the evaluation of ATOMICA. J.L. provided guidance on the evaluation of ATOMICA. M.D. and B.L.P. performed the experimental evaluation of heme association for ATOMICA-Ligand candidates. A.F. and M.Z. designed the study. All authors contributed to writing and editing the manuscript.

## Methods

The Methods describe (1) dataset curation, (2) the hierarchical all-atom architecture, (3) training of ATOMICA, (4) RNA and protein annotation benchmarks, including protein-pocket classification, ATP–ADP discrimination, and pocket retrieval, (5) cross-modality PPI-inhibitor and AF3 comparisons, (6) dark-proteome analysis with ATOMICA-Ligand, (7) quantifying interface-block importance with ATOMICASCORE, (8) masked block-type prediction and sequence recovery, (9) metrics and statistical analyses, and (10) experimental evaluation of selected heme candidates.

### 1 Curation of datasets

#### 1.1 Small molecule structures

We extract structures of small molecule interactions from the 2023 release of the Cambridge Structural Database (CSD). We retain CSD entries that are organic and non-polymeric, have 3D coordinates, have no disorder, errors, or metals, are described by a single SMILES string (in other words, each crystal contains only one chemical compound), and consist of molecules with 6–50 heavy atoms. CSD entries are unit cells of infinitely repeating crystal lattices. For our purposes of learning intermolecular interactions, we sample pairs that represent the intermolecular interactions in a given unit cell. Given an entry of the CSD, we iterate through each unique conformer in the unit cell and extract all pairs of interactions with neighboring peripheral conformers that are within 4 Å of the central conformer using the CSD Python API. In total, there are 1,767,710 structures of molecular pairs from 375,941 CSD entries. Inspired by fingerprint-based similarity measures used in chemistry [99], we use a one-hot encoding of the molecular complex from a vocabulary of 290 common chemical motifs [53] (which is later used in our assignment of blocks to small molecules). We use Manhattan distance between the embeddings to sample 1,000 molecular complexes and their 100 nearest neighbors for both the validation and test splits, resulting in 10,000 molecular complexes for both splits, that are distinct from the training set. Manhattan distance is applied to split small molecules into train/validation/test sets because it measures molecular similarity based on our vocabulary of common chemical motifs and is fast to compute at our scale, enabling large-scale nearest-neighbor retrieval for dataset construction.

#### 1.2 General biomolecular structures

We extract the structures of the interacting molecules from Q-BioLiP (June 2024). This includes structures of proteins that interact with ions, ligands, DNA, RNA, peptides, and proteins, and nucleic acids interacting with ions and ligands from the Protein Data Bank (PDB). For proteins, DNA, and RNA, we crop the complex to keep all residues within 8 Å of any atom, amino acid, or nucleic acid residue in the other molecule. In total, there are 124,541 protein–protein interaction complexes, 119,017 protein–small-molecule interaction complexes, 74,514 protein–ion interaction complexes, 8,475 protein–peptide interaction complexes, 5,185 nucleic acid–ligand interaction complexes, 3,511 protein–RNA interaction complexes, and 2,750 protein–DNA interaction complexes. For protein–ion, protein–small-molecule, protein–peptide, and protein–protein molecular complexes, we cluster each set of complexes at 30% protein sequence identity using MMseqs2 with a coverage of 80%, sensitivity of 8, and cluster mode 1[100]. For protein–protein complexes, we also ensure that for any two complexes in different clusters, there is a maximum of 30% sequence identity between all chains in the two complexes. For protein–RNA and protein–DNA complexes, we cluster at 30% protein sequence identity and 30% nucleotide sequence identity using MMseqs2 with the same settings as above. This ensures that complexes in different clusters have a maximum of 30% protein sequence identity and 30% nucleotide sequence identity. For nucleic acid–ligand structures, we cluster at 30% nucleotide sequence identity. Finally, we split clusters into train, validation, and test splits using an 8:1:1 ratio.

#### 1.3 Construction of hierarchical graphs of interacting molecules

Given the atomic structure of two molecules interacting, an atom-level graph is constructed. Each atom in the complex maps to an atom node in the graph with the following features: element and 3D coordinates of the atom. Intramolecular atom edges are defined for each atom to the *k* nearest atoms in the same molecule. Intermolecular atom edges are defined for each atom to the *k* nearest atoms in the other molecule. In total, there are 118 atom types based on the elements of the periodic table. We additionally define a special mask atom used when masking a block during pretraining and evaluation.

Atom nodes are connected to block nodes. The block nodes have the following features: block type and 3D coordinates of the block given by the mean of the atomic coordinates of the atoms in the block. Each atom is connected to one block node. For proteins, peptides, DNA, and RNA, we define the atoms that belong to a given block by the amino acid and nucleotide residues.

For small molecule ligands, we follow the definition of Kong et al. [53] to group atoms of small molecule ligands into blocks. In their framework, small molecule ligands were tokenized into blocks representing common chemical motifs using a graph-based Byte Pair Encoding algorithm that learns a vocabulary from ligands found in the PDB. This results in 290 common chemical motifs. Atom nodes are then grouped into blocks by matching against the predefined vocabulary of common chemical motifs. The algorithm greedily merges neighboring atoms into the most frequent common chemical motif present in the vocabulary until no further vocabulary matches are found. Each identified common chemical motif is assigned its corresponding vocabulary symbol, and its constituent atoms are grouped into a single block. Sections of the molecule that cannot be assigned to the vocabulary become blocks comprising one atom. This hierarchical representation enables the model to process the common chemical motifs of a ligand as blocks rather than as individual atoms, and conveys the covalent connectivity of its atoms to the model.

Blocks therefore vary in atom count and physical scale across modalities and represent chemically defined units rather than uniformly sized molecular units.

Intramolecular block edges connect each block to the *k* nearest blocks in the same molecule. Intermolecular block edges connect each block to the *k* nearest blocks in the other molecule. In total, there are the following block types: 20 for canonical amino acids, 4 for DNA nucleotides, 4 for RNA nucleotides, 290 for common chemical motifs, and 118 for elemental blocks.

In addition, there are three special block types: mask, unknown, and global. The mask node is used during pretraining for masked block-type prediction. Unknown nodes are used for nodes that do not fall into the defined vocabulary, such as non-canonical amino acids and nucleotides. There are also two global-type nodes, one at the atom level and one at the block level. The two global nodes are connected to all nodes in each molecule at their respective level.

### 2 Hierarchical all-atom geometric deep learning model

#### Overview

ATOMICA uses an SE(3)-equivariant 3D message passing network on graphs of molecular complexes to learn representations that are informative of the intermolecular interactions between molecules.

Let *D* = {*G^i^* | *i* = 1*, …, N*} denote a pretraining dataset of molecular-complex graphs. Our goal is to pretrain a model *F* on *D* such that it generates chemically informative representations h*_i_* = *F*(*G^i^*) for every intermolecular complex *G^i^*. In addition to representation learning, *F* can also be fine-tuned on datasets of labeled graphs of molecular complexes, such as 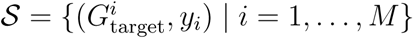, where *M* ≪ *N*, to predict *y_i_* for every 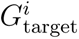.

#### Atom-level representation learning

Here we outline the SE(3)-equivariant 3D message passing network for ATOMICA on the nodes of the graph *G^i^*. Several rotational equivariant neural networks have been introduced for modeling molecules [54, 101–103]. We build on the E(3)equivariant neural network layers presented by tensor-field networks implemented in e3nn [104] and DiffDock [56]. Message passing for the intermolecular edges and intramolecular edges is done separately, but the message passing framework for the two edge types is the same.

The feature vector of atom 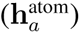 node *a* in *G^i^* is a geometric object comprised of a direct sum of irreducible representations of the O(3) symmetry group. The feature vectors 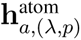 are indexed with *λ, p*, where *λ* = 0, 1, 2*, …* is a non-negative integer denoting the rotation order and *p* ∈ {o, e} indicates odd or even parity, which together index the irreducible representations (irreps) of O(3). In the ATOMICA model, we set *λ*_max_ = 1 for 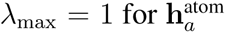, and we denote the number of scalar (0e) and pseudoscalar (0o) irrep features in 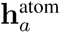 with ns, and the number of vector (1o) and pseudovector (1e) irrep features in 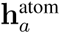 with nv.

The atom-type of node *a*, determined by the element of the atom, is embedded with a normal distribution and trainable weights as a scalar ns × 0e. There are *L*_GNN_ layers of message passing between atom nodes. At each layer *l*, the node updates for node *a* in the graph of interaction complex *G^i^* are given by:

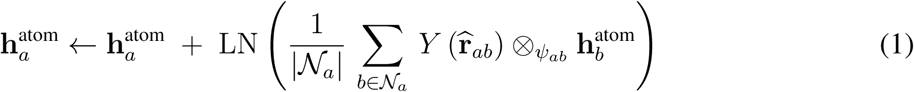

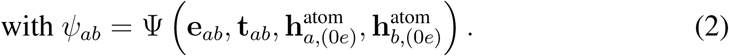

where LN is layer norm. After each layer *l* of message passing, 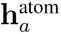 is filtered down to irreps with *λ*_max_ = 2. After *L* layers, the 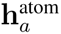 embedding is projected with a 2-layer MLP to a *d*_node_-dimensional vector.

#### Block-level representation learning

The feature vector of block 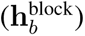 node *b* in *G^i^* is also a geometric object defined in the same way as 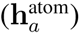. We initialize block nodes using a scalar, ns × 0e, trainable embedding of block types.

Let *d*_node_ be the dimension of 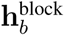 and *n*_heads_ be the number of attention heads. We define *d_h_* = *d*_node_*/n*_heads_ as the dimension per head. The multi-head cross-attention operation can be expressed as:

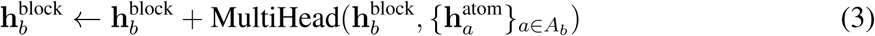

where *A_b_* is the set of atoms in block *b*, and MultiHead is defined as:

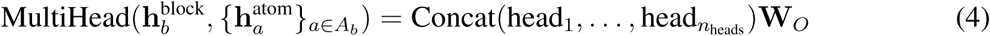

and each head is computed as:

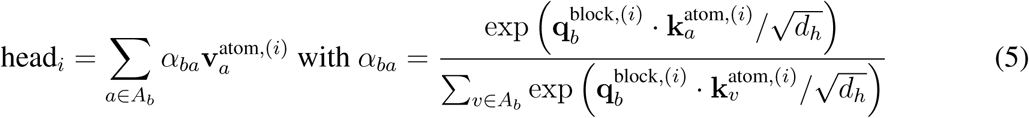

where 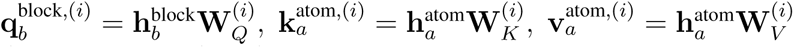, and 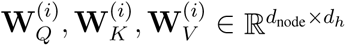 and 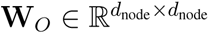. Message passing between the block nodes is specified by the same architecture as the atom nodes described in equation (1) and has separate model parameters.

#### Graph- and interface-level representations

Throughout, **h** denotes representations learned by ATOMICA, with **h**_(_*_λ,p_*_)_ denoting their individual irrep components when these are referred to explicitly. For the block-, graph-, and interface-level representations used for direct embedding comparisons, we construct a *d*_node_-dimensional representation from the *λ* = 0 components of the equivariant block features and retain the notation 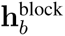 for this representation.

To pool 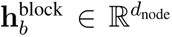 for blocks *b* ∈ *G^i^* into a graph-level representation 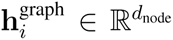, we apply multi-head self-attention for *L*_pool_ layers and sum the resulting block representations over all *b* ∈ *G^i^*. When the same pooling operation is applied only to the blocks belonging to one molecular interface A, we denote the resulting representation as 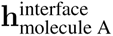. Thus, **h**^graph^ and **h**^interface^ are obtained using the same pooling operation and differ only in the set of blocks that is pooled. These representations are used for the direct cross-modality embedding comparisons in Fig. 4.

#### Rotation-invariant representations for downstream tasks

The graph- and interface-level representations above are constructed from the *λ* = 0 components of the learned equivariant features. The underlying ATOMICA representations **h** also contain *λ >* 0 channels generated during equivariant message passing. For downstream prediction and frozen-representation benchmarks, we construct deterministic rotation-invariant representations **z** from **h** so that information carried by these higher-order channels is retained without introducing dependence on the orientation of the input structure.

For an atom feature 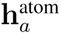, we define the atom-level invariant representation 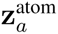 by concatenating the 0e and 0o components with the within-degree Gram entries

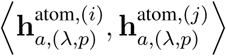

over all pairs of channels *i ≤ j* belonging to the same irrep (*λ, p*) with *λ >* 0. These quantities are invariant to global rotation because ⟨*D^λ^*(*R*)**x***, D^λ^*(*R*)**y**⟩ = ⟨**x**, **y**⟩ for any rotation *R*, and together with the *λ* = 0 channels they form the complete set of rotation invariants of degree two in **h**. Inner products between channels of different rotation degree are not invariant and are therefore not included. Invariance to translation is inherited from the architecture because all messages are constructed from relative coordinates.

The block-level invariant representation 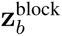 concatenates three components: (1) the *d*_node_-dimensional block representation 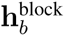; (2) the within-degree Gram entries of the equivariant block feature, constructed analogously to the atom-level representation; and (3) the mean and standard deviation of 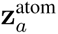 over the atoms *a* ∈ *A_b_* belonging to the block. The standard deviation is retained because it captures heterogeneity among the atomic environments within a block that is discarded by the mean alone.

Graph-level and interface-level invariant representations 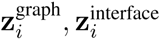, are pooled from
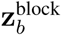 without learned parameters. We use two fixed pooling rules according to how the representation is subsequently evaluated, rather than selecting the pooling separately for individual tasks. When a prediction head is trained on top of the representation, 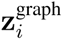 is the concatenation of the mean over blocks, the per-dimension standard deviation over blocks, and the representation of the global block node. For direct comparison of frozen representations without fine-tuning, 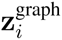 is formed from the mean over blocks, with each of its three constituent components independently *L*_2_-normalized to unit length before concatenation. Component normalization prevents differences in numerical scale between the three components from dominating the resulting distance.

#### Self-supervised learning with denoising and block-type masking

Node-level denoising has been an effective pretraining objective on 3D molecular datasets of DFT-generated structures, where it counteracts over-smoothing in GNNs [105], and it has been shown to be related to learning a force field of per-atom forces [59, 106]. In addition, denoising is linked to score matching, which has been used to train generative models [56, 107] and for unsupervised binding-affinity prediction [61]. This motivates the use of denoising as an objective for self-supervised training.

Each *G^i^* ∈ *D* is composed of atom and block nodes from two interacting molecules. We write *G̃^i^* for the perturbed graph created by applying two transformations to one molecule in *G^i^*:

- Rigid rotation and translation: A rotation vector ***ω*** ∼ *p*(***ω***) = *N_SO_*_(3)_ is sampled, and we rotate all atom and block coordinates of the selected molecule about its center. A translation vector is sampled 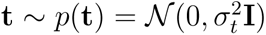 and we apply this translation to all atom and block coordinates of the selected molecule.
- Torsion angle noising: Torsion angles are sampled 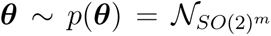 where *m* is the number of rotatable bonds in the molecule. For peptides, proteins, RNA and DNA we only perturb rotatable bonds in the side chain.

Noise for the SE(3) and torsion perturbations is sampled whenever a complex is loaded, and the perturbed coordinates replace the clean coordinates. Within every forward pass, atom-level edges are rebuilt from the perturbed coordinates, while block centroids and block-level edges are recomputed from those coordinates. No clean coordinates or precomputed or cached edge list is supplied to or reused by the model.

To predict the rotation score **s*_ω_*** ∈ ℝ^3^ and the translation score **s_t_** ∈ ℝ^3^ from *G̃^i^*, the node representations at the atom and block level are convolved with the center of the graph using a tensor field network [56]:

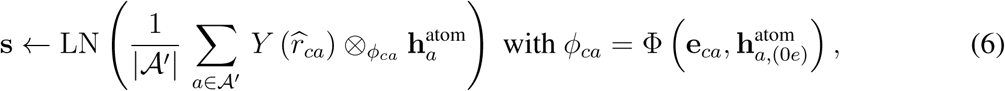

where nodes *a* ∈ *A^′^* are the atom nodes in the perturbed molecule and *c* is the center of the perturbed molecule. This is a weighted tensor product, with the weights given by a 2-layer MLP, Φ, which takes as input the Gaussian smearing *d*_edge_-embedding of the Euclidean distance between coordinates of the center *c* and node *a*, and the scalar component of 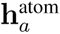.

Finally, the rotation score is given by the pseudovector irrep component 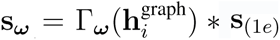 and the translation score is given by the vector irrep component 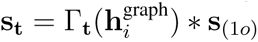, where Γ***_ω_*** and Γ**_t_** are 2-layer MLPs that project the graph representation of *G^i^* to a single scalar.

To predict the torsion score **s*_θ_*** ∈ ℝ*^m^*, the atom nodes are convolved with the center of the rotatable bonds connecting atoms *a_z_*_0_*, a_z_*_1_. Let *z* denote the center of one of the rotatable bonds. We define 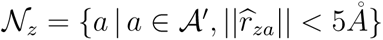 as the set of all atoms in the perturbed molecule within 5 Å of the center of the bond:

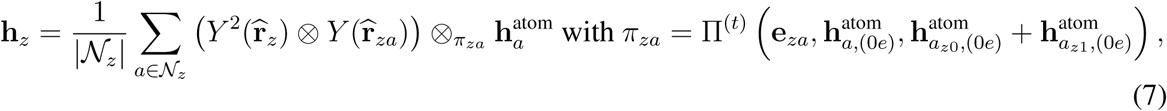

The first tensor product is between the second-order irreps of the unit direction vector along the two atoms *a_z_*_0_*, a_z_*_1_ of the bond *z*, *Y*^2^(**r̂***_za_*), and the unit direction vector between the center of the bond and atom *a*, *Y* (**r̂***_za_*). This is followed by a weighted tensor product with the weights given by a 2-layer MLP, Π, which takes as input the Gaussian smearing *d*_edge_-embedding of the Euclidean distance between coordinates of the bond center *z* and node *a*, the scalar component of 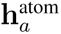, and the sum of the scalar components of the two atoms in the bond, 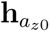 and 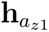. Finally, we sum the scalar and pseudoscalar components of **h***_z_* and project the result to a single scalar 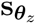 using a 2-layer MLP.

We calculate the loss components as follows:

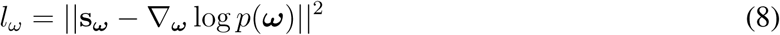

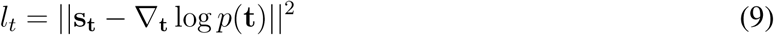

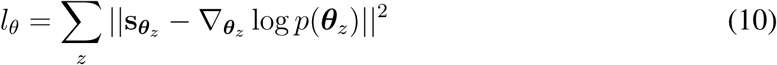

where 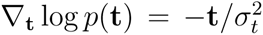. The values of ∇**_t_** log *p*(**t**) and 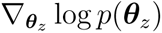 can be calculated by precomputing a truncated infinite series following [56, 61].

In addition to denoising, we also pretrain the model by masking out block types and predicting the type of each masked block. For each graph *G^i^*, 10% of blocks are randomly sampled, their block types are replaced with the special ‘mask’ block, and we denote these blocks as *B*. The atom nodes that correspond to each masked block are also replaced by a single atom node that is assigned the special ‘mask’ atom type so that the block’s atom composition is also masked. For a masked block *b* ∈ *B*, the probability vector over block types is predicted with 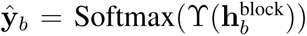, where Υ is a 2-layer MLP. We calculate the masked loss using a cross-entropy loss:

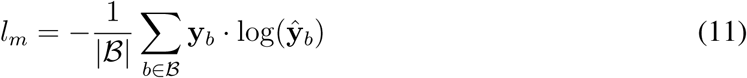

The pretraining loss is then calculated by a weighted sum of the above loss functions:

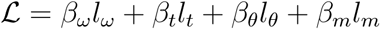

### 3 Training details for ATOMICA

#### Overview

We pretrain ATOMICA on the training split of biomolecular structures and small molecule structures to generate embeddings of molecular complexes at the atom, block, and graph scales. During pretraining, interaction complexes are sampled uniformly from the pooled dataset without stratifying by modality pair. To learn representations in a self-supervised manner, during training, we apply noise to the atomic coordinates and mask block types of the input graphs of the molecular complex. At inference time, embeddings from the graphs are generated without noise or masked blocks.

#### Pretraining dataset composition

CSD-derived data account for 85.8% of complexes, 58.6% of blocks, and 44.3% of atoms in the training split, while PDB-derived data account for 14.2%, 41.4%, and 55.7%, respectively.

#### Hyperparameter tuning

We employed a hyperparameter optimization strategy using Ray Tune [108] in conjunction with Optuna [109] and the Asynchronous Successive Halving Algorithm (ASHA) scheduler [110]. The hyperparameter space includes the number of nearest neighbors used to define graph edges *k* ∈ [4, **8**, 16], dropout in the tensor field network [**0**.**00**, 0.01, 0.05, 0.10], edge dimension *d*_edge_ ∈ [16, 24, **32**], node dimension *d*_node_ ∈ [16, 24, **32**], and the number of tensor field network layers *L* ∈ [**4**, 6, 8]. The best hyperparameters are shown in bold and chosen based on the lowest validation loss when trained on a random 10% subsample of the training set.

To determine the level of noise to apply to the interaction complexes, we conducted a second hyperparameter search on rotation *σ_ω_* ∈ [**0**.**25**, 0.5, 1], *ω*_max_ ∈ [0.25, **0**.**5**, 1], translation *σ_t_* ∈ [0.5, **1**, 1.5], and torsion *σ_θ_* ∈ [0.25, **0**.**5**, 1]. Again, the best hyperparameters are shown in bold and chosen based on the highest masked block-type prediction accuracy when trained on a random 10% subsample of the training set. For the loss function, we set *β_ω_* = 0.1*, β_t_* = 1*, β_θ_* = 1*, β_m_* = 1, and the block types are randomly masked with 10% probability. We train ATOMICA on the full training set with these hyperparameters. The learning rate cycles between 1e-4 and 1e-6 using Cosine Annealing Warm Restarts, with a cycle length of 400,000 steps, and training proceeds for 150 epochs.

#### Implementation

ATOMICA is implemented with PyTorch (Version 2.1.1) [111] and PyTorch Geometric (Version 2.1.1) [112]. Training runs were monitored with Weights and Biases [113]. The final ATOMICA model was trained on 4 NVIDIA H100 Tensor Core GPUs in parallel for 24 days.

### 4 Evaluation on RNA and protein annotation tasks

We evaluate ATOMICA on benchmark tasks for RNA and protein annotation. Here we describe how we adapt ATOMICA to each task and the baselines used for comparison.

#### 4.1 Datasets

We evaluate ATOMICA on five benchmark tasks from RNAglib [73] and MaSIF [16] spanning RNA and protein structure annotation: RNA–protein binding-site prediction, RNA Gene Ontology (GO) term prediction [77], RNA–small-molecule binding-site prediction, RNA–ligand pocket classification, and protein–ligand pocket classification.

##### RNAglib RNA–Protein

This task from RNAglib requires predicting whether each RNA residue is part of a protein binding site. The dataset contains 891 training structures with 52,175 residues of which 26.7% are part of a protein binding site, 191 validation structures with 11,063 residues of which 27.1% are part of a protein binding site, and 190 test structures with 10,851 residues of which 27.4% are part of a protein binding site. Positive labels are assigned to residues within 8 Å of any protein atom in a protein–RNA complex. This is a binary classification task at the residue level. Benchmark dataset splits are defined by clustering with USalign at a similarity threshold of 0.5.

##### RNAglib RNA-GO

This task involves predicting Gene Ontology (GO) terms associated with RNA molecules. The dataset includes 349 training samples, 75 validation samples, and 75 test samples. We predict membership in 5 GO term classes: GO:0000353 formation of quadruple SL/U4/U5/U6 snRNP (33.5%), GO:0010468 regulation of gene expression (20.0%), GO:0005682 U5 snRNP (16.2%), GO:0005688 U6 snRNP (15.6%), and GO:0005686 U2 snRNP (14.2%). This is a multilabel classification task where each RNA can be assigned to multiple GO terms. Of the 499 samples in this dataset, 161 samples are assigned no GO terms, 179 samples are assigned one GO term, and 159 samples are assigned two GO terms. Benchmark dataset splits follow a 60% sequence-identity split.

##### RNAglib RNA-Site

This task predicts whether each RNA residue is part of a small molecule binding site. The dataset contains 157 training structures with 10,092 residues of which 7.8% belong to a small molecule binding site, 34 validation structures with 2,162 residues of which 7.8% belong to a small molecule binding site, and 33 test structures with 2,158 residues of which 7.8% belong to a small molecule binding site. A residue is labeled positive if it lies within 8 Å of any atom of a bound ligand. This is a binary classification task at the residue level. Benchmark dataset splits are defined by clustering with USalign at a similarity threshold of 0.5.

##### RNAglib RNA–Ligand

This task predicts the ligand type for RNA binding pockets. The dataset includes 203 training pockets, 43 validation pockets, and 44 test pockets across 3 ligand classes: Paromomycin (PAR, 22.3%), Gentamycin C1A (LLL, 67.2%), and Aminoglycoside TC007 (8UZ, 10.0%). Pockets are defined by expanding from residues within 8 Å of ligand atoms. This is a multiclass classification task at the pocket level. Benchmark dataset splits are defined by clustering with USalign at a similarity threshold of 0.5.

##### MaSIF-ligand

This benchmark evaluates protein pocket classification across 7 common small molecule ligands: ADP (28.9%), CoA (12.6%), FAD (16.2%), heme (12.8%), NAD (11.4%), NAP (8.0%), and SAM (10.2%). The dataset contains 2,509 total pockets split into 1,839 training, 203 validation, and 467 test pockets. Binding pockets are defined as residues within 8Å of heavy atoms.of the ligand

#### 4.2 Evaluation of ATOMICA for prediction tasks

##### Representations and task heads

We use **z** as defined in Methods 2, with 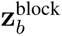 for the residue-level RNA–Protein and RNA–Site tasks and 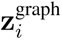 for RNA–GO and RNA–Ligand.

##### Pretraining-exclusion checkpoints and leakage control

For RNA–Protein, we use a checkpoint pretrained from scratch after excluding all protein–RNA complexes from the pretraining dataset. For RNA–Site and RNA–Ligand, we analogously use a checkpoint pretrained with all nucleic-acid–ligand complexes excluded. Consequently, the checkpoints reported for these three tasks have no direct pretraining exposure to the interaction type that defines the downstream labels, including homologous examples of that interaction type. RNA–GO uses the standard checkpoint because GO annotations are not inputs or objectives during pretraining. For every task, the down-stream model is fit only on the benchmark training split, hyperparameters and checkpoints are selected using the validation split, and the test split is reserved for final evaluation. Thus, no downstream test labels are used during either pretraining or downstream model fitting.

##### Frozen and fine-tuned evaluation

A result labeled ATOMICA uses frozen ATOMICA embeddings. For the frozen embeddings the ATOMICA model is fixed and its embeddings are passed to the same four-layer MLP protocol used for the frozen baseline embeddings below. A result labeled ATOMICA (fine-tuned) jointly updates ATOMICA and the task head. Fine-tuning is initialized from the same task-appropriate checkpoint used in the frozen comparison—the protein– RNA-exclusion checkpoint for RNA–Protein and the nucleic-acid–ligand-exclusion checkpoint for RNA–Site and RNA–Ligand. RNA–GO initializes from the standard checkpoint. Fig. 3 reports a fine-tuned ATOMICA only for RNA–GO.

##### RNA–GO fine-tuning

For the explicitly labeled fine-tuned RNA–GO model, we use a graph-level multi-label classifier for the five GO terms and focal loss with *γ* = 2.0, FL(*p_t_*) = −(1 − *p_t_*)*^γ^* log(*p_t_*). We use a four-layer MLP head, a constant learning rate of 4 × 10^−5^, weight decay of 1 × 10^−3^, gradient clipping at 1.0, and F1 macro for validation selection, and train for 200 epochs. During training, 10% of residues are randomly masked in 80% of samples. We train five random-seed replicates and ensemble their predictions by averaging logits.

##### MaSIF-ligand

For protein–ligand pocket classification, we use a pocket-level multiclass classifier to predict one of 7 ligand classes (ADP, COA, FAD, HEM, NAD, NAP, SAM). To address class imbalance, we use weighted cross-entropy loss with class weights inversely proportional to their training set frequencies. We pass 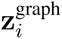 to a four-layer MLP trained for 300 epochs with a constant learning rate of 3 × 10^−5^, weight decay of 1 × 10^−3^, no gradient clipping, and F1 macro for validation selection. We train 5 models with different random seeds, identical architecture, and hyperparameters, and ensemble the models by taking the mean over the predicted logits.

#### 4.3 Benchmarking models for RNAglib tasks

We compare ATOMICA with structure- and sequence-based models on the RNAglib tasks. For every frozen pretrained encoder, including ATOMICA, we keep the encoder fixed and train the same task-specific four-layer MLP classifier on its embeddings. We sweep learning rate over {10^−3^, 5 × 10^−4^, 10^−4^, 5 × 10^−5^, 10^−5^}, dropout over {0.0, 0.1, 0.2, 0.3, 0.4}, and MLP hidden dimension over {64, 128, 256, 512, 1024}. For RNA–GO, we select between binary cross-entropy and focal loss on the validation set. For RNA–Ligand, we select between cross-entropy and weighted cross-entropy, with weights inverse to the training-label frequencies. We train five random-seed replicates and ensemble their predicted probabilities. Error bars and intervals are 95% percentile bootstrap confidence intervals over test examples using 2,000 resamples and macro metrics are resampled within class.

##### RNAglib

The RNAglib baseline is a graph neural network trained end-to-end on RNA 2.5D structure graphs, with nucleotides as nodes and edges given by backbone connectivity and base-pair interactions. We use the hyperparameters for the RGCN model specified by RNAglib [73].

##### RiNALMo, RNA-FM, and RNAErnie

We encode each RNA chain and extract residue-level embeddings from the final layer of each frozen pretrained model. For the graph-level RNA–GO and pocket-level RNA–Ligand tasks, we average the embeddings over residues in the evaluated sub-structure; for RNA–Protein and RNA–Site, we retain the residue-level embeddings. We then train the common MLP described above while keeping each RNA language-model backbone frozen.

##### gRNAde

Geometric RNA Design (gRNAde) [70] is a structure-based sequence-design model that applies graph vector-perceptron layers [114] to RNA backbone graphs. We pass each RNA structure through the frozen pretrained gRNAde encoder and extract residue-level embeddings, averaging them for RNA–GO and RNA–Ligand. We train the MLP while keeping the gRNAde encoder frozen.

##### RiNALMo parameter-efficient fine-tuning

For the bars labeled RiNALMo (PEFT), we adaptc RiNALMo using low-rank adaptation (LoRA) [79] on the attention projections in all 33 transformer blocks. Rank, learning rate, and loss are selected on the validation set. The selected rank is 16 for RNA–Protein, RNA–Site, and RNA–Ligand (4.5 million trainable parameters) and 32 for RNA–GO (9.0 million trainable parameters).

#### 4.4 Benchmarking models for MaSIF-ligand prediction tasks

We compare ATOMICA to geometry-based, protein language model (PLM), and contrastive-learning baselines on the MaSIF-ligand benchmark. For PLM baselines, we extract pocket embeddings by mean pooling the residue representations for pocket residues. We then train a four-layer MLP classifier on top of these embeddings using cross-entropy loss, and sweep hyperparameters over learning rate {10^−3^, 5 × 10^−4^, 10^−4^, 5 × 10^−5^, 10^−5^}, dropout {0.0, 0.1, 0.2, 0.3, 0.4}, and MLP hidden dimension {64, 128, 256, 512, 1024}. We report the performance when ensembling the predicted logits of models trained across 5 random seeds.

##### Pretraining-overlap control and late fusion

The frozen ATOMICA results for MaSIF-ligand in Fig. 3f–h use a checkpoint pretrained after excluding every protein–ligand item whose receptor is a MaSIF-ligand test receptor chain, shares an MMseqs2 cluster with a test receptor at 30% sequence identity and 80% coverage, or has Foldseek TM-score ≥ 0.5 to a test receptor. We apply the same filtering to the pretraining validation split and leave the other interaction types unchanged. For late fusion, we take the unweighted average of the class probabilities from the frozen ATOMICA model and ESM-2 (3B) without fitting an additional model.

##### MaSIF

MaSIF (Molecular Surface Interaction Fingerprinting) [16] is a geometry-based approach that characterizes protein surfaces using shape complementarity, electrostatics, hydrophobicity, and chemical features. We use the published MaSIF-ligand implementation.

##### Fpocket

fpocket [81] detects candidate cavities from protein geometry alone by Voronoi tessellation and alpha-sphere clustering, and reports hand-crafted descriptors for each detected cavity, including pocket volume, polar and apolar solvent-accessible surface area, alpha-sphere count, density and mean radius, apolar alpha-sphere proportion, and hydrophobicity, polarity, charge, and flexibility scores. We remove all seven benchmark ligands from each structure before running fpocket, so that no descriptor is computed from ligand atoms, and we describe the detected cavity with the largest number of atoms within 4 Å of the ligand. This supplies the binding-site location that every other baseline is also given, without supplying any ligand-derived feature. We concatenate the 19 cavity descriptors with the 20 residue-type counts of the cavity, z-score normalize features using training-set statistics, and use these features for the MLP.

##### ESM-2, ESM-C

We use ESM-2 (3B) [4] and ESM-C (600M) [115] to encode full protein chains and extract pocket residue embeddings from the final layer. We obtain pocket-level embeddings by mean pooling over pocket residues and pass them to the MLP.

##### ESM-3

We use ESM-3 (1.4B) [116] to encode protein chains using the sequence and structure tracks and extract pocket residue embeddings from the final layer. We obtain pocket-level embeddings by mean pooling over pocket residues and pass them to the MLP.

##### SaProt

SaProt (1.3B) [6] is a PLM that incorporates structural information via structure-aware tokenization. We extract residue embeddings from the pretrained SaProt model using Foldseek 3Di tokens for protein chains. We obtain pocket-level embeddings by mean pooling over pocket residues and pass them to the MLP.

##### ProstT5

ProstT5 [7] is a T5-based PLM trained on both sequence and structure. We encode protein chains with sequence tokens and with Foldseek 3Di structure tokens. We mean pool over pocket residues in each track, concatenate the resulting pocket embeddings, and pass the concatenated embedding to the MLP.

##### DrugCLIP

DrugCLIP [23] is a contrastive framework that learns aligned protein and smallmolecule representations. We extract pocket embeddings from the pretrained DrugCLIP protein encoder and feed them to the MLP. We also evaluated the original retrieval setup (encoding ligand conformers and selecting by highest cosine similarity to pocket embeddings), but it performed worse than the MLP.

##### DeeplyTough

DeeplyTough [117] learns pocket descriptors via metric learning on pocket similarity tasks. We extract pretrained pocket descriptors from DeeplyTough and classify them with an MLP.

##### PocketAnchor

PocketAnchor [118] learns pocket representations via contrastive learning by anchoring pockets to their ligands. We extract pretrained pocket embeddings from PocketAnchor and feed them to an MLP.

##### Label-transfer baselines and test-set similarity stratification

We search each of the test-pocket receptor chains against the training-pocket receptors with MMseqs2 at 30% sequence identity and 80% coverage, and with Foldseek. The two label-transfer baselines assign each test pocket the ligand class of its highest-scoring training receptor under the respective search, without a learned representation or a trained head. We stratify the test pockets by the Foldseek TM-score of the best alignment to a training receptor, giving 302 pockets at TM ≥ 0.8 and 165 below that threshold, and separately by whether an MMseqs2 hit exists at the above identity and coverage, giving 458 pockets with a hit and 9 without. Within each stratum we recompute macro-F1, micro-F1, macro-AUPRC and macro-AUROC from the test predictions. We also recompute the same metrics on cumulative subsets defined by a maximum Foldseek TM-score from 0.775 to 1.00 in steps of 0.025, where each subset contains every test pocket at or below that value and the largest subset is the full test set of 467 pockets. At a maximum Foldseek TM-score of 0.775, 138 pockets remain in the test set. We do not extend the sweep below a maximum Foldseek TM-score of 0.775 because subsets below this value do not contain all seven ligand classes, and macro-averaged metrics computed over different class sets are not comparable across subsets.

#### 4.5 ATP–ADP nucleotide-state discrimination

To test whether local pocket geometry captures nucleotide-associated conformational state rather than differences between unrelated proteins, we identify sequence-similar groups containing experimentally determined structures in both ATP- and ADP-bound states. The resulting benchmark contains 404 unique ligand-free pockets, including 223 ATP-associated and 181 ADP-associated pockets, from 120 PDB entries. We remove ATP or ADP, every other ligand, and waters before generating representations and define each pocket as the 50 protein residues nearest the bound ligand. For corresponding ATP- and ADP-bound structures, we retain only binding-site positions shared between the two structures that match in amino-acid identity. The 404 pockets are assigned to 60 MMseqs2 sequence clusters at 50% sequence identity and 80% coverage, each containing both nucleotide states, and these clusters define the units used for fivefold cluster-disjoint cross-validation.

ATOMICA uses the frozen 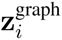 representation of each ligand-free pocket, whereas each protein language model embeds the full protein chain and averages the representations of the corresponding pocket residues. Each learned representation or geometric descriptor is evaluated through the same trained head, and no encoder is fine-tuned. Within each fold, we standardize features using the mean and standard deviation of the training pockets and remove columns whose training standard deviation is at most 10^−6^. We then train a four-layer MLP with ReLU activations and dropout using binary cross-entropy and Adam for 60 epochs at a batch size of 32. Hidden width ∈ {64, 256}, dropout ∈ {0.1, 0.3}, and learning rate ∈ {10^−4^, 10^−3^} are selected using a validation fold drawn from the training clusters rather than the test fold. Each configuration is fit with five independent seeds and the predicted probabilities are averaged.

We pool the out-of-fold probabilities across the five cross-validation folds and calculate ATP– ADP AUROC separately within each sequence cluster. We report the unweighted mean across clusters, such that each cluster contributes equally irrespective of the number of pockets it contains. The random reference is 0.500. We report 95% confidence intervals from 2,000 percentile bootstrap resamples of sequence clusters.

For the metal-concordant analysis, we restrict the benchmark to ATP- and ADP-associated pockets whose corresponding structures have the same metal-ion state within 4 Å of the nucleotide. The metal-free analysis further requires the corresponding structures to lack a metal ion within 4 Å of the nucleotide. The held-out analysis requires the evaluated structures to be outside the ATOMICA pretraining set. Within each subset, cluster-macro AUROC is calculated only for sequence clusters containing both nucleotide states.

#### 4.6 Same-ligand pocket retrieval

We construct this benchmark from the held-out protein–ligand test split, and define each pocket as the 50 protein residues nearest the bound ligand, which is removed before representation generation. Given a query, we rank candidate pockets by representation similarity. A candidate is relevant if it binds the query’s ligand and has no detectable Foldseek structural alignment to the query. A same-ligand pocket that does have an alignment is removed from that query’s ranking rather than counted against the representation, since it is a true same-ligand pair and scoring it as a negative would penalize a representation for recognizing the site. Every pocket is used as a query, and the 428 that retain at least one relevant partner under this rule form the query set. The benchmark contains 892 pockets, 428 queries, 44 ligand classes, and 2,349 positive query–reference pairs.

For the protein language models, we embed the full protein and average the representations of the pocket residues, while ATOMICA uses the frozen 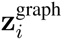 representation of each ligand-free pocket. We report mean average precision (mAP), normalized discounted cumulative gain (nDCG), and R-precision in the main text and Supplementary Table S2.

##### Macro averaging over sequence clusters

All reported values are macro-averaged over the 30% sequence identity MMseqs2 clusters used to construct the split. The 428 queries fall into 205 clusters, so one cluster can contribute several near-duplicate query pockets and a plain average over queries would weight those clusters more heavily than clusters represented by a single pocket. The exact-permutation chance value is aggregated under the same weighting. The 95% confidence intervals for each representation are obtained by bootstrap resampling over the 30%-sequence-identity clusters.

##### Sensitivity to the Foldseek TM threshold of relevant pairs

The benchmark places a single restriction on a pair, the requirement of no detectable Foldseek alignment. To characterize how the comparison depends on it, we set up different thresholds for relevant pockets up to a maximum Foldseek TM of 0, 0.1, 0.2, 0.3, 0.4, 0.5, 0.6, 0.7, 0.8 and 1, and rescore every representation at each value (Fig. S4h) with Foldseek TM of 0 matching the values reported in Fig. 3k.

### 5 Cross-modality comparison for orthosteric PPI inhibitors

#### 5.1 Dataset

We used the 2P2Idb database, which contains experimentally determined structures of protein– protein interactions and their inhibitors from the Protein Data Bank. PPI structures were processed to retain the target protein A and partner B, while protein–inhibitor structures were processed to retain protein A and inhibitor I. The final analysis contains 26 protein–peptide complexes with 965 matched protein–inhibitor structures and six protein–protein complexes with 187 matched protein– inhibitor structures.

#### 5.2 Protein–peptide inhibitor analyses

For protein–peptide complexes where peptide B contains ≤ 30 residues, we performed block-level comparisons between inhibitor-bound structures and peptide B to assess whether ATOMICA embeddings capture structural similarities at the interface.

##### 5.2.1 Structure alignment

We aligned inhibitor target chain structures to PPI target chain structures. We first aligned the sequences using the BLOSUM62 substitution matrix with a gap opening penalty of −10 and a gap extension penalty of −1, then superposed the matched C*α* atom coordinates using the Kabsch algorithm. Iterative outlier rejection was then applied and residue pairs with RMSD *>* 2.0 Å were removed. The alignment was refined over 5 cycles until convergence.

##### 5.2.2 ATOMICA embedding computation

We computed ATOMICA embeddings for both inhibitors and peptide B using the pretrained ATOMICA model. For the inhibitor embedding, we embedded the inhibitor bound to the target protein pocket and extracted the block embeddings of the inhibitor, 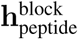. For the protein– peptide embedding, we embedded the peptide as an isolated chain and extracted the block embeddings of the peptide, 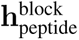.

##### 5.2.3 Block pairwise comparison

For each inhibitor-PPI pair, we performed the following block-level comparison. We applied the rotation matrix *R* and translation vector *t* from the alignment step to transform the inhibitor block coordinates into the PPI reference frame. Block coordinates, **x**, are the average of the atom coordinates within each block. We filtered out global-type blocks and single-atom inhibitor blocks, retaining only blocks with more than one atom. Pairwise embedding distances, 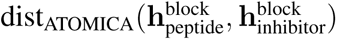, between all inhibitor blocks and peptide B blocks were calculated as the cosine distance between block embedding vectors. Pairwise spatial distance was defined as the Euclidean distance between block centers, 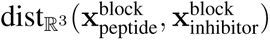. We retained matched structures with at least ten inhibitor– peptide block pairs, resulting in 26 protein–peptide complexes and 965 matched protein–inhibitor structures. The number of inhibitor structures matched to each PPI complex is provided in Table S3.

##### 5.2.4 Retrieval analysis

For each matched protein–inhibitor structure, we ranked all inhibitor–peptide block pairs by embedding distance and selected the ten pairs with the lowest distance. Pairs with 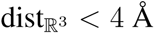 were considered spatially close. Precision@10 is the fraction of the ten retrieved pairs that are spatially close, and Fold Change@10 is Precision@10 divided by the fraction of all block pairs that are spatially close.

#### 5.3 Protein–protein inhibitor analyses

For protein–protein complexes where protein B has more than 30 residues, we used surface sampling to test whether ATOMICA embeddings identify patches near the inhibited A–B interface. We retained the six complexes with a binary A–B structure and excluded the INTEGRASE/LEDGF and TNFA oligomeric structures. The number of matched inhibitor structures for each complex is provided in Table S4.

##### 5.3.1 Surface point sampling

We generated molecular surface representations and sampled points uniformly across the surface of each protein B. We used the MSMS tool [119] to compute solvent-excluded surfaces with a vertex density of 3.0 vertices per Å^2^ and a probe radius of 1.5 Å. For each protein B chain, we extracted atomic coordinates and generated a triangular mesh with vertices, faces, and surface normals. We then sampled 1,000 points per protein B surface using area-weighted triangle sampling. For each sampled triangle, we uniformly sampled a point using barycentric coordinates. For each point, we drew *r*_1_*, r*_2_ *∼ U* (0, 1), computed the barycentric coordinates 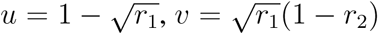, and 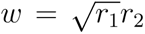, and set the point position to **p** = *u***v**_0_ + *v***v**_1_ + *w***v**_2_, where **v**_0_, **v**_1_, and **v**_2_ are the triangle vertices.

##### 5.3.2 Interface patch definition

For each sampled surface point, we defined a local interface patch based on spatial proximity to protein residues. We computed the Euclidean distance from the surface point to all protein B residue centers. We selected the residues within 16.0 Å of the surface point and discarded points with fewer than eight nearby residues. For each interface patch center on protein B, we computed the distance to the nearest C*α* atom on protein A.

##### 5.3.3 ATOMICA embedding computation

###### Inhibitor embeddings

For each protein–inhibitor complex (*A*–*I*), we applied ATOMICA to the inhibitor bound to the target protein pocket and used the model’s learned attention-pooled representation over the contextualized inhibitor blocks to obtain 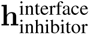. The attention pooling is learned during self-supervised pretraining and is not fitted on this retrieval task.

###### Interface patch embeddings

For each protein–protein complex (*A*–*B*), we defined each local surface patch as the protein B blocks within 16 Å of a sampled point. We applied ATOMICA to each patch and used the same learned *h*^interface^ representation to obtain 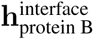. These patch embeddings serve as retrieval candidates. We evaluate two alternative interface representations in Supplementary Table S7, an unweighted mean over the block representations and **z**^interface^ of Methods 2, both of which reach lower Fold Change@10 than the *h*^interface^ representation used here.

##### 5.3.4 Retrieval analysis

We formulate retrieval as follows: given an inhibitor embedding (query), we rank sampled interface patches on protein *B* (candidates) by embedding similarity and evaluate whether high-similarity patches localize to the native *A*–*B* binding site.

To prevent direct access to the native A–B interface, the query representation pools only contextualized inhibitor ligand blocks from the A–I complex and the candidates are encoded separately using only patches on protein B. Proteins A and B are never jointly embedded, and no protein A block representation enters the similarity calculation. The experimental coordinates of protein A are accessed only after ranking to label the proximity of each patch to the native A–B interface.

We compute the following distance metrics: embedding distance 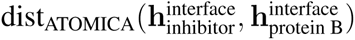, given by the cosine distance between inhibitor embedding and interface patch embedding, and spatial distance 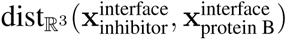, given by the Euclidean distance from the interface patch center to the nearest C*α* atom on protein A.

###### Retrieval procedure

For each inhibitor, we ranked all interface patches by dist_ATOMICA_ (ascending). The ten patches with the lowest dist_ATOMICA_ were selected. Patches within 12.0 Å of protein A at the native interface were considered “close.” We selected 12.0 Å because it is the 25th percentile of the distances from sampled patches to the PPI interface. We computed *Precision@10*, the fraction of the top 10 patches within the geometric threshold, and *Fold Change@10*, Precision@10 divided by the baseline rate. The baseline rate is given by the proportion of all patches on protein B that are within 12.0 Å of the A–B interface.

#### 5.4 AlphaFold3 representation comparison

We ran AlphaFold3 v3.0.1 [1], saved the embeddings, and extracted the post-recycling Pairformer trunk state, which contains a 384-dimensional single representation for each token and a 128-dimensional pair representation for each token pair. For each complex A–B, protein A was co-folded with its inhibitor to provide the ligand query context, while protein B was folded alone to provide the peptide blocks or protein surface patches used as retrieval candidates. The inhibitor and protein B were therefore never included in the same AF3 input. Outside representation construction, the experimental coordinates of protein A were used only to align structures for the block task or to label patch proximity after ranking.

For the AF3 single representation, we averaged the token representations belonging to each inhibitor block, peptide block, or protein B patch. Table 1 defines the three AF3 pair pooling schemes and the row and column tokens used for each object. Individual block descriptors were used for protein–peptide retrieval, while the inhibitor block descriptors were averaged with equal weight to form each protein–protein interface query. We then used cosine distance and the same retrieval candidates, geometric labels, and Fold Change@10 calculation as for ATOMICA.

**Table 1.** AF3 pair pooling schemes. For each protein B patch, inhibitor, peptide block, or inhibitor block, the AF3 pair representations are pooled over the corresponding row tokens and the specified column tokens. The three schemes differ in which column tokens are included.

| AF3 pair pooling scheme | Protein–protein complexes |  | Protein–peptide complexes |  |
| --- | --- | --- | --- | --- |
| | $\mathbf{h}_{\text{protein B}}^{\text{interface}, i}$ | $\mathbf{h}_{\text{inhibitor}}^{\text{interface}}$ | $\mathbf{h}_{\text{peptide}}^{\text{block}}$ | $\mathbf{h}_{\text{inhibitor}}^{\text{block}}$ |
| Self-pair only | $\frac{1}{\vert T_{\text{patch}}\vert } \sum_{s \in T_{\text{patch}}} \mathbf{p}_{ss}$ | $\frac{1}{\vert I\vert } \sum_{b \in I} \frac{1}{\vert T_b\vert } \sum_{s \in T_b} \mathbf{p}_{ss}$ | $\frac{1}{\vert T_b\vert } \sum_{s \in T_b} \mathbf{p}_{ss}$ | $\frac{1}{\vert T_b\vert } \sum_{s \in T_b} \mathbf{p}_{ss}$ |
| Own sub-matrix | $\frac{1}{\vert T_{\text{patch}}\vert ^2} \sum_{s \in T_{\text{patch}}} \sum_{t \in T_{\text{patch}}} \mathbf{p}_{st}$ | $\frac{1}{\vert I\vert } \sum_{b \in I} \frac{1}{\vert T_b\vert \vert T_I\vert } \sum_{s \in T_b} \sum_{t \in T_I} \mathbf{p}_{st}$ | $\frac{1}{\vert T_b\vert \vert T_{\text{peptide}}\vert } \sum_{s \in T_b} \sum_{t \in T_{\text{peptide}}} \mathbf{p}_{st}$ | $\frac{1}{\vert T_b\vert \vert T_I\vert } \sum_{s \in T_b} \sum_{t \in T_I} \mathbf{p}_{st}$ |
| All tokens | $\frac{1}{\vert T_{\text{patch}}\vert \vert T_B\vert } \sum_{s \in T_{\text{patch}}} \sum_{t \in T_B} \mathbf{p}_{st}$ | $\frac{1}{\vert I\vert } \sum_{b \in I} \frac{1}{\vert T_b\vert \vert T_A \cup T_I\vert } \sum_{s \in T_b} \sum_{t \in T_A \cup T_I} \mathbf{p}_{st}$ | $\frac{1}{\vert T_b\vert \vert T_{\text{peptide}}\vert } \sum_{s \in T_b} \sum_{t \in T_{\text{peptide}}} \mathbf{p}_{st}$ | $\frac{1}{\vert T_b\vert \vert T_A \cup T_I\vert } \sum_{s \in T_b} \sum_{t \in T_A \cup T_I} \mathbf{p}_{st}$ |

**Notation.** $s$ and $t$ index AF3 tokens, and $\mathbf{p}_{st}$ denotes the AF3 pair representation for row token $s$ and column token $t$ . $T_{\text{patch}}$ denotes the AF3 tokens in the sampled surface patch on protein B, $T_B$ denotes all AF3 tokens in protein B, and $T_{\text{peptide}}$ denotes all AF3 tokens in the peptide. $b$ denotes a block and $T_b$ its AF3 tokens, $I$ denotes the set of inhibitor blocks and $T_I$ all inhibitor tokens, and $T_A$ denotes all AF3 tokens in protein A.

#### 5.5 Complex- and superfamily-level aggregation

For both representation models and both retrieval tasks, Fold Change@10 was first computed for each matched inhibitor query and then averaged within each complex. Spearman correlation between embedding and spatial distance was retained as a descriptive complex-level statistic. We assigned complexes using the 2P2Idb SuperFamily annotation and took an unweighted mean over the constituent complex-level values, yielding 14 protein–peptide and five protein–protein superfamilies. Each superfamily therefore contributes one value irrespective of its number of complexes, inhibitor structures, block pairs, or sampled patches. No across-system significance claim is based on these nested observations.

### 6 Analysis of dark proteome with ATOMICA-Ligand

#### Overview

We demonstrate the versatility of ATOMICA by fine-tuning it to assign ions and ligands to binding sites. The fine-tuned version of the model is applied to putative binding sites in the dark proteome.

#### Training ATOMICA-Ligand

The objective is to predict the probability of a specific ion or ligand binding to a given protein interface pocket. We frame this as a binary prediction task and finetune a separate model for each ion and small molecule. A predictive head is a *L*_ligand_-layer MLP. For each ion and small molecule, we use Ray Tune with Optuna and ASHA to fine-tune ATOM-ICA-Ligand from ATOMICA and find the optimal hyperparameters among *L*_ligand_ ∈ [3, 4, 5], learning rate ∈ [10^−6^, 10^−3^], non-linearity ∈ [`relu, gelu, elu`], hidden dimension of MLP [16, 32, 64], gradient clipping ∈ [`None`, 1], and the number of nearest neighbors used to define graph edges *k∈* [4, 6, 8]. To address class imbalances in our dataset, we apply a weighted sampling strategy during training, where each protein pocket receives a sampling weight inversely proportional to the total count of its label class. For each ion and small molecule, we fine-tune ATOMICA-Ligand for 50 epochs on 1 NVIDIA H100 Tensor Core GPU. Three replicate models are trained for each ion and small molecule. For binary classification of binding sites, we set thresholds that maximize the F1 score, constraining these values to fall within the range of 0.05 to 0.95. We use a one-vs-rest binary formulation for ATOMICA-Ligand because it yields ligand-specific decision boundaries.

#### Dataset curation

For each ion or small molecule, we collect all graphs in the pretraining set in which that ligand is bound to a protein. We cluster protein binders at a 30% protein sequence identity cutoff, coverage of 80%, sensitivity of 8, and cluster mode 1 using MMseqs2 [100]. The clusters are then divided into training, validation, and test sets in an 8:1:1 ratio. We set up this split for the following metal ions: Ca, Co, Cu, Fe, K, Mg, Mn, Na, Zn, and the following small molecules with these PDB chemical codes: ADP (adenosine diphosphate), ATP (adenosine triphosphate), CIT (citric acid), CLA (chlorophyll A), FAD (flavin adenine dinucleotide), GDP (guanosine diphosphate), GTP (guanosine triphosphate), HEC (heme C), HEM (heme B), NAD (nicotinamide adenine dinucleotide), NAP (NADP+, nicotinamide adenine dinucleotide phosphate, oxidized form), NDP (NADPH, nicotinamide adenine dinucleotide phosphate, reduced form).

#### Dark proteome annotation

We define dark proteins using Foldseek clusters from the AlphaFold Protein Structure Database with `isDark == 1`, indicating that the cluster lacks functional annotation [45]. We limit our analysis to the 33,482 clusters with an average pLDDT *>* 90. For each cluster, we take the representative protein and run PeSTo [93] on the protein structure to predict ion and small molecule binding sites. We keep residues with PeSTo binding score *>* 0.8 as the putative binding site, with a minimum of 5 residues required. In total, we extract 2,851 ion binding proteins and 969 small molecule binding proteins from the 33,482 representative proteins. Given these binding interfaces, we run all fine-tuned ATOMICA-Ligand ion and small molecule models to assign chemical identities to the binding sites. To assess cross-model concordance, we fold ATOMICA-Ligand predicted protein–ligand complexes with AlphaFold3 [1] and score each prediction both globally and against the PeSTo pocket that ATOMICA-Ligand was given a priori. Let *L* denote the ligand tokens and *P* the pocket tokens, let *e_ij_* be the AlphaFold3 predicted aligned error, which is the expected error in token *j* when the structure is aligned on token *i*, and let *c_ij_* be the corresponding contact probability. We record the global ipTM, the mean pLDDT over ligand atoms, the largest contact probability max*_i∈P,j∈L_ c_ij_*, and the mean of the symmetrized error (*e_ij_* + *e_ji_*)*/*2 over *i ∈ P* and *j ∈ L*. Following AlphaFold3’s contact-probability-weighted aggregation of PAE, we calculated a pocket-restricted contact-weighted PAE by restricting the pairwise PAE and contact-probability matrices to the PeSTo-predicted pocket and ligand tokens,

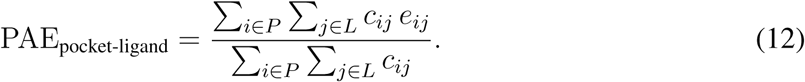

We take PAE_pocket-ligand_ as the primary measure because weighting by contact probability ensures that the residues that AlphaFold3 places against the ligand are included in the average.

ATOMICA-Ligand annotates 2,978 ion and 85 small molecule pocket-ligand pairs as positive, and we fold each of these. For comparison, we sample the same number of random control complexes from dark proteome proteins with predicted ion and small molecule binding sites, matched to the annotated complexes in both the number and the identity of the ligands. This control is drawn without reference to the ATOMICA-Ligand prediction and therefore still contains pairs that ATOMICA-Ligand calls positive, so we define a stricter predicted-negative control as the subset of the random control that ATOMICA-Ligand scores below its ligand-specific threshold.

To assess sequence similarity to the classifier training data, we search the 969 screened proteins against the 2,807 HEM and 1,933 HEC training-positive receptor chains using MMseqs2 [100] and phmmer. We additionally run three-iteration jackhmmer [120] against the full 154,400-chain classifier training split and identify recovered HEM or HEC training positives. We define a significant hit as one with *E ≤* 10^−3^. We define coverage-weighted identity as the number of identical aligned residues divided by the longer full sequence and compute its Spearman rank correlation with the corresponding ATOMICA-Ligand score. We stratify the cluster-disjoint HEM and HEC held-out test splits using MMseqs2 alignments to same-cofactor training positives, with the no-training-alignment stratum containing test pockets without a hit that covers at least 50% of both sequences. Held-out scores are averaged over the three classifier replicates, and lift is the AUPRC divided by the class prevalence within each stratum.

For sequence-based annotation we run the Google Colab notebook with ProtNLM [121].

### 7 Quantifying the importance of interface blocks with ATOMICASCORE

#### Definition

ATOMICASCORE measures how much the representation of an interaction complex depends on each of the blocks at its interface. For block *i* of the interaction graph *G*, we construct *G* \ *i* by replacing block *i* with the special mask block and substituting its constituent atoms with a single special mask atom, and define

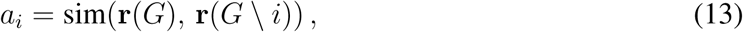

where sim denotes the cosine similarity and **r**(·) is a readout of the interaction graph. A low *a_i_* indicates a large change in the readout and therefore high importance for block *i*. We take **r** to be the component-normalized mean of 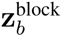 over the blocks of the ligand molecule, following Methods 2, and we mask and evaluate only amino-acid residue blocks.

#### Evaluation

We annotate protein–small-molecule complexes of the held-out test set with PLIP [65]. A residue is labeled positive when it participates in a PLIP-annotated non-covalent interaction with the ligand. These annotations comprise hydrogen bonds, hydrophobic interactions, *π*-stacking, metal complexes, and halogen bonds. We keep the 5,497 complexes with at least 12 scored residues, at least one annotated residue, and at least one residue of each label. Residues are ranked by ascending ATOMICASCORE, and we report the fraction of the 10 top-ranked residues involved in an annotated non-covalent interaction as precision@10, the area under the ROC curve over all scored residues of a complex, and the partial Spearman correlation between ATOMICASCORE and the label given the distance from the residue to the nearest ligand atom. Each quantity is computed per complex and averaged over complexes, with 95% percentile bootstrap confidence intervals resampled over complexes.

#### Baselines

For reference we select 10 interface residues at random. We compare ATOMICAS-CORE to ESM-2 (3B) [4], scoring each amino acid by the log likelihood of the masked mutant relative to the native residue [66], and to LigandMPNN [67], which conditions on the non-protein atoms of the complex and is therefore given the same interface geometry as ATOMICA. LigandMPNN is read out in two ways: (1) the masked-marginal log likelihood of the native residue given the backbone and the ligand, which is analogous to how we obtain scores from ESM-2, and (2) the difference between that quantity and the same one computed with the ligand context removed, which is a leave-the-ligand-out perturbation and the closest published analogue of the leave-the-block-out construction of ATOMICASCORE. Both are computed on the full complex with the native identity of every other residue visible to the model, and readout (2) has higher precision@10 and AUROC and is reported in Fig. 2.

### 8 Masked block-type prediction and sequence recovery

#### Training ATOMICA on a single pair of interacting modalities (modality-specific training)

To test whether the representations learned by ATOMICA generalize across modalities, we train models with identical architecture and hyperparameters on single pairs of interacting modalities (small molecules, protein–ion, protein–small-molecule, protein–DNA, protein–RNA, protein– peptide, protein–protein, nucleic acid–small-molecule). These models use the ATOMICA training setup and training data, filtered to a single pair of interacting modalities. The models are trained for 150 epochs on 4 NVIDIA H100 Tensor Core GPUs in parallel. The model checkpoint with the lowest validation loss is then used for further fine-tuning on masked block-type prediction on the same training data for 50 epochs with a learning rate of 1e-4. We also fine-tune ATOMICA for 50 epochs on masked block-type prediction for each pair of interacting modalities. To compare the quality of embeddings generated by ATOMICA and versions of it trained on single modalities, we evaluate the accuracy of masked block-type prediction on a test set. This test set was not seen by any of the models and has at most 30% sequence identity and minimal small molecule fingerprint similarity to any training or validation data.

Because block definitions vary across modalities, masked block-type prediction is interpreted through within-modality comparisons between ATOMICA and the modality-specific models rather than through absolute values across modalities.

Macro-AUPRC is the mean of one-versus-rest average precision over the block classes present, computed within each of the five masking seeds and then averaged across seeds. We report a 95% confidence interval from 1,000 bootstrap replicates that resample the test structures, drawing one resample per replicate and applying it to all five seeds. Two properties of this metric shape how the interval is formed. Resampling structures changes which classes are present, and the classes lost are the rare ones with the lowest average precision, so each replicate is scored against the same class set it is computed on and the change in class composition cancels. Average precision also rises when a structure is drawn more than once, because a duplicated positive both scores higher and counts twice, and the effect is largest for classes with few instances. We therefore take the dispersion from the bootstrap and center the interval on the estimate, reporting the value plus or minus 1.96 standard deviations of the bootstrap distribution.

Most small-molecule–small-molecule complexes from the CSD contain two conformers of the same compound with identical block-type sequences. Masking a block in only one conformer can therefore allow the model to recover its identity from the corresponding block in the other conformer. To prevent this shortcut, we evaluate held-out CSD graphs with identical, index-aligned block-type sequences and mask position *k* in both conformers simultaneously. We apply this masking rule to both the ATOMICA and small-molecule-only checkpoints over five masking seeds and report the mean across seeds with the same class-level confidence interval. Macro-AUPRC is averaged one-versus-rest over the block classes represented in each seed. An average of 179 classes is represented in the paired-masking evaluation, giving a chance level of 1*/*179 = 0.0056.

To evaluate the contribution of CSD pretraining to common chemical motif recovery, we use the held-out protein–small-molecule test split and compare the full checkpoint pretrained on PDB and CSD complexes with an architecture-matched checkpoint pretrained without the CSD. Both checkpoints contain the 436-class masked block-type prediction head learned during pretraining and are evaluated without retraining or fine-tuning. For the primary evaluation, we consider only ligand-side blocks assigned to the vocabulary of 290 common chemical motifs and mask each eligible block once in a separate forward pass while leaving every other block in the complex unmasked. Single-atom ligand blocks, whose labels represent elements rather than entries in the common chemical motif vocabulary, are not included in the primary common chemical motif analysis. We report top-1 accuracy across common chemical motifs and macro-AUPRC, computed as the mean one-vs-rest average precision across observed common chemical motif classes.

#### Amino-acid sequence recovery

We use the multimodal pretrained ATOMICA model for all six protein-containing interaction types. Masked block-type prediction is one of the pretraining objectives, so the pretrained model performs this task without task-specific fine-tuning, and no fine-tuning was performed for this evaluation. We measure top-1 amino-acid recovery after simultaneously masking 10%, 30%, or 50% of the amino-acid positions in each interaction interface. All models are scored on identical positions over five mask seeds on the held-out 30%-sequence-identity test splits. We report the mean over seeds together with a 95% confidence interval obtained by resampling test structures with replacement over 2,000 bootstrap replicates, which accounts for the dependence between positions within one structure. ATOMICA operates on the interaction-interface patch, with predictions restricted to its 20 amino-acid logits. ProteinMPNN [68] and LigandMPNN [67] receive the complete parent complex, with only the selected positions redesignable and every other protein residue fixed to its native identity. For ProteinMPNN and LigandMPNN, we decode one sequence at *T* = 10^−6^. For ProteinMPNN, we use the `v_48_020` and `v_48_002` checkpoints. We evaluate LigandMPNN with and without fixed-residue side-chain context before retaining the configuration with the highest recovery for each interaction type. ESM-2 (3B) [4], ESM-C (600M) [115], and ESM-3 [116] receive the full sequence of the chain containing the masked positions. SaProt [6] and ProstT5 [7] additionally receive that chain’s structure as Foldseek 3Di tokens [80]. All selected positions are masked simultaneously and recovered by argmax over the 20 amino-acid probabilities. The released baseline models cannot be retrained on the exact ATOMICA pretraining split, and their PDB or UniRef training datasets overlap the test set.

#### Nucleotide sequence recovery

We report ATOMICA (fine-tuned) for this evaluation, using the 50-epoch per-modality fine-tuning protocol described in Methods 8 for protein–RNA, protein–DNA, and RNA–ligand interfaces, with nucleotide positions additionally upweighted in the masked block-type loss at 10% masking. Nucleotide positions are upweighted in the masked block-type loss by a factor of 20 for protein–RNA and protein–DNA and 5 for RNA–ligand, selected on validation nucleotide recovery. Around 70% of maskable blocks at these interfaces are amino acids or common chemical motifs, so the unweighted pretraining objective assigns most of its gradient to positions that this evaluation does not score. We evaluate top-1 recovery after simultaneously masking 10%, 30%, or 50% of the nucleotide interface positions, restricting ATOMICA predictions to the four canonical nucleotide logits. All methods are scored on identical positions over five mask seeds on the same held-out 30%-sequence-identity test splits, and we report the mean over seeds with a 95% confidence interval from the same structure-level bootstrap used for amino-acid recovery. NA-MPNN [69] receives all protein and nucleic-acid polymer chains from the parent complex, but does not encode small-molecule atoms. The selected nucleotide positions are jointly redesigned in one autoregressive decode at *T* = 10^−6^, with all other polymer residues fixed to their native identities. gRNAde [70] and RhoDesign [71] receive neither the interacting protein nor ligand and are therefore not evaluated on protein–DNA interfaces. For gRNAde, we recover each evaluated position by argmax from RNA structure without native sequence context. For RhoDesign, we obtain argmax predictions in a teacher-forced autoregressive pass conditioned on the RNA structure and native preceding residues. RhoDesign retains all evaluated positions and fills the remaining spatial context with their nearest RNA residues.

### 9 Additional quality metrics and statistical analyses

#### Visualization of embeddings

We embed all molecular interaction complexes from training, validation, and testing datasets, and project these embeddings into two dimensions using Uniform Manifold Approximation and Projection for Dimension Reduction (UMAP) [62] with 30 neighbors and a minimum distance of 0.001. We visualize all interaction complexes with PyMOL [122].

#### Visualization of atom and block type

From molecular interaction complexes in training, validation, and testing datasets, we take the element-wise average atom and block embeddings for each element and block type. We project the atom and block embeddings into two dimensions using Principal Component Analysis (PCA). Elemental categories and properties of amino acids and nucleotides are assigned post hoc for visualization and are not provided as model input features.

#### Metal-coordination linear probes

We matched the metal sites in the protein–ion splits to MetalPDB records [63] using the metal element, PDB residue identifiers, and a rotation- and translation-invariant fingerprint of metal–donor distances, retaining only unique matches after excluding conflicting or ambiguous assignments. The full coordination-number target counts every donor in the deposited MetalPDB coordination sphere, including non-protein donors that are absent from the model input, while the protein-donor coordination number counts only donors from protein residues present in the input graph. We use the polyhedron assignments distributed by MetalPDB and generated with FindGeo [64], retaining geometries with vacancies as distinct labels and grouping less frequent polyhedra into an other class to obtain 14 classes after excluding irregular or unassigned sites. This procedure yields verified coordination-number labels for 3,654 of the 7,200 held-out test sites and FindGeo geometry labels for 2,313 of these sites.

We freeze the pretrained backbone and represent each metal-ion block *b* with **z**^block^ defined in Methods 2. We fit the same head to a one-hot encoding of the metal element as an element-only control. The coordination-number tasks use eight labels comprising coordination numbers one through seven and a class for eight or more, with 20,159 training sites, 2,421 validation sites, and 3,654 test sites, while the geometry task uses 13,839 training sites, 1,535 validation sites, and 2,313 test sites. We standardize features using the training-set statistics and fit an *L*_2_-regularized multinomial logistic regression without hidden layers, selecting the inverse regularization strength from 0.003, 0.01, 0.03, 0.1, 0.3, 1.0 by validation balanced accuracy before evaluating the held-out test set. The fixed-coordination-number analysis fits a separate probe after restricting the data to one deposited coordination number and retaining geometry classes represented by at least ten test sites. We use balanced accuracy as the primary metric because the classes are imbalanced, and we report 95% percentile confidence intervals from 2,000 bootstrap resamples of PDB entries so that all sites from the same structure remain grouped.

### 10 Experimental evaluation of selected heme candidates

#### 10.1 Selection of heme candidates for experimental evaluation

From the dark proteome, 59 proteins are predicted to bind heme non-covalently (HEM ligand) and eight are predicted to bind heme covalently (HEC ligand), with two proteins carrying both predictions, giving 65 distinct heme candidates. Nine of these were selected for experimental evaluation. In addition to their ATOMICA-Ligand score, they were selected based on surpassing a minimum AlphaFold3 ipTM score of 0.85 and the absence of a large transmembrane domain, which would make protein synthesis difficult. The UniProt [94] identifiers of these proteins are A0A7W1B5T5, A0A1T4N4K0, A0A4P5TA35, A0A7W0X6V6, A0A2V6P8N7, A0A136KY61, A0A7Y8LED7, A0A7V7N0X5, and V5BF69.

#### 10.2 Synthesis of proteins

A0A7W1B5T5 and A0A1T4N4K0 were synthesized with automated flow peptide synthesis. In addition, all nine proteins were synthesized and purified as Strep-tagged recombinant proteins by GenScript (Piscataway, NJ, USA). Chemical characterization of the two synthesized proteins is given in Figs. S10 (A0A7W1B5T5) and S11 (A0A1T4N4K0).

##### Reagents and supplies

Disposable fritted polypropylene reaction vessels were purchased from Torviq (cat. no. SF-0500-LL for 6 mL reactors). H-RAM-ChemMatrix resin (0.17 mmol/g, batch no. 18007556) was purchased from ChemMatrix. Fmoc-amino acids were purchased from the Novabiochem product line of Millipore Sigma and were used as received. OmniSolv® grade *N,N*-Dimethylformamide (DMF, biosynthesis grade) was purchased from Millipore Sigma (product DX1732-1) and treated with AldraAmine trapping agents (for 1000–4000 mL DMF, catalog number Z511706). *N,N*-Diisopropylethylamine (DIPEA; ReagentPlus ≥ 99%), piperidine (ACS reagent, ≥ 99.0%), trifluoroacetic acid (HPLC grade,≥ 99.0%), triisopropylsilane (TIPS, ≥ 98.0%), acetonitrile (ACN, HPLC grade), 1,2-ethanedithiol (EDT, 98+%), trimethylsilyl chloride (TMS-Cl, 98.0% GC), triphenylphosphine (PPh3, ReagentPlus 99%), and formic acid (FA, ≥ 95.0%) were all purchased from Sigma-Aldrich and used as received. O-(7-azabenzotriazol-1-yl)-*N,N,N’,N’*-tetramethyluronium hexafluorophosphate (HATU, ≥97.0%) was purchased from P3 Biosystems. Dichloromethane (DCM, HPLC grade *>* 99.9%) was purchased from Fisher. Hemin (BioXtra, ≥96.0%, cat. no. 51280-1G) was purchased from Sigma-Aldrich. Water was deionized in-house using a Milli-Q water purification system from Millipore. Bovine serum albumin (BSA, protease-free, cat. no. A7030), casein (technical grade, cat. no. C7078), and lysozyme (chicken egg white, cat. no. L6876) were purchased from Sigma-Aldrich and used as negative controls in the heme-binding experiments.

##### Automated flow peptide synthesis

125 mg of H-RAM ChemMatrix® resin was weighed in a 6 mL Luer Torviq reactor and swollen using DCM (15 min) and DMF (15 min). The reactor was drained and placed in the heating reactor of the “Amidator,” an automated-flow peptide synthesizer built in the Pentelute Lab. Sequences for A0A7W1B5T5 and A0A1T4N4K0 were taken directly from UniProt. Synthesis was carried out as previously described [123, 124]. After synthesis, the resin was drained, washed with DCM ×3, and left to dry on the vacuum manifold.

##### Cleavage protocol

Half of the peptide-anchored resin was transferred to a 10 mL Luer-Lock Torviq syringe equipped with a frit and was cleaved for 4 hours at room temperature using 5 mL of a Met-Reducing cleavage cocktail containing 77.5% TFA, 5% thioanisole, 5% dimethylsulfide, 5% TIPS, 5% TMS-Cl, and 2.5% EDT. After 4 hours, the cleavage mixture was filtered and evaporated under N2 flow. The resulting solid was dissolved in 20 mL of 50% ACN in water with 0.1% formic acid, flash-frozen using liquid nitrogen, and lyophilized.

##### Preparative liquid chromatography-mass spectrometry (Prep LC-MS)

Preparative HPLC purification was performed on an Agilent Technologies 1260 Infinity LC mass-directed purification system coupled to a 6130 Single Quad MS. A Timberline Instrument TL105 HPLC column heater was used to heat the column to 50 *^◦^*C. The crude lyophilized peptide powder was solubilized in 6 mL of a denaturing buffer consisting of 6 M Gdm HCl and 100 mM TRIS HCl at pH 7.5.

The mixture was filtered using a 0.22 *µ*m nylon syringe filter. An analytical HPLC run was first performed using a low volume of protein to estimate %B, and a focused gradient was used for the purification runs as follows: Analytical run: Column: Agilent Zorbax 300 Stable Bond C3 PrepHT, 21.1 × 100 mm, 5 *µ*m. Flow Rate: 20 mL/min. Solvents: A: H_2_O + 0.1% TFA, B: ACN + 0.1% TFA. Column Temperature: 50 *^◦^*C. Gradient: 0-10 min 5% B, 10-40 min 5-95% B, 40-45 min 95% B.

Focused gradient run: To generate the final preparative HPLC method, we estimated the %B of elution during the analytical run and performed a preparative run using a linear gradient of −10% to +5%B in 60 minutes around the %B identified in the analytical run. Column: Agilent Zorbax 300 Stable Bond C3 PrepHT, 21.1 × 100 mm, 5 *µ*m. Flow Rate: 20 mL/min. Solvents: A: H_2_O + 0.1% TFA, B: ACN + 0.1% TFA. Column Temperature: 50 *^◦^*C. Gradient: see description.

Fractions were analyzed by LC-MS, chosen based on apparent purity, flash-frozen using liquid nitrogen and lyophilized.

##### Protein folding via dialysis

The purified lyophilized fractions were solubilized with a denaturing solution containing 6 M guanidinium chloride, HEPES 10 mM, pH = 7.4 and grouped, using a total volume of less than 6 mL. Solubilized protein was dialysed against a 20 mM TRIS, 200 mM NaCl, pH 7.5 buffer using a 3.5 kDa MWCO dialysis membrane (Slide-A-Lyzer® MINI Dialysis Devices, 2 mL, cat. no. 88403) from Thermo. Buffer changes were performed after 2 and 24 hours for a total of three dialysis cycles over 48 hours.

##### Liquid chromatography-mass spectrometry (LC-MS)

Mass analysis was performed on an Agilent Technologies 1290 Infinity II UHPLC coupled to a 6550 iFunnel Q-TOF LC-MS system. MS spectra were acquired in positive ionization mode with an m/z range of 300–2000 Da:

C4-1-91-10min: Phenomenex Aeris C4 column (2.1 x 150 mm, 3.6 *µ*m, 0.2 mL/min, 40 °C), 1% B for 0–1 min, 1–91% B over 1–7 min, 1% B for 7–9 min. (ESI+, mass range 100–1700 m/z). Column: Phenomenex Aeris WIDEPORE C4 200 Å (2.1 x 150 mm, 3.6 *µ*m, Cat. no. 00F-4486-AN.). Flow Rate: 0.3 mL/min. Solvents: A: H_2_O + 0.1% FA, B: ACN + 0.1% FA. Column Temperature: 40 *^◦^*C. Gradient: 0-2 min 1% B, 2-8 min 1%-91% B, 8-10 min 91%-95% B (MS acquisition from 2 to 8 min).

Data were processed using Agilent MassHunter BioConfirm software version 10.0. Mass deconvolution was carried out using the 600–2000 m/z mass range, and output deconvoluted masses were fixed between 10,000 and 20,000 Da.

##### Protein concentration

After three cycles of dialysis (48 hours total), the protein solution was concentrated using pre-humidified 3 kDa MWCO centrifugal filters (Amicon® Ultra – 4). Samples were deposited onto the filters and spun at 4,000 RPM for 99 minutes, or until the combined remaining solution volume was under 500 µL. The resulting solution was transferred to protein lo-bind Eppendorf tubes and assayed for protein concentration (A280) using a Tecan Spark® plate reader. Extinction coefficients were estimated using the ProtParam tool from the ExPASy Bioinformatics Resource Portal, based on the primary sequence of the protein candidates and specifying a C-terminal amide and reduced cysteines [125].

##### Analytical high-performance liquid chromatography (HPLC)

Analytical HPLC was carried out on an Agilent 1290 series system with UV detection at 214 nm. C3-5-65-60min: Zorbax 300-SB C3 column (2.1 × 150 mm, 3.6 *µ*m, 0.2 mL/min, 40 *^◦^*C), 5% B for 0–5 min, 5–65% B over 5-65 min, 65-95% B for 65-66 min, 95% B for 66-70 min, 95-5% B for 70-75 min. Column: Zorbax 300-StableBond C3 (2.1 × 150 mm, 5 *µ*m, cat. no. 883750-909.) Flow Rate: 1 mL/min. Solvents: A: H_2_O + 0.1% FA, B: ACN + 0.1% FA. Column Temperature: 40 *^◦^*C. Gradient: 0-5 min 5% B, 5-65 min 5%-65% B, 65-66 min 65%-95% B, 66-70 min 95% B, 70-75 min 95-5% B. UV 214 nm integrations were performed automatically by the built-in Agilent OpenLab CDS, ChemStation Edition Rev.C.01.10[287].

##### Protein–heme sample preparation

A stock solution of hemin in DMF (concentration variable, 33.3x the concentration of the protein in solution) was added to the protein in buffer in equimolar amounts (1 eq.) with a final concentration of 3% DMF. The solution was vortexed and incubated at room temperature for 5 minutes as previously described [126].

##### UV–vis spectroscopy assay of heme association

Nanodrop methodology (Synthetic candidates): 2 *µ*L of the reconstituted protein–heme solution was deposited into a nanodrop plate (NanoQuant plate), and its absorbance spectrum was recorded using a Tecan Spark® plate reader at wavelengths between 300 and 650 nm (1 nm step, 3.5 nm slit). All samples were corrected using a blank containing 3% DMF in buffer, and negative controls of hemin alone and hemin with bovine serum albumin (BSA) were included. A red shift in the Soret maximum (Soret *λ*_max_) relative to the free-heme and all negative controls was interpreted as spectral evidence consistent with heme association under the assay conditions [126, 127]. 384-well plate methodology (Recombinant candidates): Because the concentration of most recombinant candidates was lower than the detection threshold for nanodrop quantification, 120 µL of each reconstituted protein sample was added to a 384-well plate from Greiner Bio-One (MICROPLATE, 384 WELL, PS, *µ*CLEAR®, WHITE, NON-BINDING, cat. no. 781903). Absorbance spectra were recorded using a Tecan Spark® plate reader at wavelengths between 300 and 650 nm (1 nm step, 3.5 nm slit). All samples were corrected using a blank containing 3% DMF in buffer, and negative controls of hemin alone and hemin with BSA were included. In the case of V5BF69, the concentration of the protein was still insufficient to record UV-vis spectra and was therefore concentrated from 1.28 µM to 9.17 µM prior to reconstitution using Amicon Ultra - 0.5 mL, 10 kDa spin filters (cat. no. UFC501024) obtained from Merck Millipore.

