## Supplementary Information for "Learning Universal Representations of Intermolecular Interactions with ATOMICA"

This PDF file includes:

Supplementary Figures S1 to S11
Supplementary Tables S1 to S9
Supplementary Notes 1 and 2

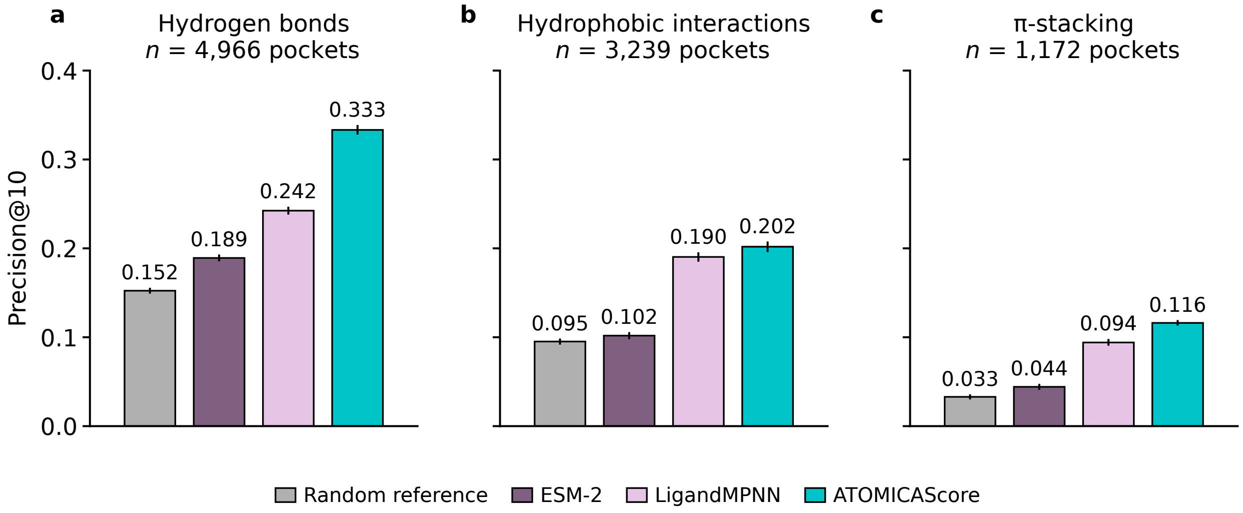


**Figure S1: Precision@10, the fraction of the 10 top-ranked residues that are involved in an annotated non-covalent interaction, for protein–small-molecule complexes in the pretraining test set.** We compare ATOMICASCORE, ESM-2 (3B), LigandMPNN, and a random reference on the recovery of the following types of annotated non-covalent interaction: **a** hydrogen bonds, **b** hydrophobic interactions, and **c** $\pi$-stacking.

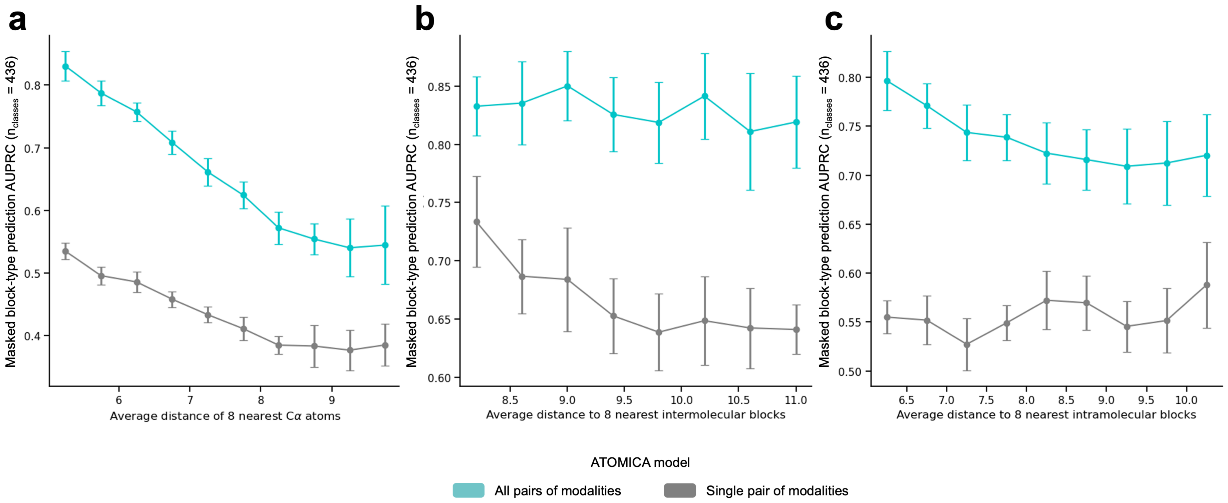


**Figure S2: Masked block-type prediction AUPRC of ATOMICA models trained on all pairs of interacting modalities compared to one pair of interacting modalities.** **a** Masked amino-acid prediction AUPRC binned by average distance to the eight nearest C$\alpha$ atoms. **b** Masked block-type prediction AUPRC binned by average distance to the eight nearest intermolecular blocks. **c** Masked block-type prediction AUPRC binned by average distance to the eight nearest intramolecular blocks. The evaluation protocol is described in Methods 8.

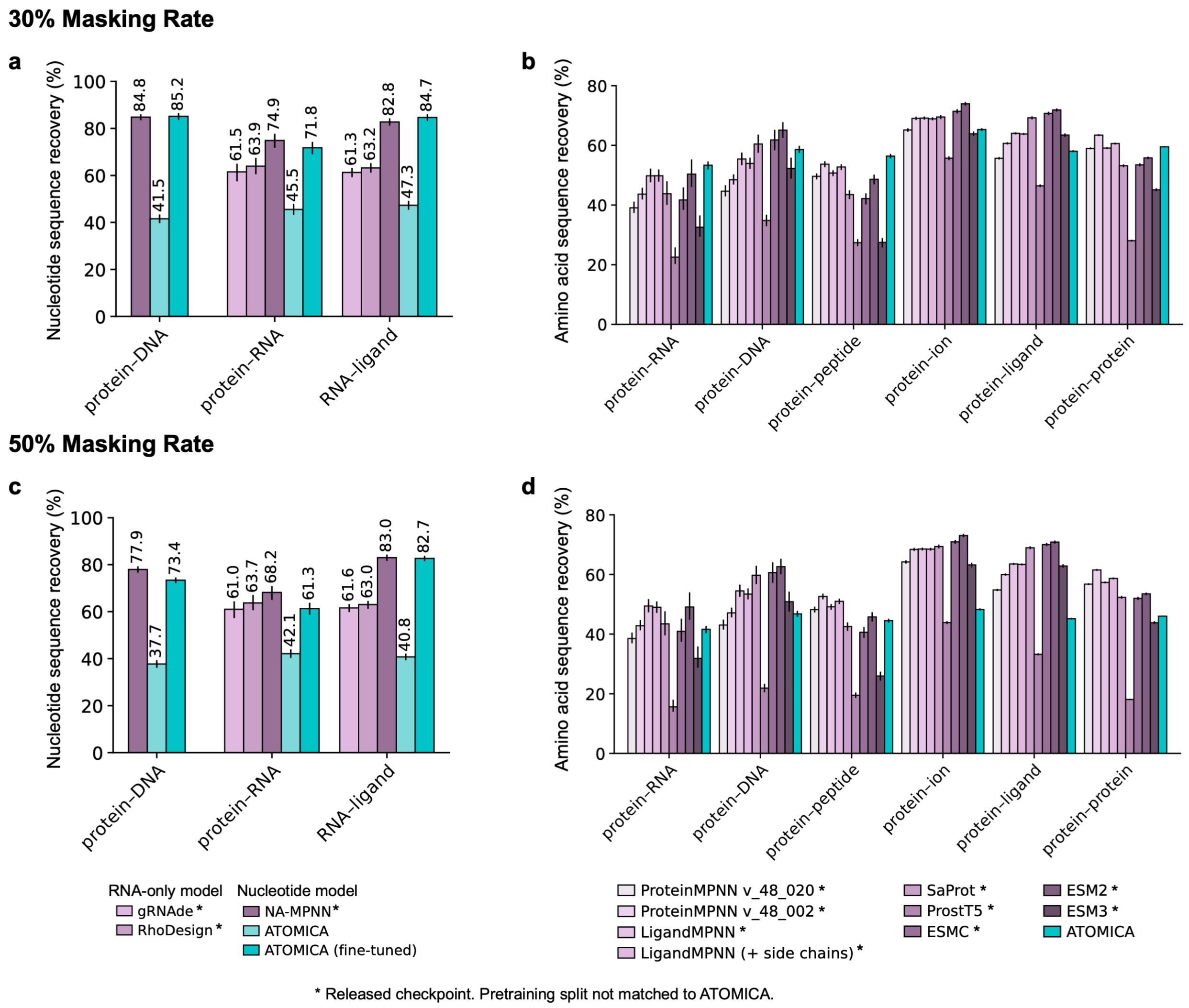


**Figure S3: Nucleotide and amino-acid sequence recovery.** Nucleotide sequence recovery at **a** 30% and **c** 50% masking of nucleotide residues at the interaction interface for protein–RNA, protein–DNA, and RNA–ligand complexes, reported for ATOMICA (fine-tuned). Amino-acid sequence recovery at **b** 30% and **d** 50% masking of amino-acid residues at the interaction interface across six protein-containing interaction modalities, reported for the pretrained ATOMICA model without task-specific fine-tuning. For the masking panels, error bars denote 95% bootstrap confidence intervals over complexes, with all models evaluated on identical masked positions within each modality. The evaluation protocol is described in Methods 8.

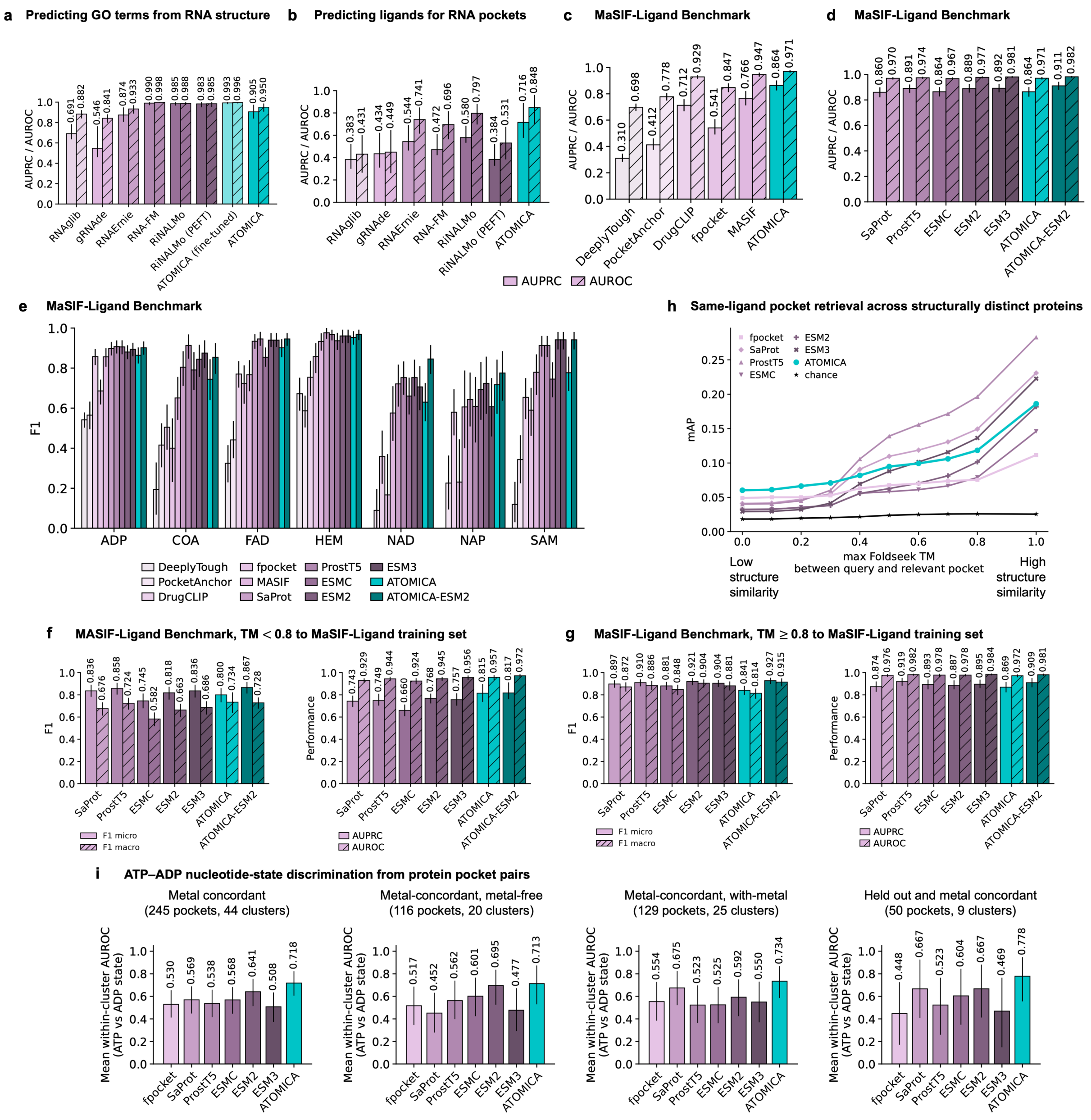


**Figure S4: Additional results for evaluation of ATOMICA on RNA and protein annotation benchmarks.** **a** AUPRC and AUROC for the prediction of GO terms for RNA. **b** AUPRC and AUROC for the prediction of ligands for RNA pockets. MaSIF-ligand AUPRC and AUROC for ATOMICA compared with **c** protein-pocket structure-based models and **d** protein language models. **e** Per-class F1 score on MaSIF-ligand for each of the seven ligand classes, comparing ATOMICA with the protein-pocket encoders and the protein language models. MaSIF-ligand performance stratified by Foldseek TM-score to the nearest training receptor, for **f** TM $< 0.8$ and **g** TM $\geq 0.8$. **h** Same-ligand pocket retrieval across structurally distinct proteins evaluated with mean average precision (mAP) at different thresholds of Foldseek TM cutoff for query and relevant pocket structure alignment. **i** ATP–ADP nucleotide-state discrimination from ligand-free protein pockets. Mean within-cluster AUROC for distinguishing ATP- and ADP-associated pocket states in the metal-concordant, metal-free, with-metal, and pretraining-held-out subsets. AUROC is calculated separately within each MMseqs2 sequence cluster and macro-averaged across clusters. Error bars denote 95% confidence intervals over pockets in **a–g** and over sequence clusters in **h** and **i**.

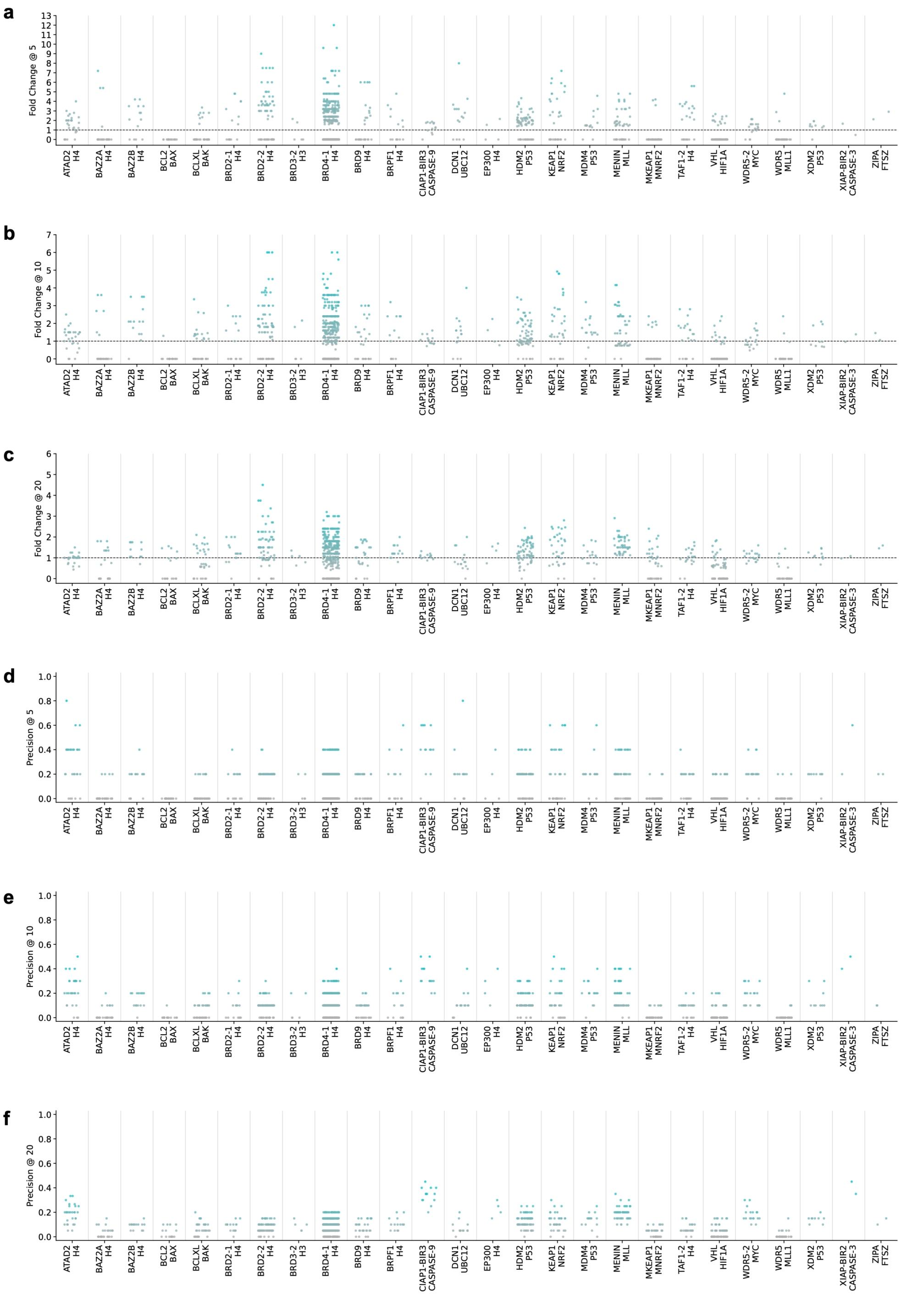


**Figure S5: Cross-modality interface comparison of orthosteric protein–peptide inhibitors.** Across protein–peptide systems, we measure the precision of the top-$k$ most similar peptide–inhibitor block pairs that are spatially proximal (aligned distance $\leq 4$ Å) and the corresponding fold change over the expected fraction among all block pairs. **a** Fold Change@5, **b** Fold Change@10, and **c** Fold Change@20. **d–f** Precision@5, Precision@10, and Precision@20.

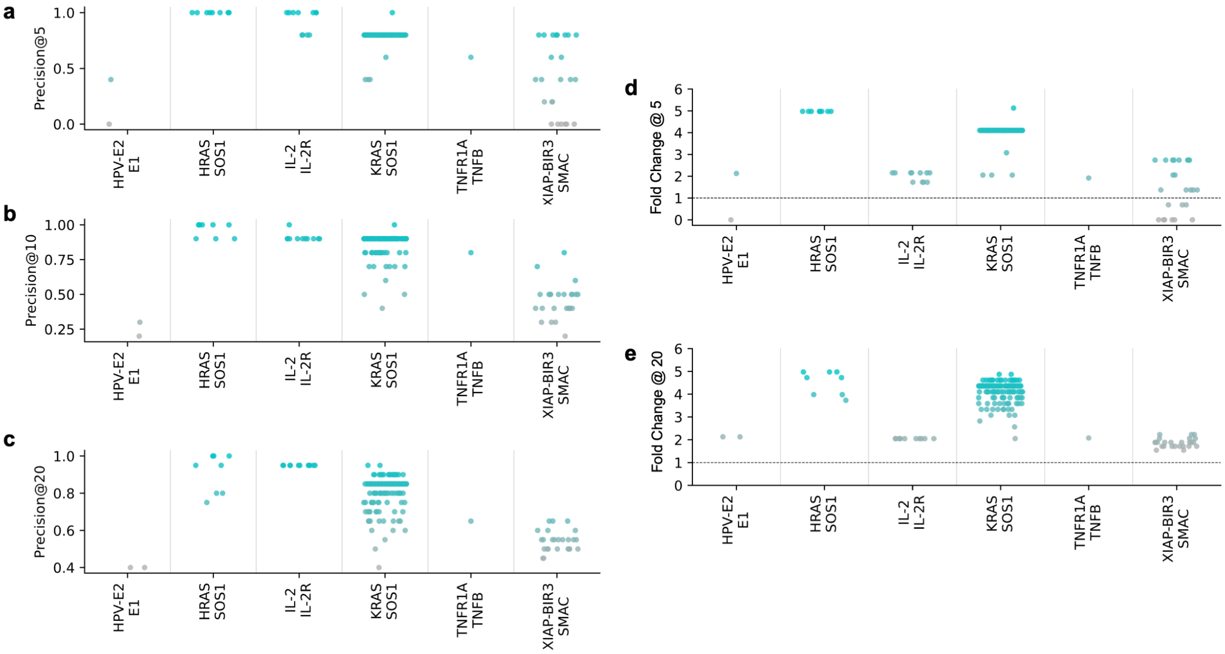


**Figure S6: Cross-modality interface comparison of orthosteric protein–protein inhibitors.** Given the PPI A–B and protein–inhibitor A–I, we define Fold Change@k for retrieval on protein B, defining positives as sampled interfaces whose centers lie within 12 Å of the native A–B interface on B (12 Å corresponds to the 25th percentile of sampled-interface distances to A); the baseline is the expected fraction over all sampled interfaces. **a** Precision@5, **b** Precision@10, **c** Precision@20, **d** Fold Change@5, **e** Fold Change@20.

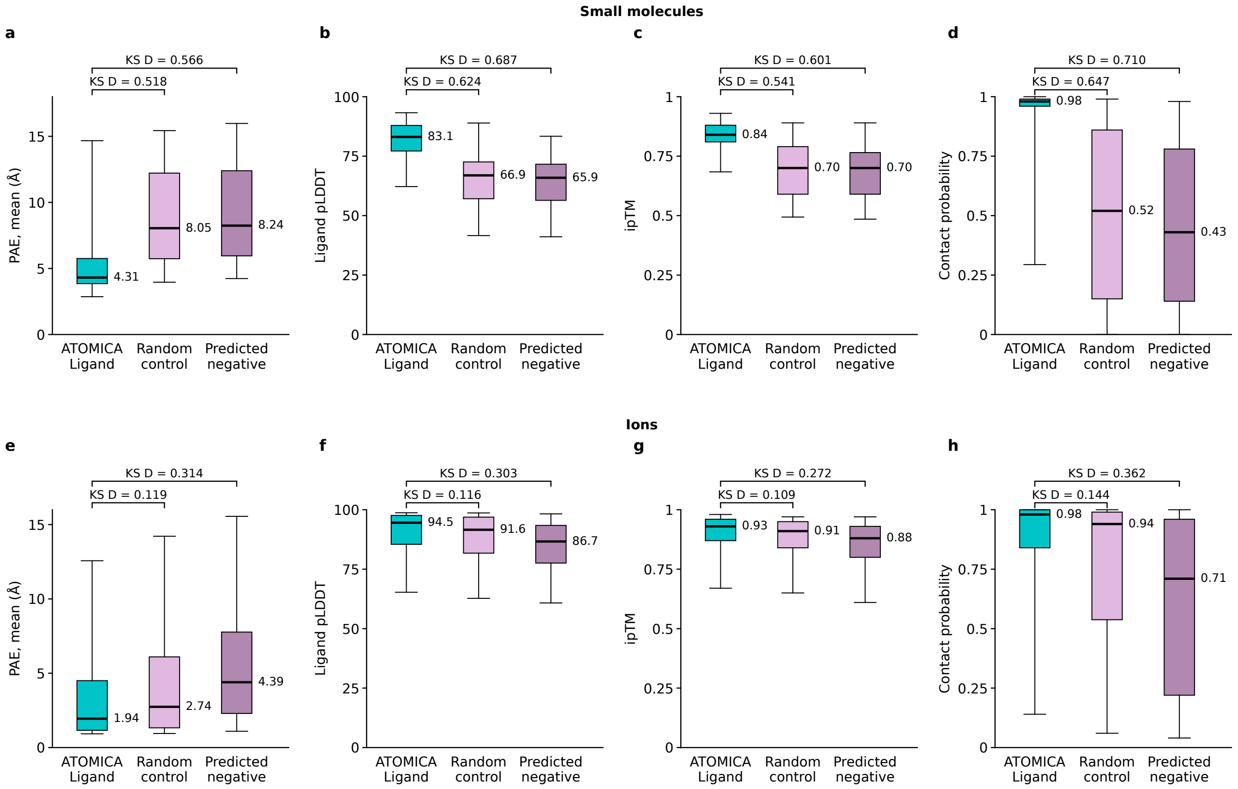


**Figure S7: Additional AlphaFold3 confidence measures for ATOMICA-Ligand dark proteome annotations.** The complexes ATOMICA-Ligand annotates are compared against a random control matched to them in the number and identity of the ligands, and against the subset of that control that ATOMICA-Ligand scores as negative (Methods 6). Boxes span the interquartile range, with the median marked and whiskers reaching the 5th and 95th percentiles. The number beside each box is its median, and brackets give the two-sample Kolmogorov–Smirnov statistic $D$ against each control. **a–d** Small-molecule ligands and **e–h** metal-ion ligands. We report the mean symmetrized predicted aligned error between the ligand and the pocket in **a** and **e**, the mean pLDDT over ligand atoms in **b** and **f**, the global ipTM in **c** and **g**, and the largest ligand-to-pocket contact probability in **d** and **h**.

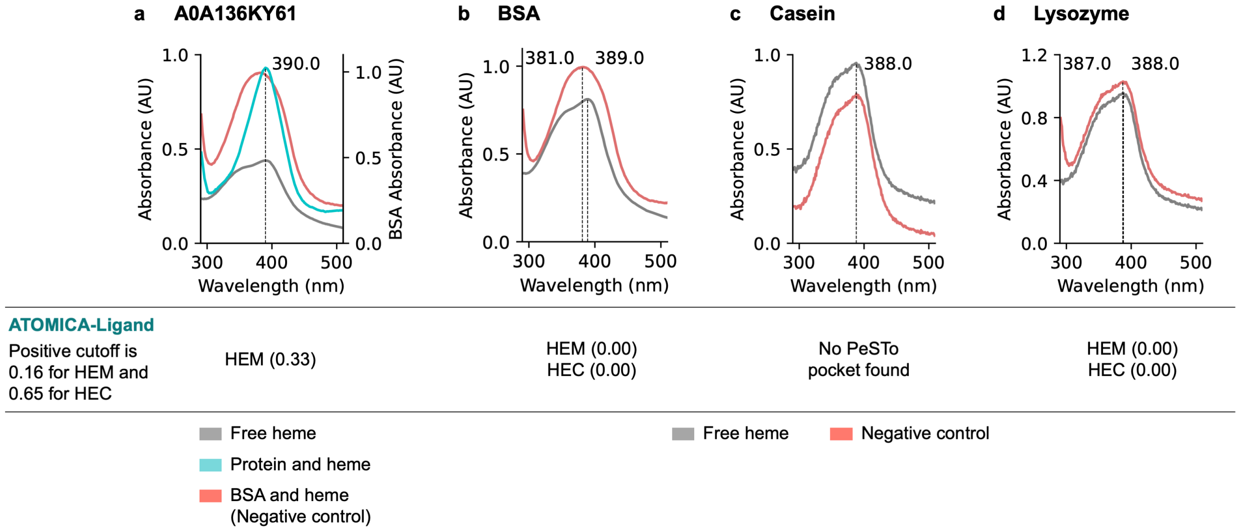


**Figure S8: UV–vis spectra for the heme candidate without a Soret shift and for the negative controls. a** A0A136KY61, an ATOMICA-Ligand HEM prediction that did not show a Soret-band red shift comparable to the candidates in Fig. 5. **b–d** Negative controls: **b** BSA, **c** casein, and **d** lysozyme. Soret-band $\lambda_{\max}$ values are indicated, and ATOMICA-Ligand predictions and their scores are shown beneath each panel.

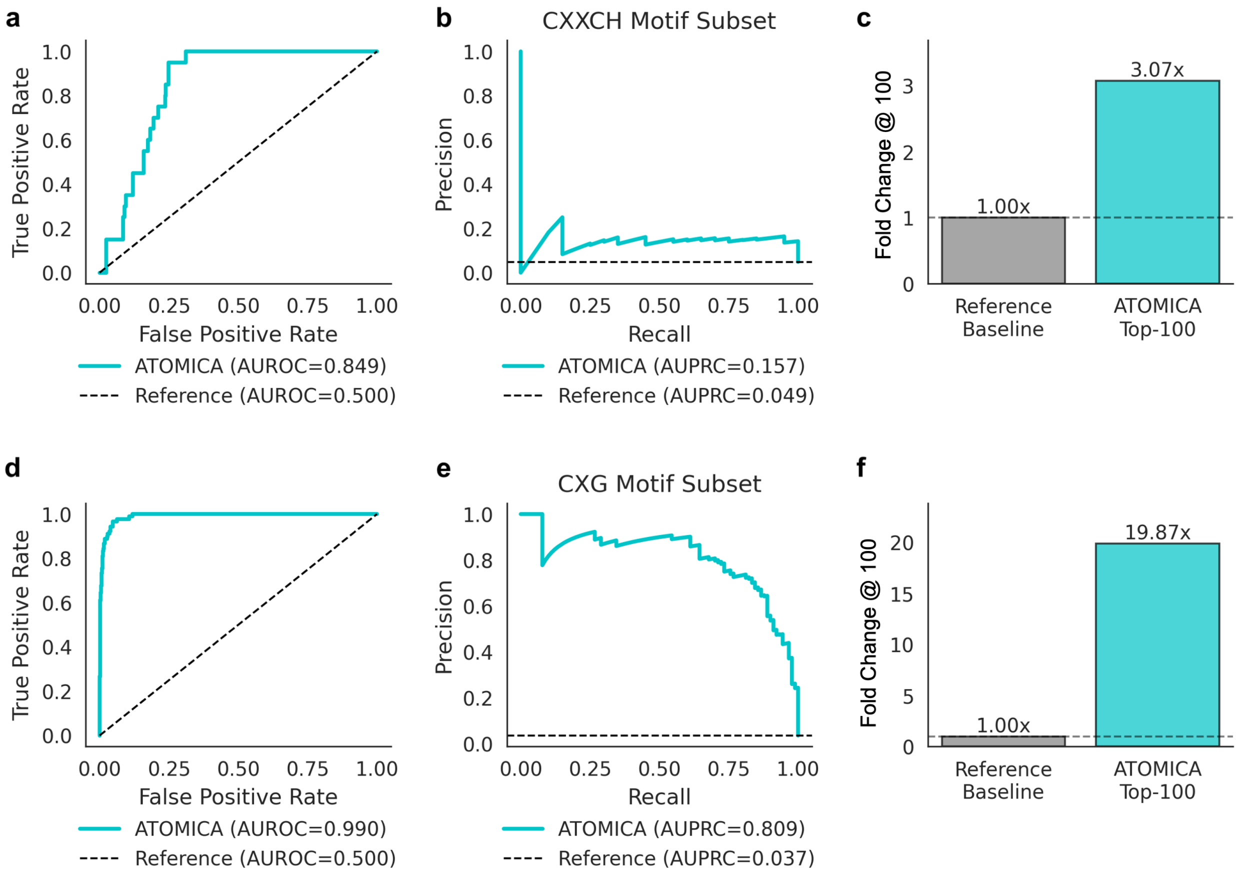


**Figure S9: Motif-controlled evaluation of ATOMICA-Ligand for heme-binding prediction. a–c** Performance on the subset of proteins in the ATOMICA-Ligand heme test set containing the canonical CXXCH motif: **a** ROC curve (AUROC), **b** precision–recall curve (AUPRC; dashed line indicates the subset prevalence baseline), and **c** Fold Change@100 of true heme binders among the top-100 ranked predictions relative to a reference baseline. **d–f** Analogous evaluation on the subset of proteins in the ATOMICA-Ligand heme test set containing the CXG motif: **d** ROC curve, **e** precision–recall curve, and **f** Fold Change@100 bar chart.

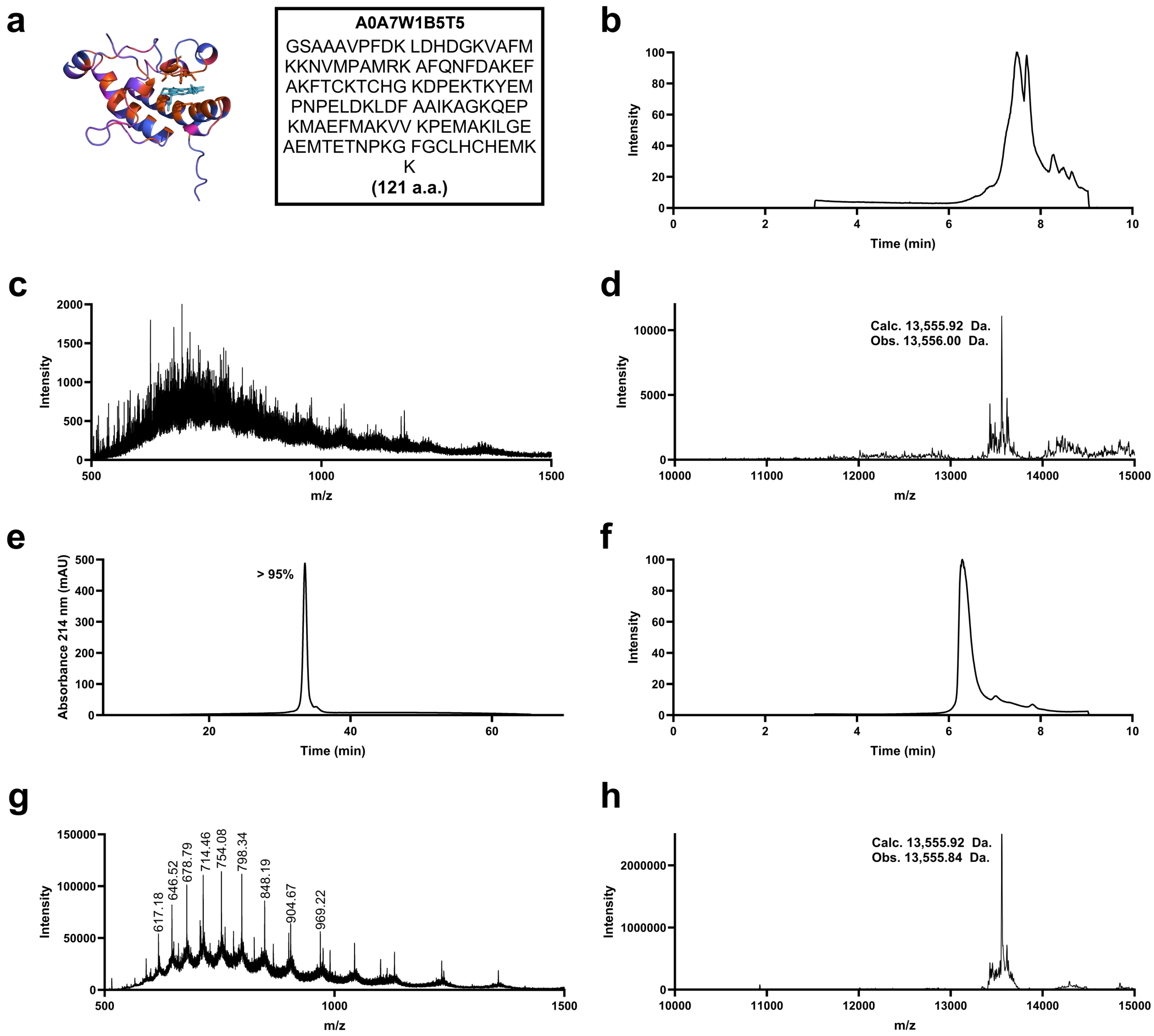


**Figure S10: Chemical characterization of AFPS-synthesized A0A7W1B5T5.** **a** AlphaFold2 predicted structure of A0A7W1B5T5 and its sequence. **b** Crude LC-MS spectra (Method: C4-1-91-10min). **c** Crude MS TIC scan (7.420-7.552 min, 13 scans). **d** Crude deconvoluted MS spectra. Calculated $[M+H]^+$ = 13,555.92 Da, Observed $[M+H]^+$ = 13,556.00 Da. **e** Purified UHPLC spectrum (214 nm, Method: C3-5-65-60min). >95% purity by UV integration. **f** Purified LC-MS spectra (Method: C4-1-91-10min). **g** Purified MS TIC scan (6.284-6.339 min, 6 scans). **h** Purified deconvoluted MS spectra of Candidate 1. Calculated $[M+H]^+$ = 13,555.92 Da, Observed $[M+H]^+$ = 13,555.84 Da.

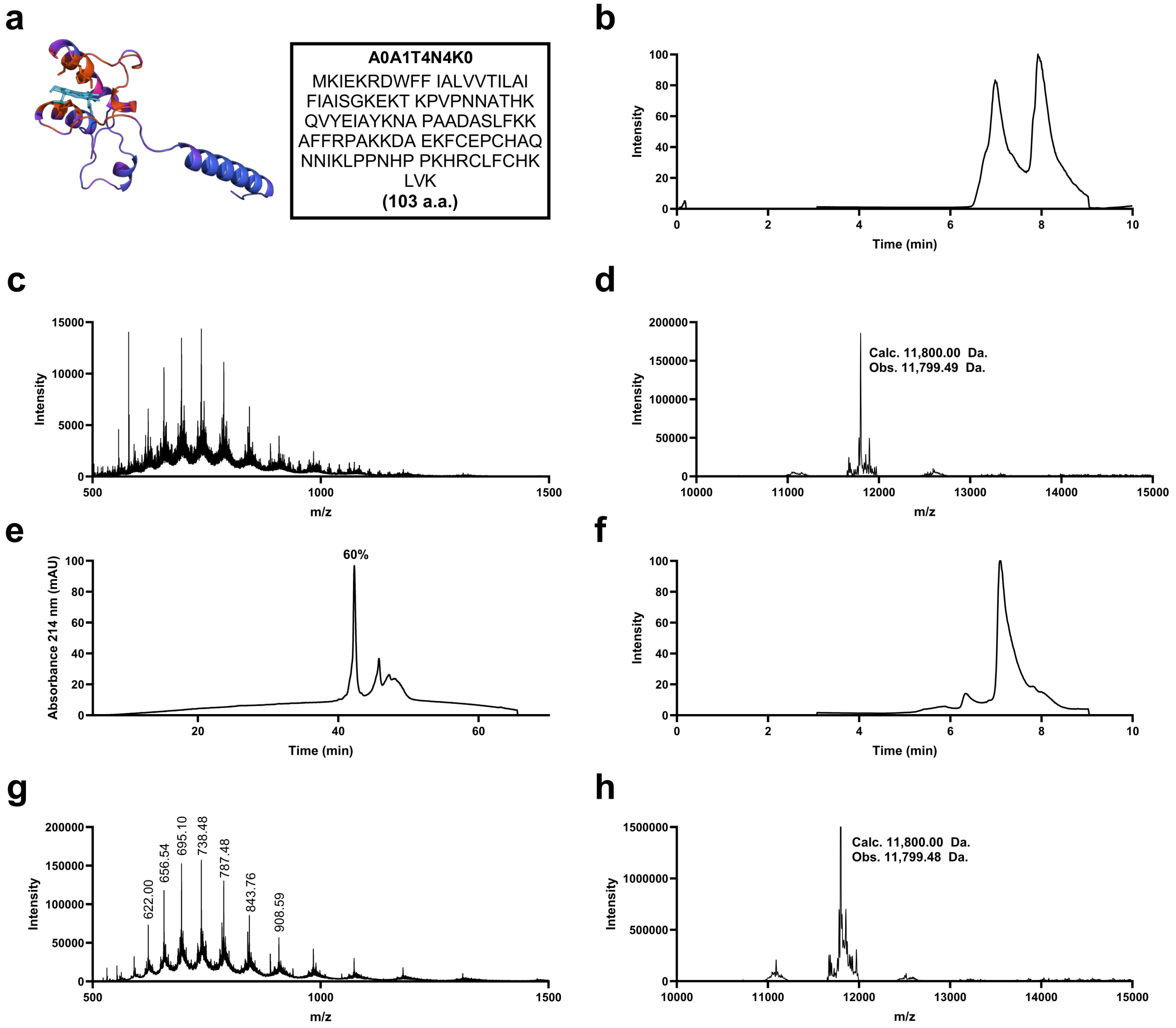


**Figure S11: Chemical characterization of AFPS-synthesized A0A1T4N4K0.** **a** AlphaFold2 predicted structure of A0A1T4N4K0 and its sequence. **b** Crude LC-MS spectra (Method: C4-1-91-10min). **c** Crude MS TIC scan (7.878-8.066 min, 18 scans). **d** Crude deconvoluted MS spectra. Calculated $[M+H]^+$ = 11,800.00 Da, Observed $[M+H]^+$ = 11,799.49 Da. **e** Purified UHPLC spectrum (214 nm, Method: C3-5-65-60min). 60% purity by UV integration. **f** Purified LC-MS spectra (Method: C4-1-91-10min). **g** Purified MS TIC scan (7.063-7.151 min, 9 scans). **h** Purified deconvoluted MS spectra of Candidate 2. Calculated $[M+H]^+$ = 11,800.00 Da, Observed $[M+H]^+$ = 11,799.48 Da.

**Table S1: Linear probes of metal coordination from $\mathbf{z}_b^{\text{block}}$.** Balanced accuracy is the primary metric for the imbalanced classes, with accuracy and macro-$F_1$ reported as secondary metrics. Brackets denote 95% percentile confidence intervals from bootstrap resampling of PDB entries. The all-donor target uses the complete deposited MetalPDB coordination sphere [1], whereas the protein-donor target includes only donors from protein residues visible in the model input. The fixed-six analysis contains the three observed FindGeo [2] classes, which are octahedron, pentagonal bipyramid with a vacancy, and other.

| Predictor | Balanced accuracy | Accuracy | Macro-$F_1$ |
|---|---|---|---|
| *Coordination number over all deposited donors* (8 classes, $n = 3{,}654$ sites from 1,562 PDB entries) | | | |
| Majority class | 0.125 [0.125, 0.125] | 0.272 [0.239, 0.311] | 0.054 [0.048, 0.059] |
| Element only | 0.275 [0.261, 0.290] | 0.387 [0.348, 0.432] | 0.183 [0.168, 0.200] |
| ATOMICA (frozen) | 0.368 [0.333, 0.410] | 0.408 [0.375, 0.446] | 0.346 [0.312, 0.389] |
| *Coordination number over visible protein donors* (8 classes, $n = 3{,}654$ sites from 1,562 PDB entries) | | | |
| Majority class | 0.125 [0.125, 0.125] | 0.282 [0.247, 0.317] | 0.055 [0.050, 0.060] |
| Element only | 0.248 [0.225, 0.269] | 0.335 [0.301, 0.373] | 0.167 [0.151, 0.184] |
| ATOMICA (frozen) | 0.578 [0.527, 0.642] | 0.648 [0.606, 0.690] | 0.554 [0.508, 0.610] |
| *FindGeo coordination geometry* (14 classes, $n = 2{,}313$ sites from 1,162 PDB entries) | | | |
| Majority class | 0.071 [0.071, 0.071] | 0.212 [0.182, 0.243] | 0.025 [0.022, 0.028] |
| Element only | 0.171 [0.161, 0.180] | 0.399 [0.362, 0.439] | 0.108 [0.100, 0.116] |
| ATOMICA (frozen) | 0.340 [0.310, 0.376] | 0.496 [0.456, 0.537] | 0.334 [0.300, 0.365] |
| *FindGeo geometry at deposited coordination number six* (3 classes, $n = 635$ sites from 338 PDB entries) | | | |
| Majority class | 0.333 [0.333, 0.333] | 0.677 [0.608, 0.742] | 0.269 [0.252, 0.284] |
| Element only | 0.333 [0.333, 0.333] | 0.677 [0.608, 0.742] | 0.269 [0.252, 0.284] |
| ATOMICA (frozen) | 0.499 [0.410, 0.584] | 0.668 [0.588, 0.737] | 0.500 [0.412, 0.586] |

**Table S2: Same-ligand pocket retrieval across structurally distinct proteins.** Retrieval statistics are reported for 892 pockets, 428 queries, and 44 ligand classes defined in Methods 4. Values are macro-averaged over 30%-sequence-identity clusters. Lift is macro mAP relative to the chance value in the first row, and brackets denote 95% bootstrap confidence intervals over the 30%-sequence-identity clusters.

| Model | mAP (lift) | nDCG | R-prec |
|---|---|---|---|
| Random reference | 0.0184 | 0.256 | 0.0123 |
| ProstT5 | 0.0401 [0.0297, 0.0541] (2.18×) | 0.273 [0.256, 0.292] | 0.0289 [0.0172, 0.0440] |
| ESM-2 (3B) | 0.0325 [0.0254, 0.0414] (1.77×) | 0.267 [0.251, 0.284] | 0.0240 [0.0157, 0.0342] |
| ESM-C (600M) | 0.0318 [0.0272, 0.0368] (1.73×) | 0.271 [0.256, 0.288] | 0.0248 [0.0178, 0.0320] |
| SaProt (1.3B) | 0.0411 [0.0334, 0.0498] (2.24×) | 0.281 [0.264, 0.299] | 0.0323 [0.0227, 0.0435] |
| ESM-3 | 0.0293 [0.0217, 0.0410] (1.59×) | 0.261 [0.246, 0.276] | 0.0198 [0.0111, 0.0317] |
| amino-acid composition | 0.0380 [0.0284, 0.0512] (2.07×) | 0.274 [0.256, 0.292] | 0.0306 [0.0198, 0.0440] |
| fpocket | 0.0491 [0.0359, 0.0653] (2.67×) | 0.286 [0.266, 0.307] | 0.0486 [0.0344, 0.0665] |
| ATOMICA (frozen) | **0.0603** [0.0443, 0.0781] (3.28×) | **0.301** [0.279, 0.323] | **0.0586** [0.0417, 0.0784] |

**Table S3: Matched inhibitor structures for protein–peptide complexes.** Counts are reported for the cross-modality comparison of orthosteric protein–peptide inhibitors. The dataset contains 26 complexes and 965 matched inhibitor structures.

| Protein–Peptide Complex | # Protein–Inhibitor Structures |
|---|---|
| BRD4-1/H4 | 338 |
| HDM2/P53 | 70 |
| BRD2-2/H4 | 67 |
| VHL/HIF1A | 64 |
| MENIN/MLL | 49 |
| MKEAP1/MNRF2 | 35 |
| KEAP1/NRF2 | 33 |
| BCLXL/BAK | 32 |
| WDR5/MLL1 | 30 |
| ATAD2/H4 | 29 |
| BRD9/H4 | 28 |
| BAZ2A/H4 | 25 |
| TAF1-2/H4 | 22 |
| WDR5-2/MYC | 19 |
| BCL2/BAX | 16 |
| BRD2-1/H4 | 15 |
| DCN1/UBC12 | 15 |
| MDM4/P53 | 15 |
| BAZ2B/H4 | 14 |
| BRPF1/H4 | 14 |
| CIAP1-BIR3/CASPASE-9 | 13 |
| XDM2/P53 | 9 |
| BRD3-2/H3 | 5 |
| EP300/H4 | 4 |
| XIAP-BIR2/CASPASE-3 | 2 |
| ZIPA/FTSZ | 2 |

**Table S4: Matched inhibitor structures for protein–protein complexes.** Counts are reported for the cross-modality comparison of orthosteric protein–protein inhibitors.

| Protein–Protein Complex | # Protein–Inhibitor Structures |
|---|---|
| KRAS/SOS1 | 140 |
| XIAP-BIR3/SMAC | 25 |
| IL-2/IL-2R | 11 |
| HRAS/SOS1 | 8 |
| HPV-E2/E1 | 2 |
| TNFR1A/TNFB | 1 |

**Table S5: Block-level embedding and geometric-distance correlations.** Spearman correlations compare ATOMICA block embedding distance, $\mathrm{dist}_{\mathrm{ATOMICA}}(\mathbf{h}^{\mathrm{block}}_{\mathrm{inhibitor}}, \mathbf{h}^{\mathrm{block}}_{\mathrm{peptide}})$, with aligned 3D distance, $\mathrm{dist}_{\mathbb{R}^3}(\mathbf{x}^{\mathrm{block}}_{\mathrm{inhibitor}}, \mathbf{x}^{\mathrm{block}}_{\mathrm{peptide}})$. Each coefficient pools block pairs from all inhibitor structures matched to one protein–peptide complex.

| Protein–Peptide Complex | Matched structures | Block pairs | Spearman $\rho$ |
|---|---|---|---|
| ATAD2/H4 | 29 | 680 | 0.051 |
| BAZ2A/H4 | 25 | 1,089 | -0.227 |
| BAZ2B/H4 | 14 | 392 | 0.005 |
| BCL2/BAX | 16 | 3,692 | 0.086 |
| BCLXL/BAK | 32 | 5,439 | -0.083 |
| BRD2-1/H4 | 15 | 592 | -0.044 |
| BRD2-2/H4 | 67 | 5,190 | 0.037 |
| BRD3-2/H3 | 5 | 288 | 0.170 |
| BRD4-1/H4 | 338 | 22,416 | -0.075 |
| BRD9/H4 | 28 | 1,530 | -0.074 |
| BRPF1/H4 | 14 | 464 | -0.067 |
| CIAP1-BIR3/CASPASE-9 | 13 | 440 | 0.000 |
| DCN1/UBC12 | 15 | 1,440 | 0.177 |
| EP300/H4 | 4 | 207 | 0.139 |
| HDM2/P53 | 70 | 6,539 | 0.057 |
| KEAP1/NRF2 | 33 | 3,568 | 0.358 |
| MDM4/P53 | 15 | 1,356 | 0.033 |
| MENIN/MLL | 49 | 3,732 | 0.321 |
| MKEAP1/MNRF2 | 35 | 4,752 | 0.139 |
| TAF1-2/H4 | 22 | 959 | 0.183 |
| VHL/HIF1A | 64 | 6,855 | 0.074 |
| WDR5-2/MYC | 19 | 624 | -0.013 |
| WDR5/MLL1 | 30 | 2,688 | -0.133 |
| XDM2/P53 | 9 | 682 | -0.102 |
| XIAP-BIR2/CASPASE-3 | 2 | 60 | 0.057 |
| ZIPA/FTSZ | 2 | 187 | 0.232 |

**Table S6: Interface-level embedding and geometric-distance correlations.** Spearman correlations compare ATOMICA interface embedding distance, $\text{dist}_{\text{ATOMICA}}(\mathbf{h}^{\text{interface}}_{\text{inhibitor}}, \mathbf{h}^{\text{interface}}_{\text{protein B}})$, with distance to the native A–B interface. Each coefficient pools sampled interfaces from all inhibitor structures matched to one protein–protein complex.

| Protein–Protein Complex | Matched structures | Sampled interfaces | Spearman $\rho$ |
|---|---|---|---|
| HPV-E2/E1 | 2 | 2,000 | 0.425 |
| HRAS/SOS1 | 8 | 8,000 | 0.078 |
| IL-2/IL-2R | 11 | 11,000 | 0.447 |
| KRAS/SOS1 | 140 | 140,000 | 0.072 |
| TNFR1A/TNFB | 1 | 1,000 | -0.148 |
| XIAP-BIR3/SMAC | 25 | 24,400 | 0.342 |

**Table S7: Interface aggregation ablation for protein–protein retrieval.** Fold Change@10 is reported for the three ATOMICA interface representations defined in Methods 2, with each of five 2P2Idb superfamilies contributing the unweighted mean over its complexes.

| Interface representation | Superfamilies with FC@10 > 1 | Mean FC@10 |
|---|---|---|
| Mean over $\mathbf{h}^{\text{block}}_b$ | 1/5 | 0.791 |
| $\mathbf{z}^{\text{interface}}$ | 2/5 | 0.925 |
| $\mathbf{h}^{\text{interface}}$ | **5/5** | **2.288** |

**Table S8: Experimental heme candidates selected using ATOMICA-Ligand.** Concentration and apparent purity are reported for candidates produced by AFPS or recombinant methods and for the negative controls.

| **Protein** | **Production method** | **Calc. M.W. (Da)** | **Conc. ($\mu$M)** | **Apparent purity (%)** |
|---|---|---|---|---|
| A0A7W1B5T5 | AFPS | 13 556 | 238.50 | $> 95^{\#}$ |
| A0A1T4N4K0 | AFPS | 11 800 | 117.00 | $60^{\#}$ |
| A0A7W1B5T5 | Recombinant | 14 728 | 26.48 | $\geq 90^{*}$ |
| A0A1T4N4K0 | Recombinant | 12 841 | 4.67 | n.d.$^{\dagger}$ |
| A0A4P5TA35 | Recombinant | 18 043 | 5.54 | n.d.$^{\dagger}$ |
| A0A7W0X6V6 | Recombinant | 9 115 | 27.43 | $\geq 70^{*}$ |
| A0A2V6P8N7 | Recombinant | 8 576 | 11.66 | n.d.$^{\dagger}$ |
| V5BF69 | Recombinant | 70 349 | 1.28 | $\geq 80^{*}$ |
| A0A136KY61 | Recombinant | 24 261 | 13.19 | $\geq 95^{*}$ |
| A0A7Y8LED7 | Recombinant | 58 116 | 1.38 | $\geq 75^{*}$ |
| A0A7V7N0X5 | Recombinant | 51 703 | 1.16 | n.d.$^{\dagger}$ |
| BSA control | Commercial | 66 430 | 22.46 | $> 98$ |
| Casein control | Commercial | 2 062 | 15.00 | n.d. |
| Lysozyme control | Commercial | 14 300 | 15.00 | $> 90$ |

The AFPS products comprise the UniProt sequences with a C-terminal amide, whereas the recombinant products contain a Strep tag. Molar concentrations were calculated using the molecular weight of the corresponding construct.
$*$: Assayed by GenScript using SDS-PAGE. #: Assayed using $A_{214\text{ nm}}$ by UHPLC.
†: Protein quantity too low to assay purity via SDS-PAGE. n.d. = not determined.

**Table S9: Agreement between ATOMICA and DeepPBS residue importance.** ATOMICA importance is the negative of ATOMICASCORE, so larger values denote a larger representational change. Statistics are computed within each of the 159 complexes with at least eight jointly scored residues and averaged across complexes. Conditioned correlations and model-free rankers are defined in Supplementary Note 2, and brackets denote 95% percentile bootstrap confidence intervals over complexes.

| | ATOMICA importance | Distance to DNA | Contact count |
|---|---|---|---|
| Spearman $\rho$ with DeepPBS | $+0.502$ [$+0.469$, $+0.535$] | $+0.135$ [$+0.103$, $+0.165$] | $+0.155$ [$+0.121$, $+0.190$] |
| conditioned on burial | $+0.482$ [$+0.447$, $+0.518$] | — | — |
| conditioned on contact chemistry | $+0.167$ [$+0.130$, $+0.203$] | — | — |
| Complexes with $\rho > 0$ | **97%** | 79% | 78% |
| FDR-significantly positive | **79%** | 4% | 6% |
| Precision@10 | **0.599 [0.565, 0.634]** | 0.441 [0.404, 0.479] | 0.406 [0.362, 0.453] |
| AUROC | **0.776 [0.754, 0.798]** | 0.576 [0.552, 0.600] | 0.567 [0.544, 0.589] |

### S1 Ligand identity as a hypothesis for the functional annotation of dark clusters

To ground functional hypotheses in dark clusters, we use ATOMICA-Ligand–predicted ligand identities at conserved pockets to interpret potential biochemical roles. For the common metal ions $Mg^{2+}$ and $Zn^{2+}$, whose coordination geometries are well characterized, we show that ATOMICA-Ligand recovers canonical structural signatures. We also compare against AlphaFill [3], which transfers ligands from structural templates onto AlphaFold models. In these dark-cluster representatives, where close annotated structural templates are often unavailable, AlphaFill rarely yields ligand placements at the nominated pockets, motivating a complementary approach that nominates ligand identity directly from pocket geometry and chemistry.

ATOMICA-Ligand predicts $Mg^{2+}$ (ATOMICA-Ligand score 0.76) at an ion-binding site in A0A0M0BG38 (Fig. S12a). This protein represents a cluster of 45 members whose lowest common ancestor is assigned only to cellular organisms. The sequence-based function prediction method ProtNLM [4] predicts that this is a phosphatidate cytidylyltransferase (score 0.82). These enzymes are $Mg^{2+}$-dependent and use the bound metal to coordinate phosphate groups, which is consistent with the ATOMICA-Ligand annotation of an $Mg^{2+}$-binding site.

For two bacterial dark protein clusters, ATOMICA-Ligand assigns $Zn^{2+}$ (ATOMICA-Ligand score 0.93 for both A0A6C0E0P4 and A0A7Y4VG56; Fig. S12b,c) to binding motifs that resemble those of metallopeptidases. ATOMICA-Ligand identifies $Zn^{2+}$-binding sites with the HEXXH motif, which is present in most metallopeptidases, specifically in zincins [5,6], showing that ATOMICA-Ligand recovers established binding-site motifs. Structural alignment of the proteins against thermolysin (PDB: 2TLX) superposes the $Zn^{2+}$-binding motifs and gives TM-align scores of 0.49 and 0.44 for A0A6C0E0P4 and A0A7Y4VG56, respectively.

ATOMICA-Ligand further characterizes a C4 zinc finger motif belonging to a protein in a cluster of previously uncharacterized bacterial proteins. Zinc-finger domains have been studied extensively in eukaryotes, whereas prokaryotic zinc-finger proteins remain comparatively uncharacterized [7]. The ion-binding site in W8VVQ2 is predicted to bind $Zn^{2+}$ (ATOMICA-Ligand score 0.98) and shares its binding motif with 61 of the 66 members of the cluster. ProtNLM predictions suggest HNH endonuclease function (score 0.47). HNH endonucleases are metal-dependent nucleases with a $\beta\beta\alpha$-metal topology [8,9], which agrees with the structure of W8VVQ2 (Fig. S12d).

Next, ATOMICA-Ligand nominates heme as the most likely cofactor for ligand-binding pockets in two ancient clusters of predicted transmembrane proteins. The representative protein

A0A6B2YTG2 of a cluster of 134 proteins is assigned heme (ATOMICA-Ligand score 0.76) (Fig. S12e). A structurally similar cluster, A0A0K3BHV0 (TM-align 0.911), whose 101 members are mostly uncharacterized, also receives an ATOMICA-Ligand heme assignment with a score of 0.33. Sequence-based function prediction with ProtNLM provides limited additional specificity. The heme assignment is further supported by A0A7X0NTZ6, a protein in the A0A0K3BHV0 cluster that is annotated as a succinate dehydrogenase/fumarate reductase cytochrome b subunit, and by the structurally similar protein A0A2T0LHX9 (TM-align to A0A6B2YTG2 0.715), which is annotated as cytochrome c/quinol oxidase subunit I.

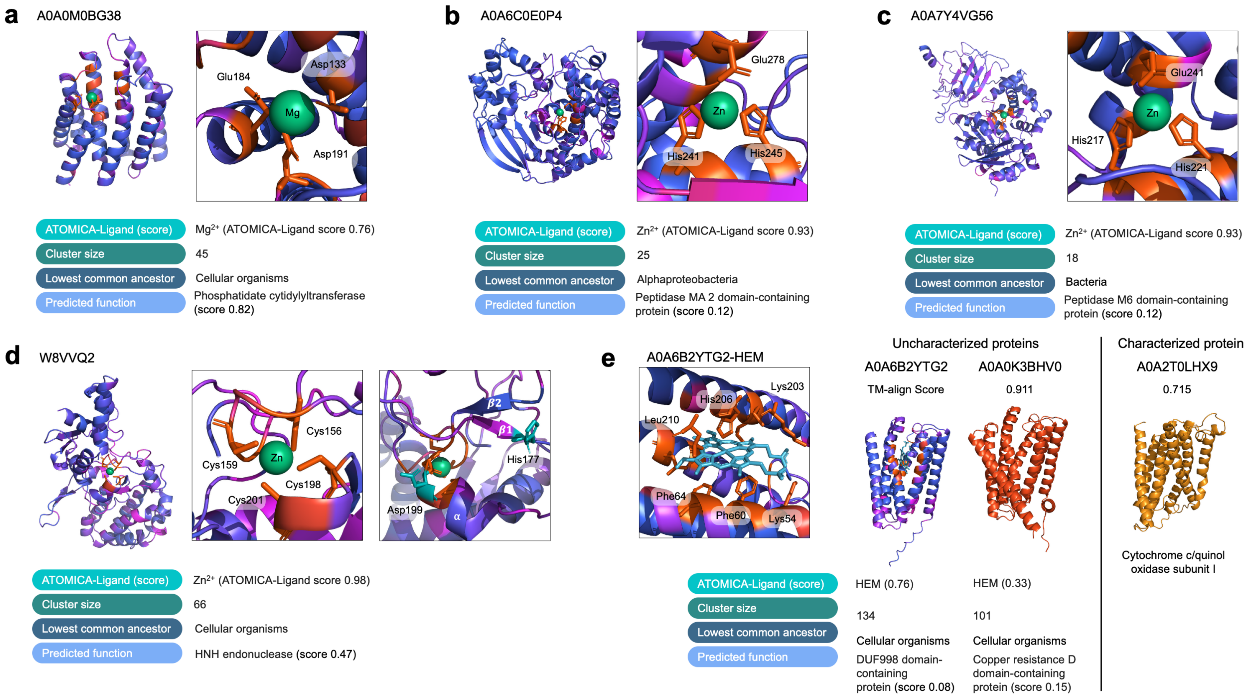


**Figure S12: Examples of dark proteome annotation with ATOMICA. a** $Mg^{2+}$ annotation for A0A0M0BG38, a putative phosphatidate cytidylyltransferase, where $Mg^{2+}$ binding is required for enzymatic activity. **b** A0A6C0E0P4 and **c** A0A7Y4VG56 are annotated with $Zn^{2+}$ and predicted to be metallopeptidases. Both contain the conserved HEXXH $Zn^{2+}$-binding motif characteristic of this family. **d** $Zn^{2+}$ annotation for W8VVQ2, with the predicted binding site indicating a C4 zinc finger domain, a motif that remains undercharacterized in Bacteria. **e** Heme annotation for A0A6B2YTG2, with comparison to neighboring clusters A0A0K3BHV0 and A0A2T0LHX9 suggesting these uncharacterized clusters may contain putative cytochrome proteins.

### S2 Comparison of AtomicaScore with DeepPBS residue importance

DeepPBS predicts DNA-binding specificity from a protein–DNA complex and assigns importance to protein heavy atoms by measuring how their perturbation changes the predicted position weight matrix (PWM) [10]. We compared these importance scores with AtomicaScore on the same protein–DNA structures. Both are leave-one-out perturbation measures. AtomicaScore measures the change in an unsupervised interaction representation, whereas DeepPBS measures the change in a supervised prediction of DNA specificity. ATOMICA was not trained with a PWM or DNA-binding-specificity objective.

**Dataset and score construction.** We used the 249 interaction graphs (188 PDB entries) in the 30% sequence identity protein–DNA test split, which was held out from ATOMICA pretraining. For protein residue $i$, AtomicaScore is the cosine similarity between $\mathbf{z}_G^{\text{ligand}}$ for the intact interaction graph and $\mathbf{z}_{G\setminus i}^{\text{ligand}}$ for the graph in which residue $i$ is replaced by the mask block, as defined in Methods 7. Because a lower cosine similarity denotes a larger perturbation, we define ATOMICA importance as the negative of AtomicaScore for the rank comparisons below. To obtain scores for DeepPBS, for every protein heavy atom within 5 Å of the DeepPBS DNA point cloud, we removed that atom's protein–DNA edges, recomputed the PWM, and used the mean absolute deviation from the unperturbed PWM as its importance. DeepPBS produces atom-level scores, so we averaged the importance of the interface atoms belonging to each residue.

**Statistical analysis and controls.** We calculated Spearman rank correlation between ATOMICA and DeepPBS importance separately in each complex and then averaged across complexes. A within-complex permutation test with 10,000 permutations tested each coefficient, and Benjamini–Hochberg correction was applied across complexes. Confidence intervals are 95% percentile intervals from 10,000 bootstrap resamples of complexes; residues from the same complex were never resampled independently. For retrieval, the ten residues with highest DeepPBS importance were defined as positives and residues were ranked by ATOMICA importance. Precision@10 is the fraction of ATOMICA's ten highest-ranked residues that are also in the DeepPBS top ten, and enrichment@10 divides this precision by the within-complex positive fraction.

We evaluated two model-free scorers on the same residues: negative minimum heavy-atom distance to DNA and the number of protein heavy atoms within 4.5 Å of DNA. We also calculated partial Spearman correlations. The burial-conditioned analysis controls for both distance and contact count. A stricter contact-chemistry analysis controls for the numbers of contacts to DNA base and backbone atoms and the distance to the nearest base atom, thereby separating direct contact with the base faces from contact with the phosphate–sugar backbone.

**Agreement with DeepPBS.** Using $\mathbf{z}_G^{\text{ligand}}$ as specified in the revised definition of ATOMICASCORE, ATOMICA importance recovered the DeepPBS residue ranking with a mean per-complex Spearman $\rho = +0.502$ $[+0.469, +0.535]$ (Supplementary Table S9). Correlations were positive in 97% of complexes and significantly positive after false-discovery-rate correction in 79%. Precision@10 was 0.599 [0.565, 0.634], corresponding to an enrichment@10 of 1.857 [1.743, 1.976]. Both agreement and retrieval exceeded the two model-free burial rankers.

The agreement was not explained solely by locating buried interface residues: conditioning on distance to DNA and contact count left $\rho = +0.482$ $[+0.447, +0.518]$. Interface geometry nevertheless accounts for an important part of the raw association. A model-free base-contact fraction reproduced the DeepPBS base-importance score at $\rho = +0.886$ $[+0.858, +0.909]$. After controlling for base contacts, backbone contacts, and distance to the nearest base, the ATOMICA–DeepPBS association was reduced but remained positive at $\rho = +0.167$ $[+0.130, +0.203]$. Under the same control, ATOMICA importance was positively associated with the DeepPBS base-importance component ($+0.144$ $[+0.110, +0.178]$) and negatively associated with its shape-importance component ($-0.118$ $[-0.157, -0.077]$). Twenty analyzed complexes were represented in DeepPBS's cross-validation folds; the mean correlation was $+0.536$ $[+0.473, +0.597]$ for these complexes and $+0.497$ $[+0.460, +0.534]$ for the 139 complexes held out from those folds, showing that the agreement was present in both strata. Together, these results show that the association between unsupervised ATOMICASCORE and DeepPBS's supervised specificity importance on double-stranded protein–DNA complexes is not explained by the measured burial and base-versus-backbone contact features.
